# Analgesia through FKBP51 inhibition at disease onset confers lasting relief from sensory and emotional chronic pain symptoms

**DOI:** 10.1101/2024.11.01.621278

**Authors:** Sara Hestehave, Roxana Florea, Samuel Singleton, Alexander J.H. Fedorec, Sara Caxaria, Oakley Morgan, Katharina Tatjana Kopp, Laurence A. Brown, Tim Heymann, Shafaq Sikandar, Felix Hausch, Stuart N. Peirson, Sandrine M. Géranton

**Author notes:** **Correspondence to:** Sandrine M. Géranton; Medawar Building, Dept. of Cell & Developmental Biology, University College London, WC1E 6BT, United Kingdom., **Email:**. **Author Contributions:** <u>Study conception and experimental designs</u>: SH SG. <u>Data collection:</u> SH, RF, SG, SC; TH synthesised SAFit2. KK prepared the VPG. <u>Data analysis and interpretation:</u> SH, SG SSingleton, AF. <u>Manuscript preparation:</u> SH, SSingleton and SG wrote the initial draft of the manuscript. All authors read and approved the final manuscript. **Competing Interest Statement:** No competing interests exist.

## Abstract

Chronic pain affects 20-30% of the population and imposes a significant socio-economic burden as it is often accompanied by substantial emotional comorbidities such as anxiety and depression. Yet, the mechanisms underlying the interactions between the sensory and emotional aspects of chronic pain remain poorly understood. Here, we investigated the role of FKBP51, a regulator of the stress response, in mediating both sensory and emotional symptoms of chronic pain. Inhibition of FKBP51, *via* genetic deletion or pharmacological blockade, in persistent joint pain reduced fast-onset sensory, functional and activity-related symptoms, as well as late anxio-depressive comorbidities. FKBP51 inhibition after the establishment of the hypersensitive state provided only temporary symptoms relief, while acute inhibition at disease onset protected from the full development of sensory and anxio-depressive symptoms for up to 6 months. Our results also indicated that early pain symptoms could predict the late sensory and emotional outcomes of chronic pain. RNA sequencing of spinal cord tissue revealed that late FKBP51 inhibition transiently altered nociceptive genes associated with mechanical hypersensitivity. In contrast, early inhibition persistently downregulated the *Naaa* gene, a key regulator of the transition to chronic pain, and reorganized spinal cilia. Our results indicate that early FKBP51 inhibition after injury can persistently reduce chronic pain and prevent the onset of associated emotional comorbidities by modulating critical spinal neurobiological pathways that play pivotal roles in the transition to chronic pain.

**Significance statement:** Our study reveals that early inhibition of FKBP51, a modulator in the stress axis, at the onset of joint damage provides sustained pain relief and significantly delays or prevents emotional comorbidities in a sex-dependent manner. In contrast, FKBP51 inhibition initiated after chronic pain is established results in only temporary symptoms improvement. These findings highlight a critical therapeutic window during which timely intervention can prevent the transition from acute to chronic pain. By establishing a predictive link between early therapeutic response and long-term outcomes, this work has important clinical implications for proactive and personalized chronic pain management.

## Introduction

Despite recent advances in our understanding of the mammalian brain circuits that contribute to the different aspects of pain, our knowledge of the neurobiology of chronic pain is still limited and we are still seeking effective therapies that would improve patients’ quality of life. Patients with chronic pain often reports negative affective symptoms such as anxiety, depression and disrupted sleep (1–3) but we still do not understand how these emotional comorbidities may be mechanistically linked to the sensory outcomes of persistent pain. Improved understanding of the interactions between the circuits that drive the emotional and the sensory components of chronic pain may therefore shed further light on the mechanisms of pain chronicity.

In this context, FKBP51, a protein known to be involved in stress-modulation, was recently highlighted as a crucial driver of chronic pain in rodents, where it acts at the level of spinal pain circuits - known to generate widespread pain in chronic pain states - in a glucocorticoid dependent manner (4–6). Genetic variations in *FKBP5*, the gene encoding for FKBP51, influence the severity of musculoskeletal pain symptoms experienced by humans after trauma, suggesting a universal role for FKBP51 in chronic pain (7–9). FKBP51 regulates the stress response through modulation of the glucocorticoid receptor sensitivity. High levels of FKBP51 protein tend to be associated with a hyper-reactive stress pathway, as seen in patients with depression, while low levels of FKBP51 dampen down the stress pathway and produce a more robust and resilient individual (10). Supporting these observations, deletion of FKBP51 in healthy mice has been shown to reduce anxiety-related behaviour (11, 12) and improve sleep quality (13). Altogether, these observations suggest that blocking FKBP51 in chronic pain states could modulate many emotional-related symptoms. Therefore, following our previous success with FKBP51-manipulation to reduce hypersensitivity in persistent pain states (4, 5), our hypothesis was that targeting the protein FKBP51 modulates sensory, functional and emotional symptoms associated with chronic pain. Moreover, we have recently demonstrated that hypersensitivity in the early stage of persistent pain can predict the intensity of later emotional comorbidities (14). Therefore, we also anticipated that early FKBP51 inhibition would have long-term implications on emotional comorbidities.

We have developed the first selective antagonists of FKBP51 described so far (15). Acute administration of these antagonists recapitulates the anxiolytic properties of FKBP51 deletion in mice (11). We have previously tested pre-clinically one of these compounds, SAFit2, and found its administration at spinal cord level relieved hypersensitivity associated with persistent pain states to the same extend as global deletion of *Fkbp5* and intrathecal delivery of siRNA against *Fkbp5* (4, 5). Here, we use genetic and pharmacological blockade of FKBP51 in a robust pre-clinical model of persistent joint pain, the MIA model (14), that induces sensory and emotional symptoms, as experienced by chronic pain patients. We report that FKBP51 inhibition improves both the sensory and emotional component of persistent pain in a time dependent manner. Crucially, acute FKBP51 inhibition at disease onset provided permanent relief of both pain and emotional symptoms and dampened the expression of *Naaa*, a crucial driver of the transition from acute to chronic pain.

## Results

Study design and all statistical tests and full analysis results are presented in **SI Appendix Table S1, S2 andS3**, with statistically significant results summarized in the main manuscript and figures.

### MIA caused immediate changes in sensory and functional outcomes and daily activity rhythms but had a delayed impact on the open field and sucrose preference test

We first established the complete behavioural characterisation of male and female mice with joint pain induced by the injection of monoiodoacetate (MIA) in the knee joint, for up to 6 months. Mechanical allodynia, assessed by application of manual von Frey filaments, rapidly developed in both sexes and was maintained throughout the 6-month period (Fig.1A1,2), with no obvious sex differences (‘sex’ effect Fig.1A2, P=0.0632). Both males and females also showed immediate and robust changes in weight bearing, with the greatest impairment in the first 3 weeks and a more pronounced effect in females (Fig.1B1,2). We also looked at bone volume, as a surrogate measure of bone deformation, and found that the injection of MIA had led to a full remodelling of the joint by 6 months, across sexes (Fig.1C1,2).

Looking at the affective aspect of the pain experience, we found no difference in stress hormone levels between injured and control mice, either at 48h or 30 days after MIA injection, suggesting that stress-hormone increase, if any, would have been short-lived (Fig.S1A). However, the duration of affective-motivational response to von Frey stimulation was promoted by MIA, especially in the first 3 weeks after injection, with a greater impact in females, as demonstrated by significant interactions between sex and injury (Fig. S1B1,2, C1,2 and D1,2). This outcome measure was more variable than mechanical thresholds, likely due to its greater complexity thought to be driven by higher-order cognitive, emotional, and environmental factors (16, 17). Since most animals had returned to baseline for this specific behavioural assessment by 6 weeks, it was discontinued at that point to reduce burden on the animals. Overall, these results suggested that the sensory impact of the MIA injection was almost immediately stable, while the functional and affective impairments peaked during the first 3 weeks of the disease state.

We next found that both male and female mice had reduced preference for sucrose from day 90 and this lasted for at least 6 months (Fig.1D1,2), suggesting depressive-like behavior. We had previously reported that reduced preference for sucrose is not universal among animals with chronic joint pain, much like depressive symptoms in chronic pain patients (14). Using an individual profiling approach, where affected animals are defined as responding one standard deviation or more from the average performance of the control group (14, 18, 19), we found that up to 75% of male mice showed reduced sucrose preference at 4 months, while 100% of females exhibited this symptom by 6 months (Table S4). Moreover, both male and female mice showed reduced exploration time of the middle of the arena in the OFT, suggesting anxiety-like behaviour, from 3 months (27% of males and 45% of females, estimated by individual profiling, Table S4), with no obvious sex differences (Fig.1E1-3). Interestingly, both sucrose preference and time of exploration of the centre of the arena correlated with mechanical allodynia at 3 and 6 months across sexes (Fig.1F1,2). Furthermore, in line with our previous work establishing that early pain-related burden may serve as a predictor of later disease outcome (14), we also found that the early mechanical allodynia significantly correlated with sucrose preference and mechanical allodynia at 3 and 6 months (Fig.1G1, 1G2, and Table S5). Time of exploration in the centre of the arena was only significantly correlated to early allodynia at 3months, while trending at 6months (Fig.S1E and Table S5 for all prediction correlations).

**Figure 1:**
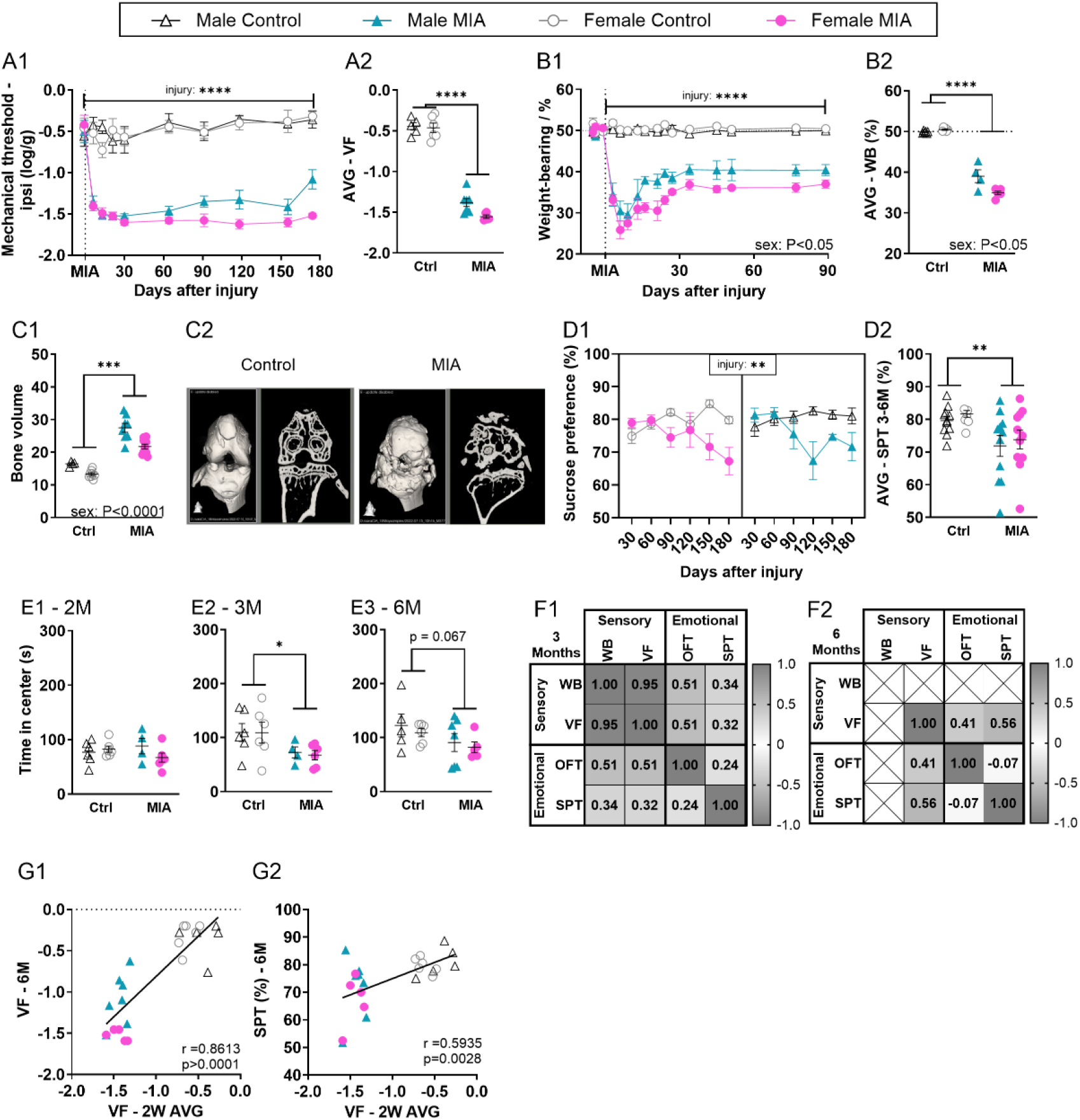
MIA induces immediate and persistent sensory and functional changes, but delayed anxio-depressive comorbidities. **(A1,2)** Mechanical hypersensitivity assessed using von Frey (vF) filaments (N=5-7). **(B1,2)** Functional deficit assessed using static weight bearing (WB) distribution (N=4-6). **(C1,2)** Bone volume around the joint, assessed using microCT at 6 months after injury (N=4-11). C2: Typical micro-CT image of control and MIA female joint. **(D1,2)** Depressive-like behaviour (anhedonia) assessed using the Sucrose Preference Test (two bottle test, water vs 1% sucrose). Data shows % consumption of the sucrose water compared with pure water. Weighted average (AVG) in D2 was calculated from 3-6months data. (N=4-12). **(E1,2,3)** Anxiety-like behaviour assessed using the Open Field Test measured at 2 (E1), 3 (E2) and 6 (E3) months after injury (N=4-7). **(F1,2)** Correlations between outcomes recorded from the same animal at the same time point at 3 (F1) and 6 (F2) months (N=22-23 across groups). **(G1,2)** Correlations between early-stage mechanical sensitivity (2-week weighted average) and the later stage (6 months) hypersensitivity (G1) and depressive-like (G2) behaviour. **A2, B2**: Weighted average (AVG) calculated across the full study period. Data shows mean ± S.E.M. NS= Not significant, *P<0.05, **P<0.01, ***P<0.001, ****P<0.0001, as determined by ANOVA-analysis. See full statistical analysis in SI Appendix Table S2.

Finally, we characterised the activity patterns of single housed mice using non-invasive motion sensor readouts validated against EEG (14, 20–22). Daily activity rhythms were no different between 8-week-old naïve male and female mice (Fig.2A,B; Fig.S2), but were significantly impacted by MIA in the early stages of the joint pain state, specifically during ZT14 and ZT1, where mice would normally be active. There was a significant increase in the proportion of time spent immobile, or in a sleep-like state, in that period, along with a significant reduction in activity in both male and female mice (Fig.2C,D). There was a noticeable sex-effect, as injured females could maintain activity levels for longer than injured male mice (Fig.2C,D, ZT14 to ZT1: significant time x injury x sex interactions; Table S2). Summary statistics of circadian rhythm confirmed these observations and indicated that injured mice were less able to sustain long bouts of activity (Fig.2E1) and had increased immobility in the dark-phase (Fig.2E2) and decreased immobility in light-phase (Fig.2E3). The impact of MIA on home-cage behaviour was short-lasted, as previously reported for other pre-clinical models of persistent pain (23), and, from one-month post injury, MIA-induced changes were subtle, while sex differences became more obvious as mice aged (Fig.S3, S4).

**Figure 2:**
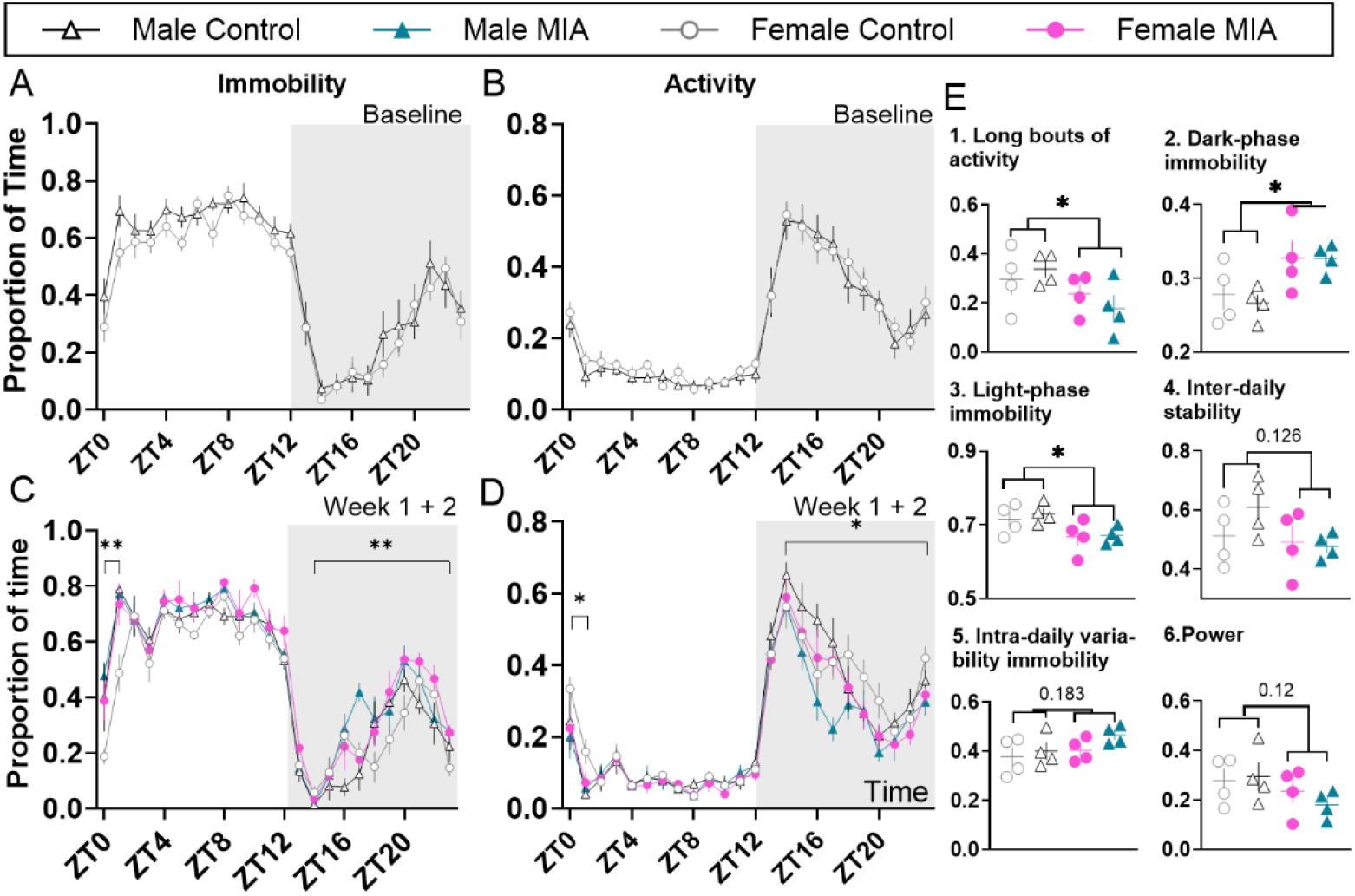
MIA-induced joint pain is accompanied by fragmented daily activity rhythms in the first 2 weeks. **(A-D)** 24h immobility/activity plots using 1h bins. **(A, B)** Average of 24h plots across the 6 days before MIA injection. N=8. **(C, D)** Average of 24h plots across the first 2 weeks after MIA injection. N=4. **(E) (E1)** proportion of time spent in long bouts (>10min) activity during the dark period, **(E2)** proportion of time spent in immobility during the dark phase, **(E3)** proportion of time spent immobile during the light phase, **(E4)** interdaily stability as a stability index, with higher values representing highly stable patterns, **(E5)**, intra-daily variability immobility as a variability index, with higher values representing highly fragmented patterns and **(E6)** strength of the rhythmicity; power of the 24-hour activity cycle calculated across 2 weeks. The power of a periodogram provides a measure of the strength and regularity of the underlying rhythm, with higher values indicating robust rhythms. In circadian disruption—where rhythms are typically less robust—the power is expected to be reduced, which may indicate the absence of a significant circadian rhythm. N=4. ZT0=7am, ZT12=7pm. Data shows mean ± S.E.M. *P<0.05; ** P<0.01, injury effect. See full statistical analysis in SI Appendix Table S2.

### *Fkbp5* global deletion reduced MIA-induced sensory, functional and depressive-like symptoms

We next asked whether the inhibition of FKBP51 could reduce the pain phenotype beyond its documented effects on mechanical hypersensitivity (4–6, 24). First, we used *Fkbp5* global KO mice (4, 5). Both male and female *Fkbp5* KO displayed MIA-induced mechanical allodynia and weight bearing deficit that lasted for 6 months but was substantially reduced compared with control WT littermates (Fig.3A1,2; B1,2). There were no sex*genotype-interaction affecting the sensory and functional outcomes, suggesting that the *Fkbp5* deletion did not affect the two sexes differently. However, as observed in Fig.1, females displayed higher functional deficits across genotype, resulting in a sex-effect in the overall statistical analysis for this dataset (Table S2). Importantly, *Fkbp5* deletion had no impact on MIA-induced bone deformation after correction for body weight, important due to the leaner phenotype of the KO mice (25, 26) (Fig.3C, Fig.S5A1,2; B). We also observed that *Fkbp5* knock-down mitigated the MIA-induced decrease in sucrose preference across sexes, as up to 55.6% and 44.4% of males and females KO mice respectively displayed a resilient phenotype (Fig.3D1,2, Table S6). *Fkbp5* deletion had a milder impact on time of exploration in the centre of the OFT arena (Fig.3E, Table S6). While these results might have been confounded by increased locomotion in the KO mice (Fig.S5C), they were confirmed by EPM (Fig.S5D). Overall, there was no significant correlation across sex and genotype between sensory and emotional outcomes, when compared at any given time point (Fig.3F1,2). Moreover, while early mechanical allodynia was a significant predictor of late pain sensitivity (Fig.3G1), it could not predict sucrose preference and time of exploration of the centre of the arena at 3 or 6 months (Table S5, Fig 3G2,3). These results suggested that *Fkbp5* global knockdown disrupted the relationship between pain burden and emotional comorbidities, likely due to the reported impact of *Fkbp5* deletion on anxiety- and depressive-like behaviour independently from the hypersensitive state (26–28).

**Figure 3:**
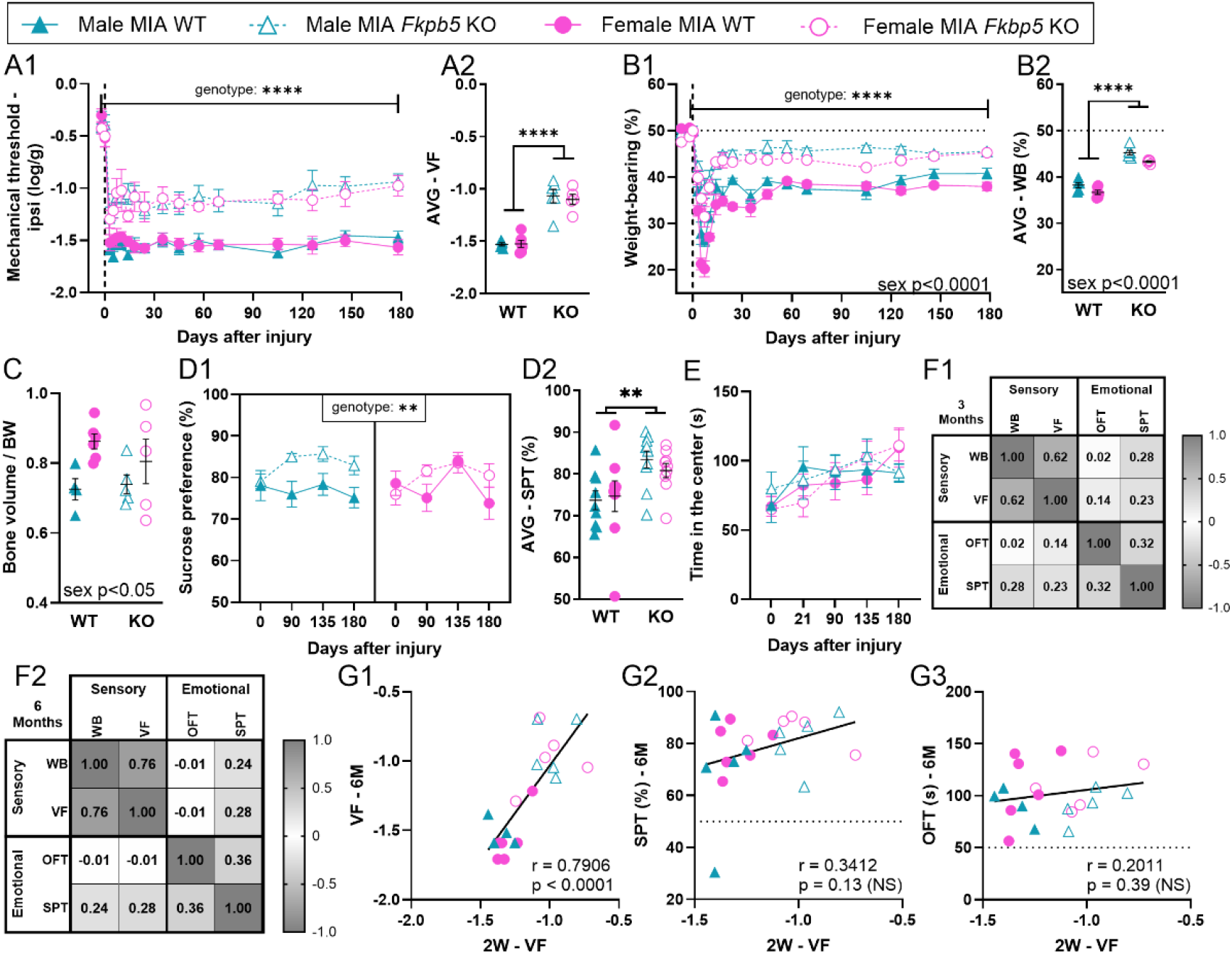
*Fkbp5* global knock-out improves sensory, functional and depressive-like behaviour induced by MIA. **(A1,2)** Mechanical hypersensitivity assessed using von Frey filaments (N=5-6). **(B1,2)** Functional deficits assessed using static weight bearing distribution (N=5-6). **(C)** Bone volume around the joint assessed by microCT at 6 months after injury and normalised to body weight (N=4-6). **(D1,2)** Depressive-like behaviour (anhedonia) assessed using the Sucrose Preference Test (two-bottle test, water vs 1% sucrose). Data shows % consumption of the sucrose water compared with pure water. Weighted average (AVG) in D2 was calculated from 3-6months data. (N=5-9). **(E)** Anxiety-like behaviour assessed using the Open Field Test (N=5-6). **(F1,2)** Correlations between outcomes recorded from the same animal at the same time point at 3 (F1) and 6 (F2) months (N=22 across groups). **(G1,2,3)** Correlations between early-stage mechanical sensitivity (2-week weighted average) and the later stage (6 months) of hypersensitivity (G1), depressive-like (G2) and anxiety-like (G3) behaviour. **A2, B2**: Weighted average (AVG) calculated across the full study period. Data shows mean ± S.E.M. *P<0.05, **P<0.01, ***P<0.001, ****P<0.0001, as determined by ANOVA analysis. See full statistical analysis in SI Appendix Table S2.

We also characterised the impact of *Fkbp5* knockdown on daily activity rhythms. While there were no obvious sex and genotype differences in 8-week-old naive mice (Fig.S6A, B, S7), *Fkbp5* KO male and female mice showed increased activity between ZT14 and ZT1 after MIA, when compared with WT mice (Fig.S6C,D). These results suggested that the deletion of *Fkbp5* provided some protection from the injury-induced disruption to daily activity rhythms. Moreover, MIA-induced disruption in a number of summary statistics of circadian parameters (Fig.2E) were resolved by *Fkbp5* knockdown, including in long-bouts of activity (Fig.S6F). As mice aged and at a time where the effects of the injury were no longer obvious in WT mice, genotype and sex differences grew stronger (Fig.S6). In particular, the KO females were more active in the dark phase and less immobile in the light phase than their wild–type counterparts (Fig.S8, S9). Overall, these results suggested that the impact of global *Fkbp5* knockdown on immobility and activity patterns went beyond the protection from MIA-induced disruptions.

### Pharmacological inhibition of FKPB51 after establishment of persistent pain provided temporary improvement of sensory, functional and depressive-like symptoms

We next asked whether pain relief could be achieved through pharmacological blockade of the protein FKBP51 using the FKBP51 inhibitor, SAFit2 (4, 5). SAFit2-VPG, a formulation designed for the sustained released of the inhibitor over 7 days via encapsulation in a vesicular phospholipid gel (VPG), was first administered 2 weeks after the MIA injection, a time point when SAFit2 had been tested in models of ankle joint inflammation and neuropathic pain (4). SAFit2-VPG attenuated the MIA-induced allodynia, with mechanical thresholds nearly returning to baseline (Fig.S10A). Weight bearing deficit were equally improved (Fig.S10B). Next, we delayed the administration of SAFit2-VPG to 1 month after MIA-induction, and, using weekly injections, we extended the duration of FKBP51 inhibition to 3 weeks. SAFit2-VPG significantly improved the MIA-induced mechanical allodynia (Fig.4A1,2) and weight bearing deficits (Fig.4B1,2), during its controlled release from VPG, when it is therapeutically effective.

**Figure 4:**
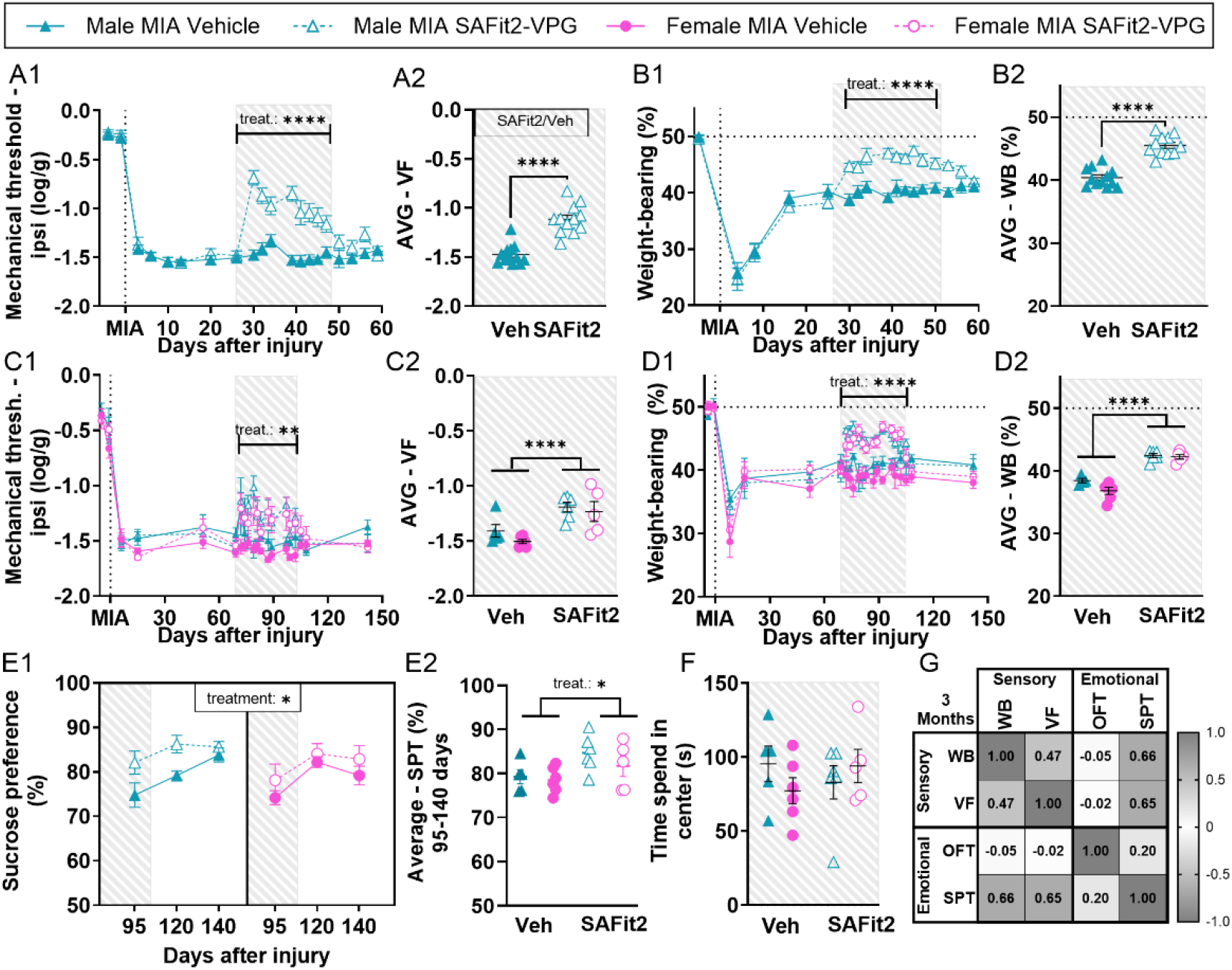
Pharmacological inhibition of FKBP51 after MIA induction transiently improves sensory and functional deficits and delays the onset of depressive-like behaviors. Hatched grey area indicates when SAFit2 was active. (**A1,2**) Mechanical hypersensitivity assessed in male mice using von Frey filaments. (N=12). (**B1,2**) Functional deficits assessed using static weight bearing distribution (N=11-12). (**C1,2**) Mechanical hypersensitivity assessed using von Frey filaments. SAFit2-VPG was administered 10-14 weeks after injury. (N=5-6). (**D1,2**) Functional deficits assessed using static weight bearing distribution. (N=4-6). (**E1,2**) Depressive-like behaviour (anhedonia) assessed using the Sucrose Preference Test (two-bottle test, water vs 1% sucrose). Data shows % consumption of the sucrose water compared with pure water. Weighted average in E2 was calculated from D95-140 data (N=5-6). (**F**) Anxiety-like behaviour assessed using the Open Field Test (N=5-6). (**G**) Correlations between outcomes recorded from the same animal at the same time point at 3 months after injury (14 weeks), when SAFit2-VPG was administered from 10-14 weeks after injury (N=21 across groups). A2, B2, C2, D2: Weighted average (AVG) was calculated for the hatched grey study period, where the compound was active. Data shows mean ± S.E.M. *P<0.05, **P<0.01, ***P<0.001, ****P<0.0001, as determined using ANOVA analysis. See full statistical analysis in SI Appendix Table S2.

We next administered SAFit2-VPG for 4 weeks during the 2- to 3-month period after MIA injection, when long-term emotional disorders manifest (Fig.1E, F). SAFit2-VPG improved mechanical threshold and weight bearing deficit across sexes (Fig.4C1,2, D1,2) and delayed the MIA-induced changes in sucrose preference (Fig.4E1,2), with up to 83.3% of males and 60% of females injected with SAFit2-VPG being resilient (Table S7). Unexpectedly, SAFit2-VPG had no impact on the OFT exploration (Fig.4F, Fig.S10C-D, Table S7), nor did it affect locomotor activity (Fig.S10E) or body weight (Fig.S10F). Furthermore, at 3 months, while SAFit2 was still in its active release phase, there were significant correlations between improvement of both functional and sensory outcome measures and the effect on sucrose preference (Fig.4G). While mechanical thresholds quickly returned to control levels when the treatment was discontinued, the effects on sucrose preference were maintained for longer, with significant predictive correlations between the mechanical thresholds measured during the active phase of SAFit2-treatment and the sucrose preference at 3, but not 6, weeks after treatment-end (Fig.S10G,H; Table S5).

Surprised by the absence of effect of SAFit2-VPG on the OFT in injured mice, we explored the effect of prolonged SAFit2-VPG in naïve mice. In a control experiment, SAFit2-VPG was administered to naïve mice for one month. This reduced the time spent in the centre of the OFT arena (Fig.S11A), which may explain its inability to prevent the MIA-induced reduction in centre time. Finally, SAFit2 had no effect on mechanical threshold (Fig.S11B) or depressive-like behaviour in naïve mice (Fig.S11C, D).

### Inhibition of FKBP51 at disease onset prevented the full development of MIA-induced symptoms

We next explored the effect of early blockade of FKBP51 during the onset of the pain state and observed a reduction in MIA-induced allodynia and both static and dynamic weight bearing deficits (Fig.5A,B,C) of similar effect size as when SAFit2-VPG was given 2 weeks and 4 weeks after MIA (Fig.S12A-F). Remarkably, for up to 6 months after injury, mechanical threshold and weight bearing never reached the levels seen in vehicle treated MIA animals, despite SAFit2 being only active until day 14. The early SAFit2 treatment resulted in a sensory profile similar to that of the *Fkbp5* KO mice. Importantly, early SAFit2 treatment did not exacerbate nor prevented the degradation of the knee bone induced by MIA (Fig.S12G).

**Figure 5:**
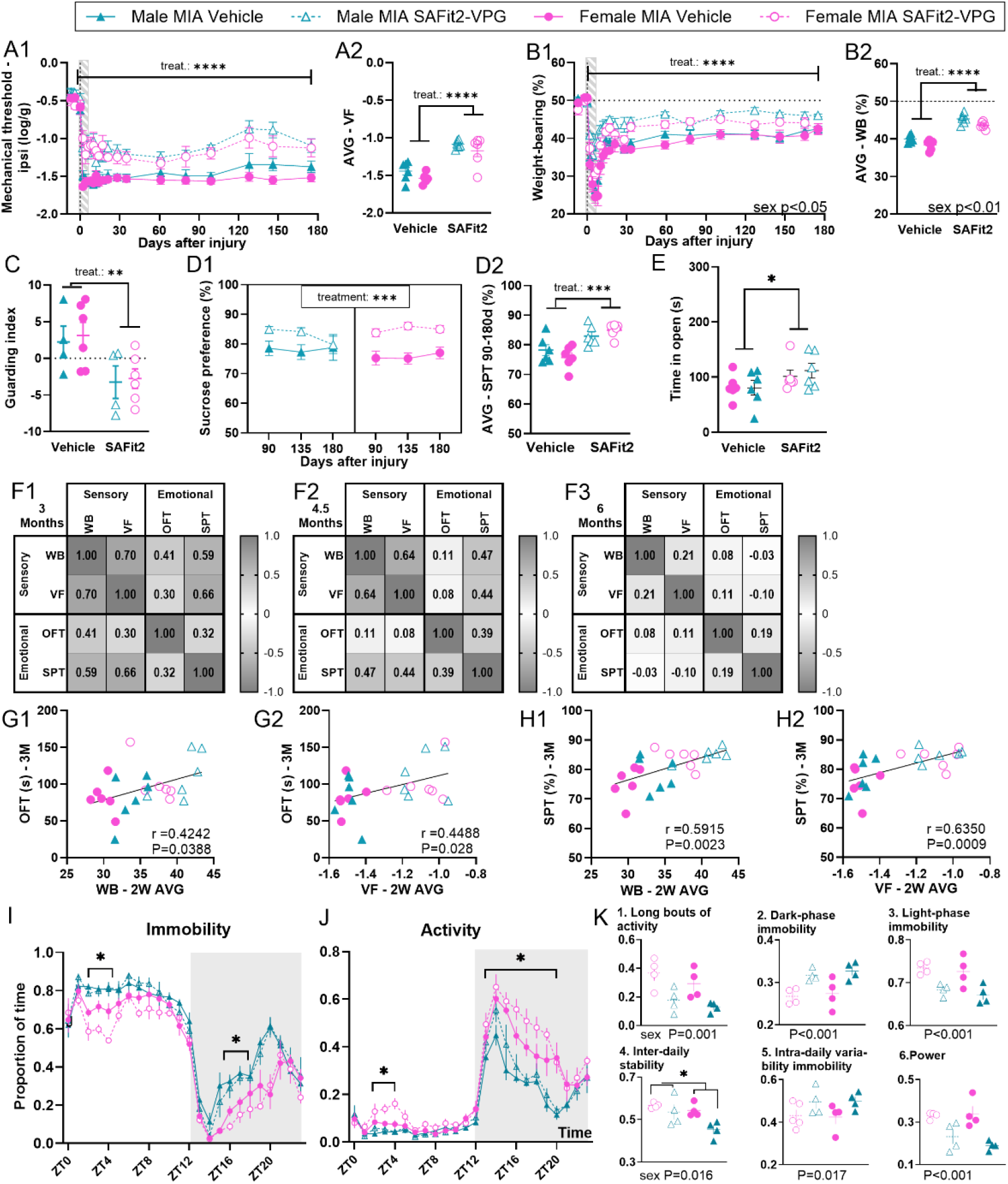
Acute pharmacological inhibition of FKBP51 initiated during joint pain onset prevents the full development of MIA-induced symptoms. SAFit2-VPG was injected 3 days before MIA. Hatched grey area indicates when SAFit2 was active. (**A1,2**) Mechanical hypersensitivity assessed using von Frey filaments (N=6). (**B1,2**) Functional deficits assessed using static weight bearing distribution (N=6). (**C**) Guarding index at 6 weeks after injury, 4 weeks after SAFit2 was active, assessed by Catwalk gait analysis (N=4-6). (**D1,2**) Depressive-like behaviour (anhedonia) assessed using the Sucrose Preference Test (two-bottle test, water vs 1% sucrose). Data shows % consumption of the sucrose water compared with pure water. Weighted average (AVG) in D2 was calculated from 3-6months data (N=6). (**E**) Anxiety-like behaviour assessed using the Open Field Test (N=6). (**F1,2,3**) Correlations between outcomes recorded **from** the same animal at 3 (**F1**), 4.5 (**F2**) and 6 (**F3**) months after injury (N=6). (**G, H**) Correlations between the weighted average weight bearing asymmetry (WB) or mechanical hypersensitivity (vF) from the first two weeks after injury, when the compound was active, and the level of anxiety-like behaviour at 3 months (**G1,2**), and the level of depressive-like behaviour at 3 months (**H1,2**). **(I, J)** 24h immobility/activity plots using 1h bins across the first 2 weeks after MIA injection. N=4. **(K)** Average across 14 days after MIA. **(K1)** Proportion of time spent in long bouts (>10min) activity during the dark period, **(K2)** proportion of time spent immobile during the dark phase, **(K3)** proportion of time spent immobile during the light phase, **(K4)** interdaily stability as a stability index, with higher values representing highly stable patterns, **(K5)** intra-daily variability immobility as a variability index, with higher values representing highly fragmented patterns and **(K6)** strength of the rhythmicity; power of the 24-hour activity cycle calculated across 2 weeks. A2, B2: Weighted average (AVG) was calculated for the full study period. Data shows mean ± S.E.M. *P<0.05, **P<0.01, ***P<0.001, ****P<0.0001, as determined using ANOVA-analysis. Full statistical analysis in SI Appendix Table S2.

Early FKBP51 inhibition reduced affective responses to von Frey filaments (Fig.S13A, B, C) and brush-induced allodynia (Fig.S13D) seen early after the initiation of the pain state. We also found that the early treatment prevented the reduction in both sucrose preference and time spent in the centre of the OFT arena (Fig.5D1,2, E, Fig.S13E) at 3 months after injury. Specifically, the drop in sucrose preference in male mice was delayed until 6 months, with up to 66.7% of male mice treated with SAFit2-VPG resilient at 3 months (Table S8). There was no drop in sucrose preference in female mice, with up to 100% of female mice treated with SAFit2-VPG resilient (Table S8). Finally, the early SAFit2-VPG administration did not lead to any change in body weight compared with vehicle treated mice (Fig.S13F) and had no impact on distance travelled in the OFT (Fig.S13G).

Pain-related outcomes (allodynia and weight bearing) showed the strongest correlations with sucrose preference and exploration of the centre of the OF arena at 3 months across sexes, weakening from 4.5 months (Fig.5F). SAFit2’s early effects on pain-related measures predicted late pain experience, as shown by correlations between early mechanical threshold and weight bearing and late mechanical thresholds (Table S5), time spent in the centre of the OF arena (Fig.5G1,2), and sucrose preference (Fig.5H1,2), with a predictive strength highest at 3 months (Table S5).

Finally, SAFit2-VPG administered during the onset of the pain state had a protective effect on the MIA-induced disruption to daily activity rhythms, as SAFit2-VPG-treated MIA mice slept less and were more active than vehicle treated controls (Fig.5I,J). However, the impact of FKBP51 inhibition on summary parameters of activity rhythms were milder and rarely reached significance (Fig.5K), and as mice aged, sex differences became more obvious (Fig.S14). As we had observed an impact of SAFit2-VPG in the OFT in naïve mice, we also looked at activity patterns in naïve mice treated with SAFit2-VPG and found an increase in activity in female and a decrease in male mice during their active phase (ZT14 to ZT1, sex*treatment interactions; Fig.S15A,B) but with no significant differences in the summary parameters of circadian disruption (Fig.S15,C-N). Overall, the impact of early and acute FKBP51 pharmacological inhibition on activity rhythms were milder than the global knock-down of *Fkbp5*, still improving some of the MIA-induced deficits at peak burden.

### FKBP51 inhibition at disease onset downregulated genes involved in the transition to chronic pain and reorganization of spinal cilia

We used RNAseq to explore the mechanisms behind early *vs* late FKBP51 inhibition. Since SAFit2 reduces pain sensitivity across all persistent pain states regardless of their origin, we focused on comparing SAFit2 *vs* vehicle in MIA-injured mice, removing the confound of MIA. One group received SAFit2-VPG or vehicle at pain onset (early), and another after pain was established (late). As before, mice that received early SAFit2 treatment maintained mechanical thresholds higher than controls beyond the compound’s active phase (Fig.6A1,2). By day 30 post-MIA, mechanical thresholds were similar across all SAFit2-VPG treated mice. Early treatment however yielded greater dynamic weight-bearing improvement (Fig.6B, S16A-D). Corticosterone levels were reduced by SAFit2-VPG (Fig.6C) and correlated with the level of allodynia (Fig.6D), supporting a functional relationship between FKBP51 activity, corticosterone regulation and mechanical hypersensitivity, as previously reported (4, 5).

**Figure 6:**
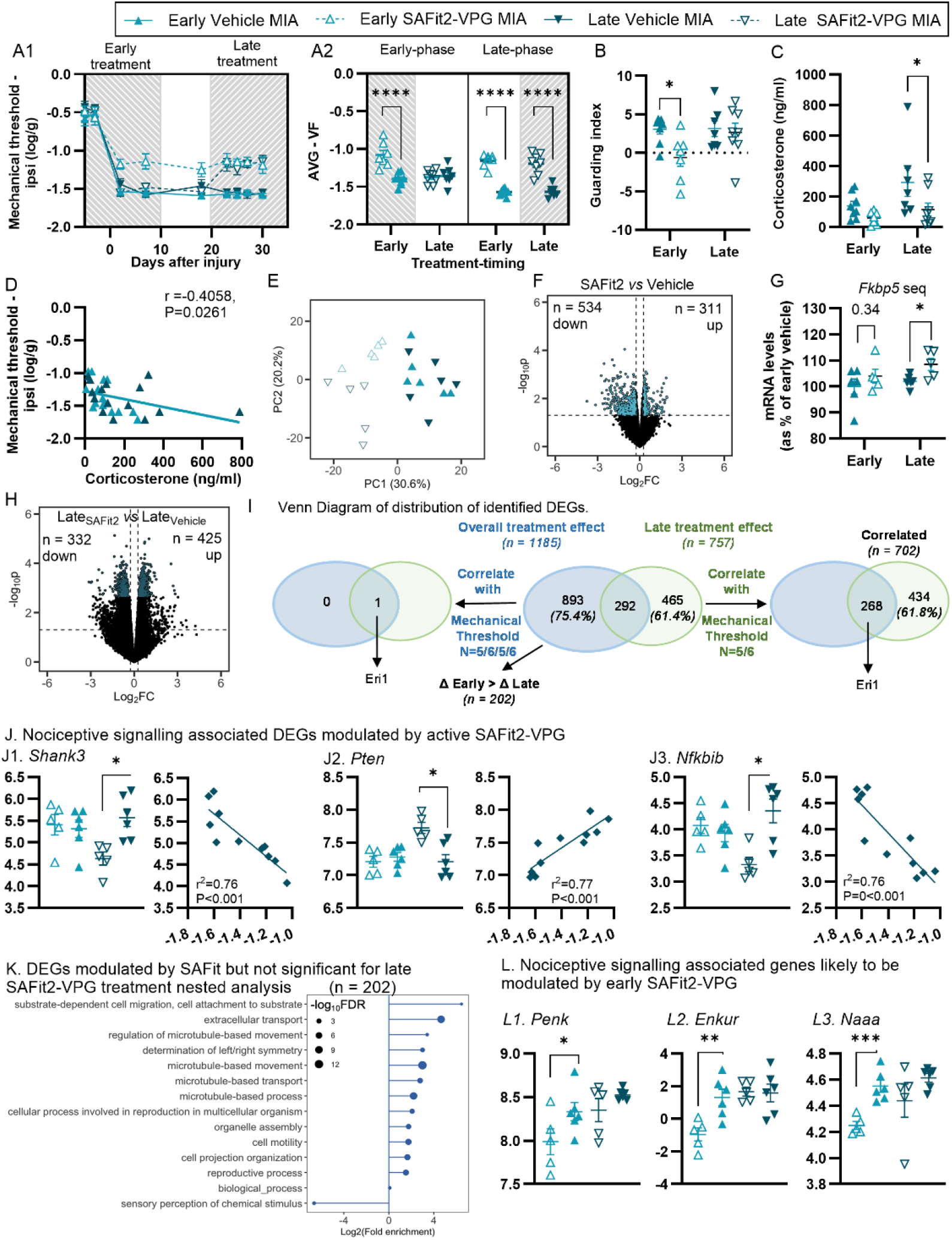
Pharmacological inhibition of FKBP51 initiated either before or after pain state induction leads to different gene expression changes. SAFit2-VPG was injected either “Early” (day –3 and day 4), or “Late” (day 20 and day 27). Hatched grey area indicates the active phase of SAFit2. (**A1,2**) Mechanical hypersensitivity assessed using von Frey filaments. **A1**: Data shows Mean ± S.E.M. (**B**) Guarding index assessed through catwalk gait analysis. (**C**) Corticosterone levels (Day 31). (**D**) Correlation between mechanical allodynia and corticosterone (Day 31). (**E**) PCA Analysis indicating sub-clusters within the second principal component of SAFit2-treated groups not present in vehicle-treated groups. (**F**) Volcano plots; x-axis: log2-transformed fold change in gene expression between SAFit2 and vehicle treated groups. (**G**) Expression levels of *Fkbp5* mRNA from sequencing counts. (**H**) Volcano plots; x-axis: log2-transformed fold change in gene expression between Late_SAFit2_ and Late_vehicle_ treated groups; (**I**) Venn Diagram of distribution of identified DEGs. **(J)** DEGs by SAFit2-VPG late treatment associated with nociceptive signalling. *P<0.05, as per nested analysis; correlation graphs show the results of Pearson Correlation analysis. y-axis: log_2_CPM, x-axis: mechanical thresholds (log(G)). (**K**) Lollipop plot of enriched Gene Ontology (GO) terms identified by PANTHER from the differentially expressed genes (DEGs) between SAFit2 and vehicle treated mice;,emphasis on genes with a greater change in expression between (Early_SAFit2_ vs Early_Vehicle_) than (Late_SAFit2_ vs Late_Vehicle_). The X-axis represents the GO terms related to biological processes while the Y-axis shows the fold change in enrichment. Significance (-log10 adjusted P-value) of each term is denoted by size. (**L**) Selected genes associated with nociceptive signalling. Y-axis: log_2_CPM. (**F,H**): y-axis represents the –log10(p-value) for differential expression; coloured points are significantly up- or down-regulated with threshold abs(log2FC) >1.2; vertical dotted lines represent log2FC thresholds: left: log2FC=-1 (2-fold downregulation) and right: log2FC=1 (2 fold upregulation); horizontal dotted line represents the p-value threshold. *P<0.05; **P<0.01; ***P<0.001, t-test Early_SAFit2_ vs Early_Vehicle._ For all graphs; (N=7-8), See full statistical analysis in SI Appendix Table S2.

RNA sequencing of the ipsilateral dorsal horn quadrants aimed to identify differentially expressed genes (DEGs) by SAFit2. We initially explored the main effect of treatment using linear regression modelling of early and late SAFit2 groups compared with vehicle-treated groups and detected 1185 DEGs (nominal P<0.05). Principal component analysis revealed sub-clusters within the second principal component of SAFit2-VPG-treated groups exposed at the early *vs* late time points absent in vehicle-treated groups, suggesting small time-dependent regulation of gene expression by SAFit2 (Fig.6E). Out of the 1185 DEGs, 311 were up-regulated and 534 were downregulated by SAFit2-VPG with a cut-off fold-change of 1.2 (Fig.6F, total 845 DEGs). Notably, *Fkbp5* was one of the 311 up-regulated DEGs (Fig.6G).

To focus on pathways modified during the drug’s active phase, we focused on the late treatment group. A nested analysis (Late_SAFit2_ *vs* Late_Vehicle_) revealed 757 DEGs (adjusted FDR P<0.05, cut-off fold-change of 1.2), with 425 upregulated and 332 downregulated genes (Fig.6H). We next hypothesized that if the DEGs were functionally relevant to the reduced mechanical threshold observed in Late_SAFit2_ treated mice, their expression level would correlate with the mechanical thresholds at time of dissection. Pearson’s correlation analysis with Bonferroni correction identified 702 DEGs that correlated with the mechanical threshold (average day 27 to day 30) but not with the guarding index (Fig.6B), suggesting that these genes were more likely linked to the sensory rather than the functional aspect of the pain state. 434 of the 702 identified DEGs were not among the initial 1185 DEGs (Fig.6I), suggesting transient modulation by SAFit2-VPG. Gene ontology (GO) analysis highlighted roles in energy metabolism, cell structure regulation, neuronal development, RNA processing, protein transport and sensory perception (Fig.S17). Downregulated DEGs linked to reduced hypersensitivity included *Shank3*, linked to autism spectrum disorder and pain signalling (29, 30) (Fig.6J1). Conversely, upregulated *Pten*, the gene encoding the Phosphatase and Tensin Homolog, may protect from neuropathic pain (31) (Fig.6J2), while inhibition of the NF-KB inhibitor *Nfkbib* suggested enhanced NF-KB signalling (Fig.6J3). However, some pain related genes were regulated in ways potentially reinforcing hypersensitivity (Fig.S18).

We next investigated DEGs that were at the intersection of the 2 analyses (SAFit2 overall effect - n=1185 and late treatment nested analysis - n=757: n=268 DEGs, Fig.6I). Of these, only one (Eri1, involved in RNA metabolism) correlated with mechanical hypersensitivity when both treatment groups were considered all together (Fig.6I). Several DEGs linked to nociceptive signaling were also identified (Fig.S19), but their regulation alone could not explain the long-term effects of early SAFit2 treatment, as late-treated animals were expected to return to full hypersensitive state after the drug’s active phase.

To uncover mechanisms of long-term pain relief driven by early SAFit2 treatment, we first drew contrasts between Early_SAFit2_ and Early_Vehicle_ treated mice but found no DEGs with FDR-corrected P<0.05. We then focused on the 1185 DEGs that were not modified by the late treatment alone (*i*.*e*. 1185 Δ 757; n=893 DEGs; Fig.6I). GO analysis revealed that these 893 DEGs were mainly related to cilia motility and cytoskeleton organisation (Fig.S20). Further analysis comparing the change in expression (SAFit2 Δ Vehicle) between early and late treatments identified 202 annotated DEGs with greater modulation by early SAFit2 (Fig.6I). While none of these genes correlated with mechanical thresholds, 32 of them correlated with the guarding index which we found to be more strongly influenced by early SAFit2 treatment.

Many were associated with cilium-related processes (Fig.6K), while several genes previously linked to pain phenotype (Fig.6L) were durably reduced by early SAFit2. These included the preproenkephalin gene *Penk*, which encodes precursors to the endogenous opioid peptides enkephalins, *Enkur*, which encodes the protein enkurin, known to interact with TRPC channels and *Naaa*, which encodes the enzyme N-acylethanolamine acid amidase, and crucially had been previously implicated in the transition from acute to chronic pain(32).

Altogether, our RNAseq analysis suggested early SAFit2 treatment induced lasting spinal cord gene expression changes absent in late treatment, potentially driving sustained pain relief. While FKBP51 is predominantly expressed in neurones in the rodent spinal cord (5), identified DEGs were likely to be expressed in various cell types but changes reported here could not be attributed to altered cell type composition (Fig.S21).

## Discussion

Our study provides a comprehensive analysis of the behavioural and molecular impact of FKBP51 inhibition in a persistent pain state (see Fig.S22: Summary of Findings). The MIA model, which replicates key histological and functional alterations seen in osteoarthritis (OA), allowed us to investigate both sensory and emotional dimensions of pain (33–39). MIA injection rapidly produced a hypersensitive state lasting at least 6 months in male and female mice, alongside progressive anxiety- and depressive-like behaviours. These emotional comorbidities emerged in a subset of animals, as seen in chronic pain patients (40). This temporal dissociation between immediate sensory changes and delayed emotional disturbances is consistent with previous clinical and pre-clinical observations indicating a gradual onset of emotional symptoms in chronic pain states (14, 40–45). Small but significant sex differences were found in pain sensitivity and functional deficits, with females more affected by MIA, while emotional comorbidities showed more complex sex effects.

Regardless of sex differences, both sensory and functional aspects of late-stage joint pain correlated with emotional comorbidities across sexes, as seen in patients, where anxiety and depression are positively associated with pain severity (45). More importantly, early sensory and functional measures correlated with late pain intensity and depressive-like behaviours, as we had previously observed in males (14). Further strengthening the idea that the early pain burden could serve as a predictor of late emotional outcomes, FKBP51 inhibition during disease onset not only permanently blunted MIA-induced allodynia, as observed in a trauma-induced pain model (6), but also substantially reduced late anhedonia and anxiety-like behaviour. Since pain behaviours in the MIA model peak early, this effect may have resulted indirectly from reducing peak pain levels rather than from the timing of the intervention itself.

As many other pain management options, FKBP51 inhibition after the establishment of the pain state (between 2 weeks and 2 months post-onset) provided transient benefit only, rather than a long-lasting cure, likely through the modulation of central signalling pathways. Overall, our results support our original hypothesis, that FKBP51 inhibition - which we previously showed improves injury-induced mechanical hypersensitivity in a glucocorticoid-dependent manner (5) - would alleviate both the sensory and emotional impact of persistent pain. When FKBP51 inhibition was provided at 2-3 months, just before the typical onset of comorbidities, it effectively delayed depressive-like behaviour. This suggests that onset of depressive comorbidity in chronic pain is likely to be secondary to sensory and functional effects, as reported by others (46). It also suggests that the impact of FKBP51 inhibition on pain-induced depression is secondary to pain improvement, supported by a significant correlation between the early pain intensity and the later depressive comorbidity. Nonetheless, a direct effect of FKBP51 inhibition on depressive-like behaviour cannot be excluded.

It is noteworthy that long-term FKBP51 blockade *(i*.*e*. for 4 weeks) in injured mice had no impact on MIA-induced anxiety-like behaviour. This may not be surprising as long-term SAFit2 treatment alone in naïve mice caused anxiety-like behaviour, unlike the acute administration that is known to have anxiolytic effects (11, 12). Any anxiolytic effects of SAFit2 resulting from pain relief in injured mice may therefore have been counteracted by the anxiogenic effects observed after long-term use. While SAFit2 never worsened anxiety-like behaviour in injured mice, it may however have significant impacts on circadian patterns. Indeed, circadian alterations outlasted the short-lived MIA-induced changes in the KO mice with no clear benefit on immobility defined sleep. Nonetheless, FKBP51 has been identified as a target for stress-induced sleep disorders, and its deletion promotes better sleep in disease states (13). Importantly, acute SAFit2-VPG treatment at disease onset had no long-lasting impact on sleep-like immobility and activity, suggesting stronger therapeutic potential for an early and acute FKBP51 targeted pharmacotherapy.

Overall, acute pharmacological inhibition of FKBP51 at the onset of the disease provided effective and long-lasting pain relief, which continued beyond the drug’s active phase. To elucidate the mechanisms underlying these observations, we investigated gene expression changes induced by SAFit2 treatment given at the onset or during the maintenance phase of the pain state in MIA mice, avoiding the confound of MIA-induced changes (for alternative approach see Table S9). We focused on the spinal cord, specifically the ipsilateral quadrant of the superficial dorsal horn, as our original work using spinal specific genetic and pharmacological manipulations of *Fkbp5* suggested that FKBP51 drives chronic pain through spinal mechanisms (5). While unlikely, as MIA injection into the knee joint does not increase *Fkbp5* expression in DRGs (Fig.S23), the effect of systemic FKBP51 inhibition observed in our study could nevertheless be secondary to changes in peripheral tissues.

Genes uniquely modified by active SAFit2, and therefore transiently modulated by FKBP51 inhibition, often correlated with mechanical hypersensitivity, suggesting some potential functional relevance. These belonged to various biological processes including sensory perception and a number had been previously involved with nociceptive signalling including *Shank3 and Pten*. Surprisingly, we did not observe changes in glucocorticoid signalling, despite SAFit2’s impact on corticosterone levels. However, this effect was relatively mild, and previous studies have also reported mild influence of SAFit2 on stress hormone levels (47). On the other hand, we observed a decrease in *Nfkbib* expression, indicating that SAFit2 may promote NF-κβ signalling. The role of FKBP51 in this context is complex, with previous studies suggesting that it could both inhibit and facilitate NF-κβ signalling in various cell types (48).

While no genes were significantly modulated by early SAFit2 treatment alone (FDR P<0.05, 1.2-fold), we identified DEGs modulated by SAFit2 overall that were not significantly changed by the late treatment. This suggested that the early treatment likely drove the differential regulation. Most DEGs showing greater changes after early treatment were associated with cilium movement and assembly. The primary cilium is a key organelle in cellular signaling, present in most post-mitotic cells, including neurons in the CNS. While its role in spinal neurons has been understudied, cilia-related genes have recently been linked to nociceptive signaling.

Primary cilia in both large and small neurons of the dorsal horn have been described (49) and their role in Sonic Hedgehog (Shh) signaling, which modulates pain, has been highlighted in nociceptors (50). Our findings suggest that long-term re-organisation of spinal cilia following early SAFit2 treatment may provide long-term pain relief. Interestingly, it was recently suggested that the µ-opioid receptor selectively localize to primary cilia in the mouse CNS (51), which may contribute to the phenotype observed in early treated mice. We also found that early SAFit2-VPG treatment persistently reduced the expression of *Penk*, the gene encoding for preproenkephalin, while *Penk* derived peptides have been shown to delay recovery from inflammatory hypersensitivity (52). Finally, one non cilium related DEG particularly stood out, the *Naaa* gene which encodes the enzyme N-acylethanolamine acid amidase (NAAA). NAAA has been identified as a crucial control point in the progression to pain chronicity and inhibition of NAAA in the spinal cord after peripheral injury during a 72h time window can prevent chronic pain (32). Here, we found that SAFit2-VPG treatment in a similar time window after MIA injection led to a long-lasting downregulation of *Naaa* compared with controls and mice that received SAFit2-VPG later.

Our study reveals key neurobiological mechanisms underlying the modulation of persistent pain through FKBP51 inhibition, focusing on spinal circuits and molecular pathways. Early FKBP51 inhibition induced long-lasting changes in nociceptive signaling, including the downregulation of *Naaa*, a gene pivotal in the transition to chronic pain, and reduced expression of *Penk*, which encodes peptides delaying recovery from hypersensitivity. We identified genes associated with cilia function and assembly, suggesting that primary cilia in spinal neurons may play a critical role in pain modulation through pathways such as Sonic Hedgehog (Shh) signaling and possibly μ-opioid receptor localization. These findings provide novel insights into the spinal mechanisms governing the sensory and emotional dimensions of persistent pain, emphasizing the importance of early molecular changes in shaping long-term pain outcomes.

## Materials and Methods

Extended materials and methods can be found in SI Appendix.

### Animalsandhousing

Male and female mice were used at 8-10 weeks of age, either C57Bl/6J, Charles River, UK or *Fkbp5* KO colony, maintained in house (C57Bl/6J and Swiss Webster background). All experiments comparing wild-type (WT) and knockout (KO) animals were performed using WT and KO littermates bred in our colony. Mice were kept in groups of 2-4 in a temperature- and light-controlled environment with *ad libitum* provision of food and water. Experiments were conducted under the Home Office License P8F6ECC28 and ARRIVE guidelines were followed for reporting.

### Compoundpreparationandadministration

SAFit2 was synthesised as before (15), encapsulated in Vesicular Phospholipid Gel (VPG) in a dose of 10mg/ml. SAFit2-VPG was provided subcutaneously under anaesthesia in a dose of 1mg/10g, providing approximately 7 days of slow release (4). SAFit2 has a terminal plasma half-life in mice of 9.7h, a rather high (15) volume of distribution, and moderate brain permeability (53).

### Studydesign

The study was divided in a series of experiments, each with different constellations of tests and duration, designed to answer specific questions (Table S1). Key outcomes, including mechanical sensitivity and weight bearing, were assessed frequently through most experiments, while catwalk, activity, anxiety- and depressive-like behavior were only assessed at selected time points, based on previous experience(14), the characterization study (Fig 1) and hypothesis of each individual experiment.

### Experimentalprocedures

*Induction of injury, (MIA);* Induction of the Monoiodoacetate Arthritis Model to the knee joint (MIA) was performed under anaesthesia as previously reported(14, 35), injecting 1mg Sodium Iodoacetate (Sigma) in 10μl saline in the left knee joint. Naïve control animals were only exposed to anesthesia.

### Behaviouraltesting

Behavioural testing was performed in randomized order by the same female experimenter, who was blinded to treatment and/or genotype.

#### Mechanical allodynia (VF)

low intensity mechanical sensitivity was assessed as previously (14), using a series of calibrated von Frey monofilaments (Ugo Basile SRL, Italy). The 50% response threshold was determined using the Dixon up-down method, using the following equation; 50% threshold (g) = 10_log(last filament)+k*0.3_, where the response pattern determined the constant, *k*(54).

#### Affective-motivational behavior (AF)

We assess the affective responses displayed after stimulation with selected filaments (low; 0.04g, medium; 0.16g, high; 1.0g as previously (14). The duration of conscious attending behavior incl. licking, biting, lifting or guarding the paw, was quantified within 30 seconds after stimulation.

#### Functional impairment - Weight bearing (WB)

static weight bearing distribution across the two hindlimbs was assessed as previously(14, 55), using the Bioseb Incapacitance Test (Bioseb), and calculated as: WB-% = (weight borne on the injured leg/weight borne on both legs) * 100%.

#### Catwalk gait analysis

analysis of voluntary movement and dynamic gait pattern was performed using the Catwalk® XT 10.0 system (Noldus Information Technology) as before (14, 42). For outcome-measures like swing-time ratio, contact area and single stance, the data was converted into a ratio between ipsi- and contra-lateral hind-limbs. The “guarding index” was calculated as previously described (56) and the higher guarding index suggests less dynamic weight bearing on the injured leg compared with the non-injured.

#### Sucrose Preference Test (SPT)

the SPT was used to measure anhedonia, as before (14). The preference for drinking water vs 1% sucrose was assessed in the home-cage environment for 12h overnight, during two nights with the sucrose bottle presented in the opposite side, to account for potential side-preference, as reported previously (42). The sucrose preference % was calculated across the two nights; SPT% = (sucrose-solution consumed / (total fluid consumption)) * 100%.

#### Elevated Plus Maze (EPM) and Open Field Test (OFT)

to assess anxiety-like behaviour, animals were assessed during a 5min EPM- or OFT-test, as before (14, 42). Recording was performed by a camera placed above the maze/arena, and movement between zones was tracked using EthoVision XT14 (Noldus Information Technology). Anxiety-like behavior was suggested by less time spent in the open arms (EPM) or center of the arena (OFT).

#### Sleep-like/activity-pattern

to measure undisturbed activity in the home-cages, we adopted the approach of Brown et al. (14, 20) using non-invasive passive infrared motion sensors to record single-housed animals every 10 seconds across multiple-day periods. As previously validated against EEG-recording, immobility defined sleep, referred to as immobility in the manuscript, was defined as periods in which no activity was measured for 40 seconds or more. Several summary statistics of circadian disruption were calculated for individual animals across the 5- to-14-day periods (depending on the experiment) and, where appropriate, on each individual day: inter-daily stability, intra-daily variability, light-phase activity, dark-phase immobility, the Lomb-Scargle periodogram, similar to the chi-square periodogram (20).

### ELISA

Plasma corticosterone levels were quantified using a commercial ELISA kit for corticosterone (Corticosterone ELISA kit, ab108821, Abcam PLC) according to the manufacturer’s instructions (4, 5).

### Sequencing

RNA was extracted from lumbar (L4-L6) spinal cord and sequencing libraries prepared using standard approaches outlined in SI Appendix. Next generation sequencing (NGS) was performed for 150 cycles on a NextSeq®500 (Illumina) to generate 75 bp paired-end reads with 19.5M coverage on average per sample. Sequencing reads were aligned to mouse GRCm38 (mm10) reference genome. RNA transcripts exceeding a 1.2-fold change in expression and Benjamini-Hochberg adjusted p-values < 0.05 were considered differentially expressed. We additionally performed exploratory analysis on transcripts with p-values exceeding nominal (p < 0.05) significance.

### CTscans

Micro-CT analyses of knee joints were performed using VECTor6CT (MILabs, The Netherlands), and analysed using BoneJ software for bone volume (BV) in region of interests (ROI). Macroscopic analysis was done using VivoQuant version 4 software (Invicro LLC, Boston, MA, USA).

### Statisticalanalysis

Statistical tests were performed in IBM SPSS Statistic Program (vers 26) or GraphPad Prism (vers 9). Interrogation of RNAseq data performed in R Studio version 4.3.3 or later. P<0.05 was considered statistically significant. Individual animals were considered as experimental unit, and sample-size was based on previous experience, based primarily on securing enough power for the assessment of depressive-like behavior across sex. All statistical tests and significant F-values are reported in SI Appendix Table S2 and S3, and key ANOVA-results are presented in the figures. Tests for normality of residuals and homogeneity of sample variances was confirmed before parametric analysis. The VF-dataset was log-transformed to ensure a normal distribution(5, 57).

## Acknowledgments

This project was funded by a Versus Arthritis grant to SG and FH; Research Award 21972. SSingleton was funded by the Advanced Pain Discovery Platform (MR/W002566/1). SNP is funded by the BBSRC (BB/X002357/1) and the NC3Rs (NC/V000977/1).

## Notes

### Competing Interest Statement

The authors have declared no competing interest.

### Summary of Updates

This version has been revised to conform with format/length requirements for a specific journal. The data behind the manuscript, the analysis and conclusions remain the same, but some elements of data and methods-description has been moved to supplementary materials. Text-sections throughout have been reworded for clarity and shortening.

