## Supplementary file for "Analgesia through FKBP51 inhibition at disease onset confers lasting relief from sensory and emotional chronic pain symptoms"

**Author Contributions:** Study conception and experimental designs: SH and SG. Data collection: All behavioural studies were done by SH; RF and SG contributed to tissue collection; SG contributed to the recordings of home cage activity. SC acquired joint CT scans. TH synthesised SAFit2. KK prepared the VPG. Data analysis and interpretation: SH and SG analysed and interpreted all data. SC contributed to CT scans analysis. SSingleton designed and ran all RNA sequencing analysis and AF contributed to analysis of activity data specifically. RF and OM provided critical insights throughout the study. Manuscript preparation: SH, SSingleton and SG wrote the initial draft of the manuscript. AF, SP provided critical comments and revisions on the draft and all authors read and approved the final manuscript.

**This PDF file includes:**

Supporting text
Figures S1 to S23
Tables S1 to S9
SI References

**Extended Methods**

***Animals and housing;***

Male and female mice (C57Bl/6J from Charles River, UK) arrived at our facility at 8 weeks of age, and were left to acclimatize for at least 7 days before experiments started. *Fkbp51* KO colony was maintained at UCL breeding +/- animals together, and experiments were started when they were 8-10 weeks old. All experiments comparing wild-type (WT) and knockout (KO) animals were performed using WT and KO littermates bred in our colony. *Fkbp5* KO and their WT littermates were previously obtained from FKBP51 heterozygous from C.A. Dickey's group (University of South Florida, USA). These mice were from mixed genetic background, C57Bl/6J and Swiss Webster. All animals were kept in groups of 2-4 in a temperature-controlled (20±1°C) environment in Individual Ventilated Cages-cages (SealSafe Plus GM500, Tecniplast, 39*19*16cm); equipped with sawdust (Lignocel Select fine), nesting material (Datesand Cocoons), wooden chew stick (LBS Small Aspen chewstick) and cardboard tunnels for shelter and handling (LBS Standard Fun tunnel); light-dark cycle of 12 hours (gradual lights on between 7-8a.m. and off between 7-8p.m; in our activity studies, ZT0=7am, ZT12=7pm); and ad libitum provision of food and water (Teklad global 2018 diet). During the activity-experiments, animals were single-housed in open wire top M1 mouse cage (North Kent Plastic Cages, 45*28*13cm) without tunnels for shelter. All experiments were carried out under the Home Office License P8F6ECC28, ARRIVE guidelines were followed, and all efforts were made to minimize animal suffering and to reduce the number of animals used (UK Animal Act, 1986).

***Genotyping***

Genotyping was performed as previously described (1, 2). DNA was extracted from a small portion of ear tissue and the following primers were used for the PCR: forward primer (for WT and 51KO): AAAGGACAATGACTACTGATGAGG; reverse WT primer: AAGGAGGGGTTCTTTTGAGG; reverse 51KO primer: GTTGCACCACAGATGAAACG. Samples from WT animals showed a single PCR product of 363bp; samples from KO animals showed a single PCR product of 510bp and samples from HET animals presented both bands.

***Compound preparation and administration:***

SAFit2 was synthesised as before (3), encapsulated in Vesicular Phospholipid Gel (VPG used at 10 mL/kg or 100 mg/kg) in a dose of 10mg/ml. VPGs (with and without SAFit2) were prepared by a dual asymmetric centrifugation (DAC) technique as described below. SAFit2-VPG was always provided subcutaneously under anaesthesia in a dose of 1mg/10g, providing approximately 7 days of slow release(1).

*Preparation of vesicular phospholipid gels by dual asymmetric centrifugation*

Vesicular phospholipid gels with a phospholipid amount of 50% (m/m) were prepared by DAC. To encapsulate the poorly water-soluble SAFit2 into the formulation by a direct incorporation method, the accurately weighted phospholipid was dissolved in ethanol (100%) and SAFit2 stock solution (20.0 mg/mL in 100% ethanol) was added in the desired amount to the phospholipid solution. The ethanol was evaporated for 2 days in a vacuum drying oven at a temperature of 25°C and a pressure of 10 mbar. Then, the solid mixture was hydrated with the accurate amount of 10 mM PBS buffer (pH 7.4) and homogenized by a DAC (Speedmixer DAC 150.1 FVZ; Hauschild GmbH &Co. KG, Hamm, Germany).

Dual asymmetric centrifugation (DAC) (a double centrifugation technique) at 3500rpm was used for homogenization of the phospholipids and buffer, as previously described by Massing et al. in 2008(4). A custom-made cooling system was installed to prevent the material from heating during the continuous mixing over 45 minutes. The final amount of SAFit2 in the formulation was 10 mg/g. Then, the solid mixture was hydrated with the accurate amount of 10 mM PBS buffer pH 7.4 and homogenized by a DAC (Speedmixer DAC 150.1 FVZ; Hauschild GmbH &Co. KG, Hamm, Germany).

The described manufacture process was used for the preparation of a second VPG formulation without the addition of SAFit2 stock solution as a control formulation (Vehicle).

***Study design;***

The study was divided in a series of experiments, each with different constellations of tests and duration, designed to answer specific questions (Table S1). All behavioral experiments generally included assessment of most behavioral outcomes in the same set of animals, except the activity assessment, which was recorded in a separate set of animals, due to the need for single-housing during these experiments. The activity studies were divided into two cohorts, including equal numbers of animals from each group. Similarly, the characterization-studies consisted of two cohorts tested for 3 or 6 months respectively, including slightly different combinations of behavioral assessment. These independent cohorts consistently yielded reproducible results on overlapping measures (*i.e.* sucrose preference and mechanical sensitivity), and results were combined. Results from the *Fkbp5* KO/WT mice were obtained from littermates from at least 3 breeding pairs.

**Table S1: Study design.**

| **Experiment purpose** | **Groups** | **Tests** | **Figures** | **Total N** (note) |
|---|---|---|---|---|
| Characterization of MIA-induced changes (3-6 months) | MIA vs control (male vs female) | WB, VF, affective-VF, SPT*, OFT, | Fig 1 + S1 | 22 + 23 (lost: 3) (across 2 cohorts) |
| | | Activity, BV | Fig 2 + S2-4, Fig 1C | 32 (across 2 cohorts) |
| Effects of *Fkbp5* genetic manipulation on MIA-induced changes (6 months) | KO vs WT (male vs female) All MIA-injured | WB, VF, BV, SPT, OFT, EPM, BW | Fig 3 + S5 | 22 (littermates from 3 breeding pairs at least) |
| | | Activity, (SPT) | Fig S6-9 (+3D) | 16 (across 2 cohorts) |
| Effects of FKPB51 inhibition after MIA-injury (3 weeks, 2 months, 4-5months) | SAFit2-VPG vs VPG/vehicle, (Only males or male vs female) All MIA-injured | WB, VF, SPT, OFT, EPM | Fig 4 + S10 | S10.A-B.: 15 (lost:1)<br>4A-B: 24<br>4C-G + S10C-H: 22 (lost: 2) |
| Effects of FKBP51 inhibition in naïve animals (4 weeks) | SAFit2-VPG vs VPG/vehicle, (male vs female) All naïve | OFT, VF, SPT 1%, SPT 0.5% | Fig S11 | 16 |
| | | Activity | S15 | 16 (across 2 cohorts) |
| Effects of FKPB51 inhibition at MIA-injury onset. (6months) | SAFit2-VPG vs VPG/vehicle, (male vs female) All MIA-injured | WB, VF, BV, affective-VF, SPT, OFT, BW, Brush | Fig 5A-G + S12-13 | 24 |
| | | Activity, catwalk | Fig 5H-L + S14 | 20 (across 2 cohorts) |
| RNA sequencing of early vs late treatment (30 days) | SAFit2-VPG vs VPG/vehicle, (male only) All MIA-injured | VF, catwalk, corticosterone, RNA sequencing. | Fig 6 + S16-21 | 30 (lost: 2) |

Abbreviations: VF = von Frey. WB = weight bearing. OFT = Open Field Test. SPT = Sucrose Preference Test. EPM = Elevated Plus Maze. BV = Bone Volume. BW = Body Weight. MIA = monoiodoacetate induced arthritis. VPG = Vesicular Phospholipid Gel. *one cage of 4 male mice were particularly aggressive, and were therefore not exposed to SPT until the last measure (6months), as the protocol of separation overnight and re-integration was associated with a risk of increased fighting.

Animals were allocated into groups using block randomisation, securing equal representation of treatment-groups in each cage, to minimise potential confounders related to housing/cage. Overall, each study included equal numbers of animals in each group, but due to the toxic nature of the MIA-chemical, a few animals were lost during model-induction in some experiments. Equally, due to the long-term nature of the experiments, some animals were lost at different stages of the experiment due to other accidental losses not related to the experiment, like fighting wounds or dermatitis. Additionally, one cage was excluded from SPT for certain periods due to aggressive behaviour in the cage, which could be worsened by the protocol, and two animals were excluded from the weightbearing (WB)-assessments in Fig 4B and 4D before treatment-initiation, as they were not compliant in the WB-assessment. Besides these few exceptions, no animals were excluded from experiments or the following analysis. Additionally, for the characterisation-study, cohorts of different durations (3 vs 6months) were combined. Due to these variations, group sizes (N-numbers) were not always equal throughout the experiment, and at times defined in a range (full overview of numbers in each study, see **Table S2+3**). A total of 334 mice were used for these studies.

Key outcomes, like mechanical sensitivity and weight bearing, was assessed frequently through most experiments, while assessment of catwalk, anxiety- and depressive-like behavior were only assessed on selected timepoints, based on previous experience(5), the characterization study (Fig 1) and hypothesis of each individual experiment.

***Experimental procedures;***

*Induction of injury, (MIA);* Induction of the Monoiodoacetate Arthritis Model to the knee joint (MIA) was performed as previously reported(5, 6). On the day of injection, the solution was freshly prepared in sterile saline for injection of 1mg Sodium Iodoacetate (≥98%, Sigma) in 10µl saline (9% NaCl). Animals were anaesthetized in an induction chamber using Isoflurane 2.5% mixed in $O_2$ at a flowrate of 1.5 L/min, and maintained via facemask at 2.0% during the injection. Anaesthetic dept was confirmed by lack of withdrawal reflex to a pinch to the tail. The animal was then placed on the back in dorsal recumbency, and the fur was shaved in the area around the knee on the left hindleg. The knee was stabilized and fixed in a slightly bent position and the patellar tendon was visualized as a white line below the skin. The injection of 10µl using a 30G insulin-syringe (BD Micro-Fine Plus Demi, 0.3ml (30G) 8mm) was made intraarticularly in the joint space by applying it perpendicularly through the tendon just below the patella, and with as minimal movement as possible. The injection volume was decreased 1-2µl in smaller subjects weighing less than 22g, to account for the smaller joint size and decrease the risk of toxicity when surpassing the maximum volume-capacity of the joint space.

*Control animals.* Naïve control animals were only exposed to anesthesia.

***Behavioural testing;***

Behavioural testing was always performed in randomized order and by the same female experimenter, who was blinded to treatment and/or genotype. Animals were always allowed at least 30min of habituation to the testing room prior to behavioural testing. Unless otherwise specified, behavioral tests were always performed between 8am and 2pm.

*Mechanical allodynia (VF);* Low intensity mechanical sensitivity was assessed similar to previously reported in our group(5), using a series of calibrated von Frey monofilaments (0.02; 0.04; 0.07; 0.16; 0.4; 0.6; 1.0) (Ugo Basile SRL, Italy). Animals were placed in Plexiglass chambers, located on an elevated wire grid, and allowed to habituate to the testing environment for at least 60 min prior testing. Once the animals were calm, the plantar surface of the paw was stimulated, starting with a 0.6g filament applied with uniform pressure for 5 seconds. A brisk withdrawal, stretching or licking of toes, was considered as a positive response, whereupon the next lower-force filament was applied. In the absence of a positive response, the next higher-force filament was applied in the next test. After the first change in response-pattern, suggesting the approximate threshold, additional 4 stimulations were applied; applying the next higher-force filament when no response, and lower-force filament following positive responses. The response pattern determined the constant, *k*(7), and the 50% response threshold was determined using the following equation; 50% threshold (g) = $10^{\log(\text{last filament})+k*0.3}$.

*Affective-motivational behavior (AF);* in addition to assessing pure reflexive sensory threshold to mechanical stimulation (VF), we also adopted and modified the protocol described(5, 8) to assess the affective responses displayed after stimulation with selected filaments (low; 0.04g, medium; 0.16g, high; 1.0g). The assessment was always performed following the VF-assessment, while the animals were still in the Plexiglas chamber, and following a period of rest. Starting with the low-intensity filament, each filament was applied once for 1sec, and the duration of affective response was recorded as the amount of time, the animal showed conscious attending behavior to the stimulated paw, by licking, biting, lifting or guarding the paw, within 30 seconds after application of the filament. The animal was left to rest for at least 10min before the next higher filament force was applied.

*Functional impairment – Static Weight bearing (WB);* To assess the functional impact of the injury the weight bearing distribution was assessed similarly to previously described (5, 9). Hindlimb weight bearing was measured using a Bioseb Incapacitance Test (Bioseb) which measures the weight distribution across the two hindlimbs of a stationary animal. Animals were habituated and trained to become comfortable with the testing paradigm for 5 days prior to induction of the model. Three readings were collected and averaged for each animal, and the weight borne by the ipsilateral limb was expressed as a percentage of the weight borne across both hindlimbs (WB-% = (weight borne on the injured leg/weight borne on both legs) * 100%).

*Catwalk gait analysis:* Analysis of voluntary movement and gait pattern was performed using the Catwalk® XT 10.0 system (Noldus Information Technology) and based on our previous experience (5, 10). Briefly, green light was internally reflected into a glass plate, on which an enclosed corridor was fixed, with red backlight above the corridor. A video-camera was mounted underneath the setup and recorded the paw prints being lit up by the green light when paws were in contact with the glass plate as the mouse walked along the corridor. A run was regarded as compliant when the animal entered in one end of the corridor, and moved fluently across the plate towards the other end of the corridor, with a running duration below 12 seconds and a maximum variation below 75%. Three compliant runs were recorded for each animal, with no previous training/habituation. For outcome-measures like swing-time ratio, contact area and single stance, the data was converted into a ratio between ipsi- and contra-lateral hind-limbs. The parameter "guarding index", was calculated as previously described(11) and the higher guarding index suggests less dynamic weight bearing on the injured leg compared with the non-injured. Therefore outcomes from the catwalk analysis are for simplicity at times referred to as "dynamic weight bearing" in the manuscript.

*Sucrose Preference Test (SPT);* To assess depressive-like behaviour, the sucrose preference test was included as a measure of anhedonia, as previously described (5). At least 5 days before SPT each home-cage was fitted with two water bottles, to acclimatise the mice to drinking from both bottles / sides of the cage. Two days before the SPT test, one of the water-bottles was filled with 1% sucrose solution for approximately 24h, swapping side halfway, to acclimatise the mice to the sweet solution, and learn that it may be presented in both sides. No sucrose was provided the last 24 hours before the actual test. For the SPT test, all animals were individually housed in clean cages (fitted with the same materials and enrichment as when group-housed) in their home-cage environment for 12h overnight, and were given free access to two pre-weighed bottles containing either normal drinking water or 1% sucrose solution. In the morning all bottles were weighed, and mice were placed back together with their cage-mates. This re-introduction released some fighting in especially some male cages, and mice were given a night to calm down before the test was repeated overnight with the sucrose bottle presented in the opposite side, to account for potential side-preference, as reported previously in rats(10). No side-preference was detected in the current study, and the sucrose preference % was calculated across the two nights, using the following calculation; SPT% = (sucrose-solution consumed / (total fluid consumption)) * 100%. The sucrose-solution was always prepared fresh before provision at 1% in the normal drinking water, although for one experiment assessing effects of treatment in naive animals (Fig S11) a lower 0.5% solution was also included. As recent meta-analysis suggests food- and water-deprivation to confound the outcome of this assay(12) no water- or food-deprivation was employed in this study.

*Elevated Plus Maze (EPM);* To assess anxiety-like behaviour an EPM was used as described previously(5, 10). The maze consisted of four arms (35*5 cm) arranged in a cross-like disposition, and 60cm above ground (Ugo Basile SRL, Italy). Two opposite arms were open, and the other two were equipped with 15 cm high walls on each side for enclosement. All were connected by a central 5*5 cm square. The animal was placed in the centre, for free exploration for 5 minutes. Recording was performed by a camera placed above the maze, and movement between zones was tracked using EthoVision XT14 (Noldus Information Technology). The maze was wiped with 70% ethanol between each animal to minimize the influence of odours between animals.

*Open Field Test (OFT):* to assess anxiety-like and locomotor activity, an OFT was used, as previously described (5). The OFT was a circular open arena with grey plastic flooring and blue plastic sides (diameter, 36cm, height 32cm). The arena was evenly illuminated by lighting placed above the arena. A video camera was positioned directly above the arena and connected to a computer performing live-tracking and recording of the behavior using Ethovision XT14. The animal was placed in the middle of the arena and allowed 5 minutes of free exploration. The proportion of time spent in the centre vs by the edges/walls of the arena, was used as a marker for the anxiety-like behavior (outer zone; 6.7cm along the edge of the arena. Centre zone diameter; 22.6cm).

*Sleep-like/activity pattern;* To measure undisturbed activity in the home-cages, we adopted the approach of Brown et al. (5, 13) using non-invasive passive infrared motion sensors. Animals were single-housed with a 12h light:12h dark cycle. The cages were open with wire tops (M1 mouse cage, North Kent Plastic Cages, floor area: 500cm$^2$) and a passive infrared (PIR) motion sensor was fitted above the cage. For accurate measurements of activity, the area below the food- and water-hopper was blocked off and tunnels / shelters were removed, allowing the animal to be in the sensors receptive field at all times. Home cage mouse activity was tracked as in Brown et al. (13), with measurements taken every 10 seconds across multi-day periods. As previously validated against EEG-recording, immobility defined sleep, referred to as “immobility” in the manuscript, was defined as periods in which no activity was measured for 40 seconds or more. Activity and immobility data were smoothed by calculating the mean in 10-minute bins as preliminary experiments demonstrated that this provided a balance between reducing measurement noise and maintaining time series features. Several summary statistics of circadian disruption were calculated for individual animals across the 5-to-14-day periods (depending on the experiment) and, where appropriate, on each individual day: inter-daily stability, intra-daily variability, light-phase activity, dark-phase immobility, the Lomb-Scargle periodogram, similar to the chi-square periodogram (14). In all figures, ZT0=7am, ZT12=7pm, with ZT = zeitgeber time, common used in circadian biology.

### ***5.2.4. ELISA***

Corticosterone levels were measured as before (1, 2). Immediately upon cervical decapitation, blood was collected in tubes containing trisodium citrate (Paediatric Tube 1.3ml Trisodium citrate 3.2% (9NC), Greiner Bio-One, LTD), gently mixed via inversion and stored on ice. Tubes were centrifuged at 10.000RPM at 4°C for 10min, plasma was collected and stored at -20°C until analysis. Plasma corticosterone was quantified using a commercial ELISA kit for corticosterone (Corticosterone ELISA kit, ab108821, Abcam PLC) according to the manufacturer’s instructions.

### ***5.2.5. Sequencing***

For the tissue collection, spinal cords were separated into ipsilateral and contralateral quadrants to site of injury, snap frozen on dry ice, and maintained at -80°C until analysis. RNA was extracted from spinal cord lumbar (L4 to L6) quadrants using the Qiagen AllPrep DNA/RNA/miRNA kit as per manufacturer’s instructions. RNA was quantified using a Nanodrop 8000 spectrophotometer and RNA integrity was measured using the Agilent 2100 Bioanalyser. All samples passed Quality Control (QC) with RIN scores >7. mRNA libraries were prepared from total RNA using NEBNext Ultra II Directional Library Preparation Kit. Fragmentation of isolated mRNA prior to first strand cDNA synthesis was carried out using incubation conditions recommended by the manufacturer for an insert size of 300bp (94°C for 10 minutes). 13 cycles of PCR were performed for final library amplification. Resulting libraries were quantified using the Qubit 2.0 spectrophotometer and average fragment size assessed using the Agilent 2200 Tapestation. A final sequencing pool was created using equimolar quantities of each sample library. 75bp paired-end reads were generated for each library using the Illumina NextSeq®500 in conjunction with the NextSeq®500 v2 High-output 150-cycle kit to obtain an average of 19.5M read pairs per sample.

Raw paired-end next generation sequencing reads (Illumina HiSeq 2500) were assessed for quality scores and trimmed using a sliding window operation with an average quality score Q > 20. Illumina adapter sequences were also removed. Reference-based alignment was performed using the mouse mm10 genome assembly and successfully aligned features quantified at the gene-level using featureCounts. Summarised gene counts were transformed into transcripts per million (TPM) to account for biases arising from differing gene lengths(15) and library sizes normalised using trimmed mean of M-values. Gene matrices were filtered by expression using a minimum count > 10 in at least 70% of samples and a minimum overall count > 15 transcripts to remove genes that were either not expressed or lowly expressed among all samples. Differential expression among treatments was established using linear modelling through limma using log2-transformed TPM. Heteroscedasticity of the mean-variance relationship in the latter was removed using limma voom. Genes exceeding a 1.2-fold change in expression and Benjamini-Hochberg (FDR) adjusted p values < 0.05 were considered differentially expressed among groups. When stated explicitly, we additionally performed exploratory analysis on genes with p values exceeding nominal (p < 0.05) significance only. Gene overrepresentation tests and gene ontology of significant gene sets were performed using a minimum term size > 2 against the mm10 array background gene set as the reference. Statistical comparisons against the observed *vs* expected gene set ratio were assessed using Fisher's exact tests and corrected using FDR at P < 0.05. All annotations were identified using the GO biological process complete database available through PANTHER. GO lists reduced into hierarchical semantically similar terms using ReViGo for plotting. Interrogation of RNA seq data performed in R Studio version 4.3.3 or later.

Raw fastq files are available in addition to the processed log cpm for each sample (GEO accession number: GSE293520).

***5.2.6. CT scans;***

Following dissection and fixing in 4% paraformaldehyde for 5 days, knee joints were stored in 70% ethanol. Micro-CT analyses were performed using VECTor6CT (MILabs, The Netherlands). Knee joints were imaged using single mouse bed with the following settings: ultra-focus magnification, accurate scan mode, settings default, for a total duration of 6min per CT position. Reconstruction was done using 20voxel size and images were analysed using BoneJ software for bone volume (BV) in region of interests (ROI) of the knee joint. Macroscopic analysis of joints was done using VivoQuant version 4 software (Invicro LLC, Boston, MA, USA).

***5.2.7. RTqPCR:***

For the tissue collection, cords of MIA injected mice were separated into ipsilateral and contralateral quadrants to site of injury for processing, snap frozen on dry ice, and maintained at -80°C until analysis. Total RNA was extracted using an acid phenol extraction method (TRIzol reagent; Qiagen), which involved homogenisation by hand and passage through a biopolymer-shredder (QIAshredder; Qiagen). The RNeasy mini kit (Qiagen) was used to complete the extraction and RNA concentrations were measured using a Nanodrop. 500ng of total RNA was then reverse transcribed to cDNA via a two-step method. First, samples were incubated with random nonamers (2uM; Sigma), oligo dT20 primer mix (1uM; Promega) and 10mM dNTP mix (0.5uM; Promega) at 65°C for 5 minutes. First-strand buffer (5x) and superscript III were then added, along with DTT (0.1M; all Promega), RNaseIN recombinant ribonuclease inhibitor (Promega) and completed by exposure to a heat cycle of 25°C for 5 minutes,

50°C for 50 minutes and 70°C for 15 minutes. Appropriate positive and negative controls were included in the process. cDNA was stored at -20°C until further processing. RT-qPCR reactions were run using the DNA Engine and the SYBR Green JumpStart Taq Ready Mix (Sigma), in the standard 3-step SYBR green heat cycle. Reactions were run in triplicates and assessed by melting curve analysis. The ratio of the relative expression of target genes to HGPRT expression was calculated using the 2ΔCt formula.

### ***5.3. Statistical analysis;***

The experimenter was blinded to genotype and treatment groups until the time of data-analysis. All statistical tests were performed in IBM SPSS Statistic Program (vers 26) or GraphPad Prism (vers 9). Interrogation of RNA seq data performed in R Studio version 4.3.3 or later. P<0.05 was considered statistically significant. Individual animals were considered as experimental unit, and sample-size was based on previous experience, based primarily on securing enough power for the assessment of depressive-like behavior across sex. All statistical tests and significant F-values are reported in supplementary table S2 and S3. Tests for normality of residuals and homogeneity of sample variances confirmed before parametric analysis. Post-hoc analysis was only performed on statistically significant F values. The VF-dataset was log-transformed to ensure a normal distribution, as the von Frey hairs are distributed on an exponential scale. This was similar to previous studies in our group(2), and as demonstrated to make more mathematical and biological sense(16). Most long-term timeline datasets were also converted into weighted average (AVG), either for the entire duration of the experiment, or for selected relevant phases of the experiment, like when a compound was active or the first 2 weeks after model induction.

***Study approval:*** All experiments were carried out under the Home Office License P8F6ECC28, ARRIVE guidelines were followed, and all efforts were made to minimize animal suffering and to reduce the number of animals used (UK Animal Act, 1986)

**Supporting figures:**

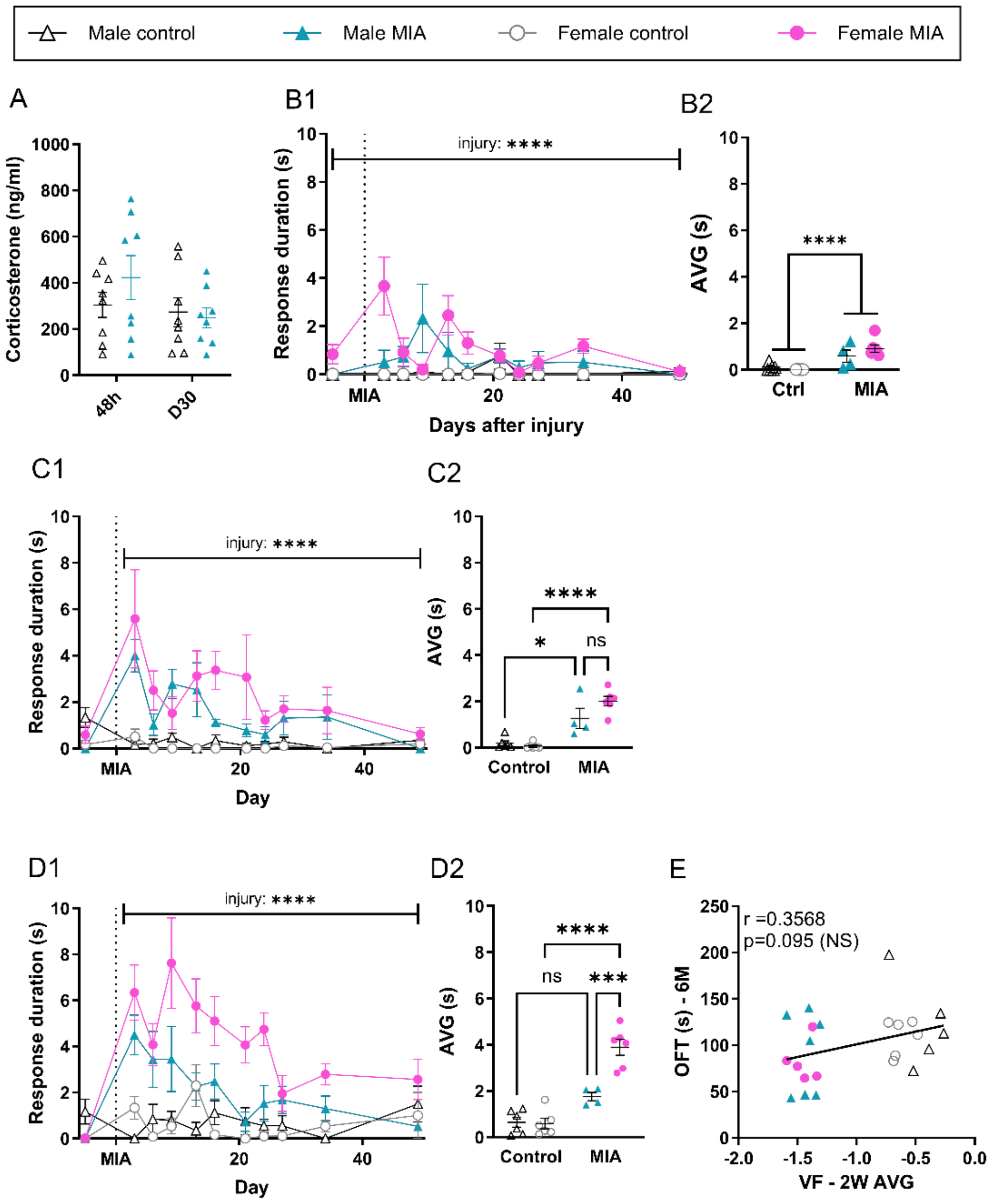

**Figure S1. MIA induces changes in affective behavior.** Male and female mice were exposed to MIA or control and followed for up to 49 days after injury. (**A**) Corticosterone was measured at 48 hours and 30 days after MIA-injury in male mice (N=8). (**B1,2**) The duration of the affective response following the application of a low force (0.04g) VF filament was recorded in seconds and followed for 49 days after injury (N=4-6). (**C1,2**) The duration of the affective response following the application of a medium force (0.16g) VF filament was recorded in seconds and followed for 49 days after injury (N=4-6). (**D1,2**) The duration of the affective response following the application of a high force (1.0g) VF filament was recorded in seconds and followed for 49 days after injury (N=4-6). (**E**) Correlations between early (2-week average) mechanical sensitivity and the later development of anxiety-like behavior at 6 months. B2, C2, D2: Weighted average (AVG) was calculated across the full study period. Data shows mean ± S.E.M. NS= Not significant, *P<0.05, **P<0.01, ***P<0.001, ****P<0.0001, as determined using ANOVA ANALYSIS for B1, C1 and D1, and appropriate post-tests. See full statistical analysis in Supplementary Table S3.

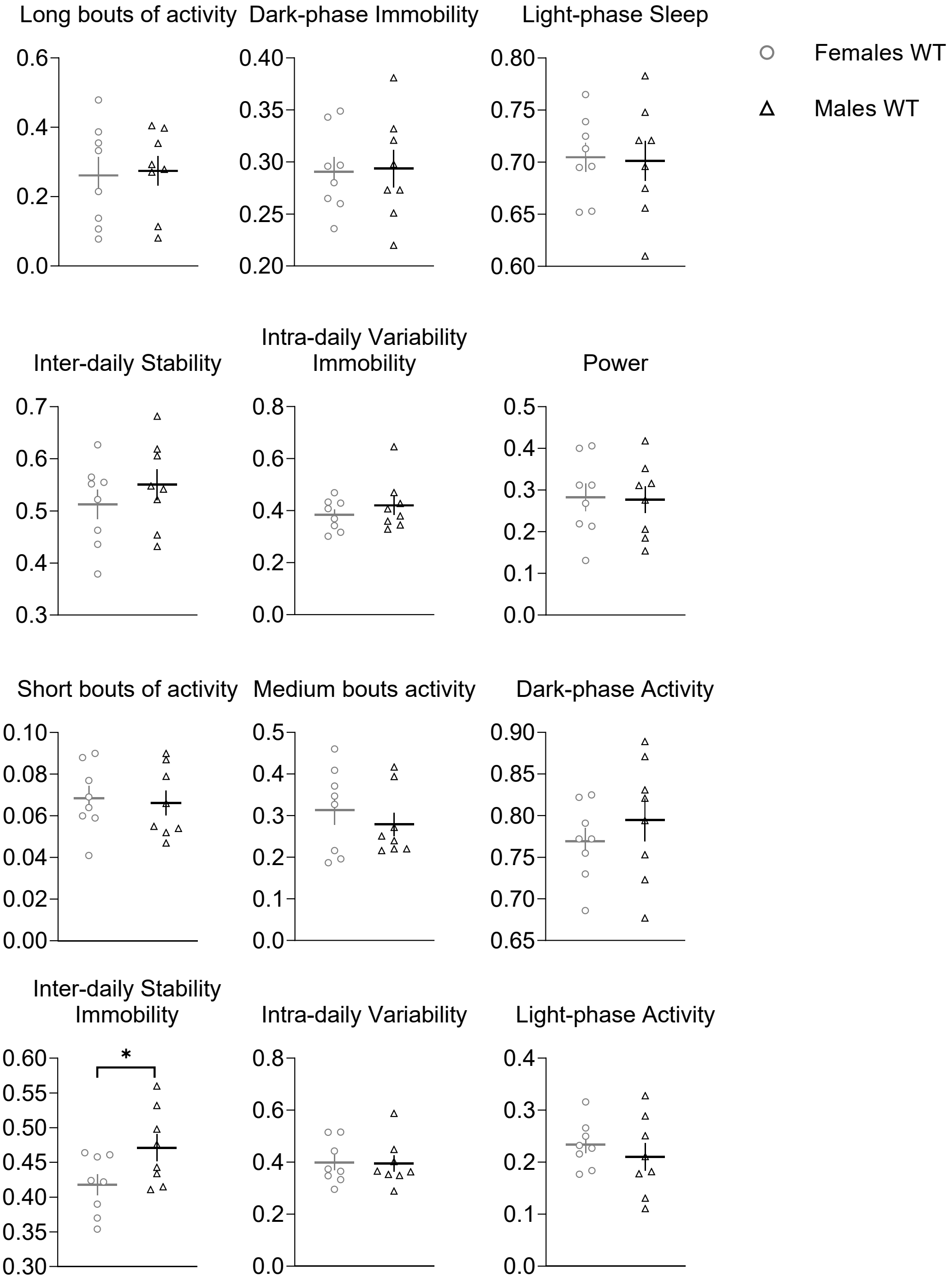

**Figure S2. There are no obvious differences in daily activity rhythms between 8-week-old male and female naïve mice.** Various summary statistics of circadian disruption calculated across a 6 day-period. N=8/8. *P<0.05. Data shows mean ± S.E.M. and single data points. See full statistical analysis in Supplementary Table S3.

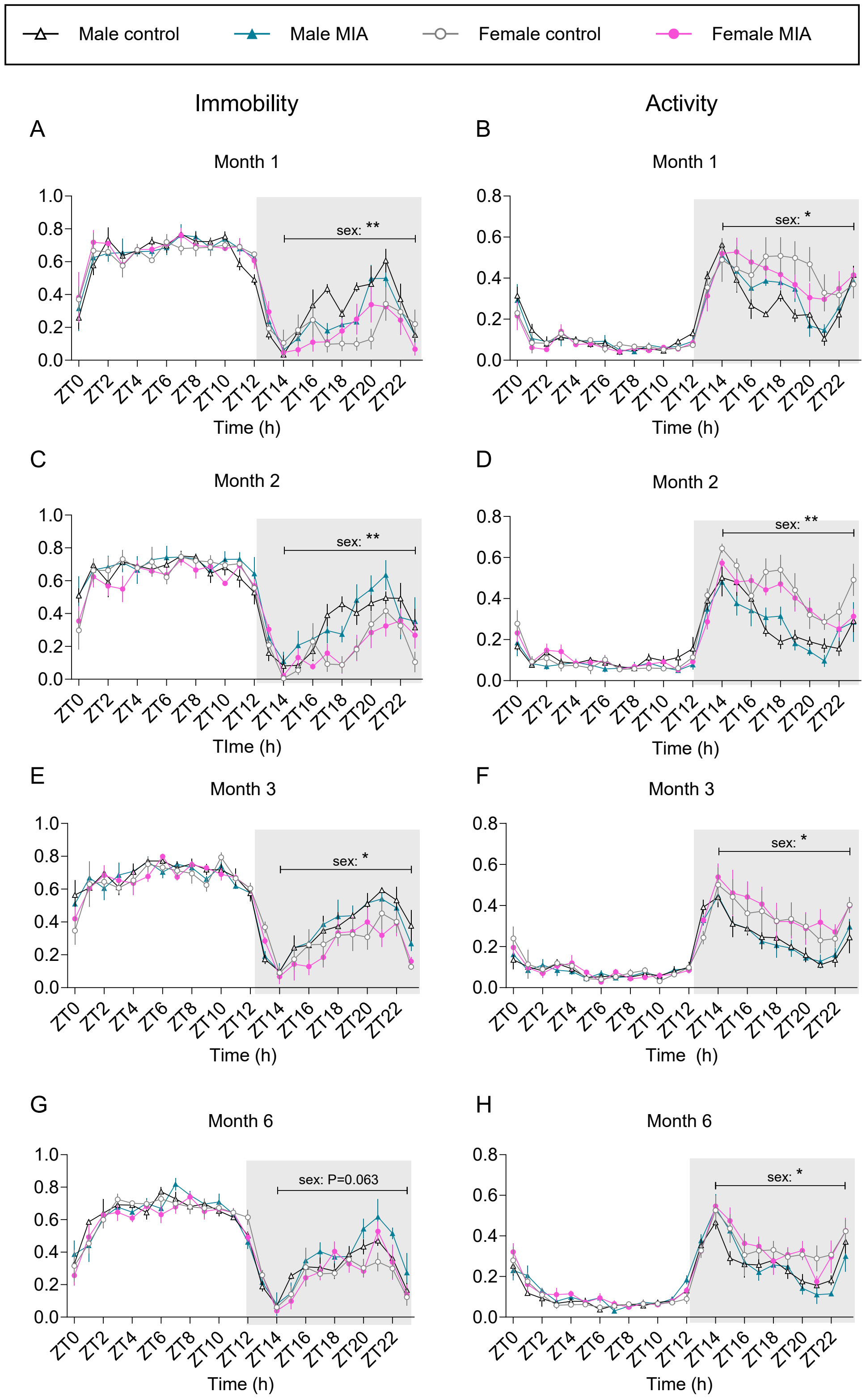

**Figure S3. From one month after MIA injection, sex differences are obvious but there are no injury-induced changes to daily activity rhythms in male and female mice.** (**A-H**) 24h immobility/activity plots using 1h bins. (**A, B**) Average of 24h plots across 7 days, 1 month after MIA injection. N=4/4/4/4. (**C, D**) Average of 24h plots across 7 days, 2 months after MIA injection. N=4/4/4/4. (**E, F**) Average of 24h plots across 7 days, 3 months after MIA injection. N=4/4/4/4. (**G, H**) Average of 24h plots across 7 days, 6 months after MIA injection. N=4/4/4/4. ZT0=7am. Data shows mean ± S.E.M. *P<0.05; ** P<0.01, sex effect. See full statistical analysis in Supplementary Table S3.

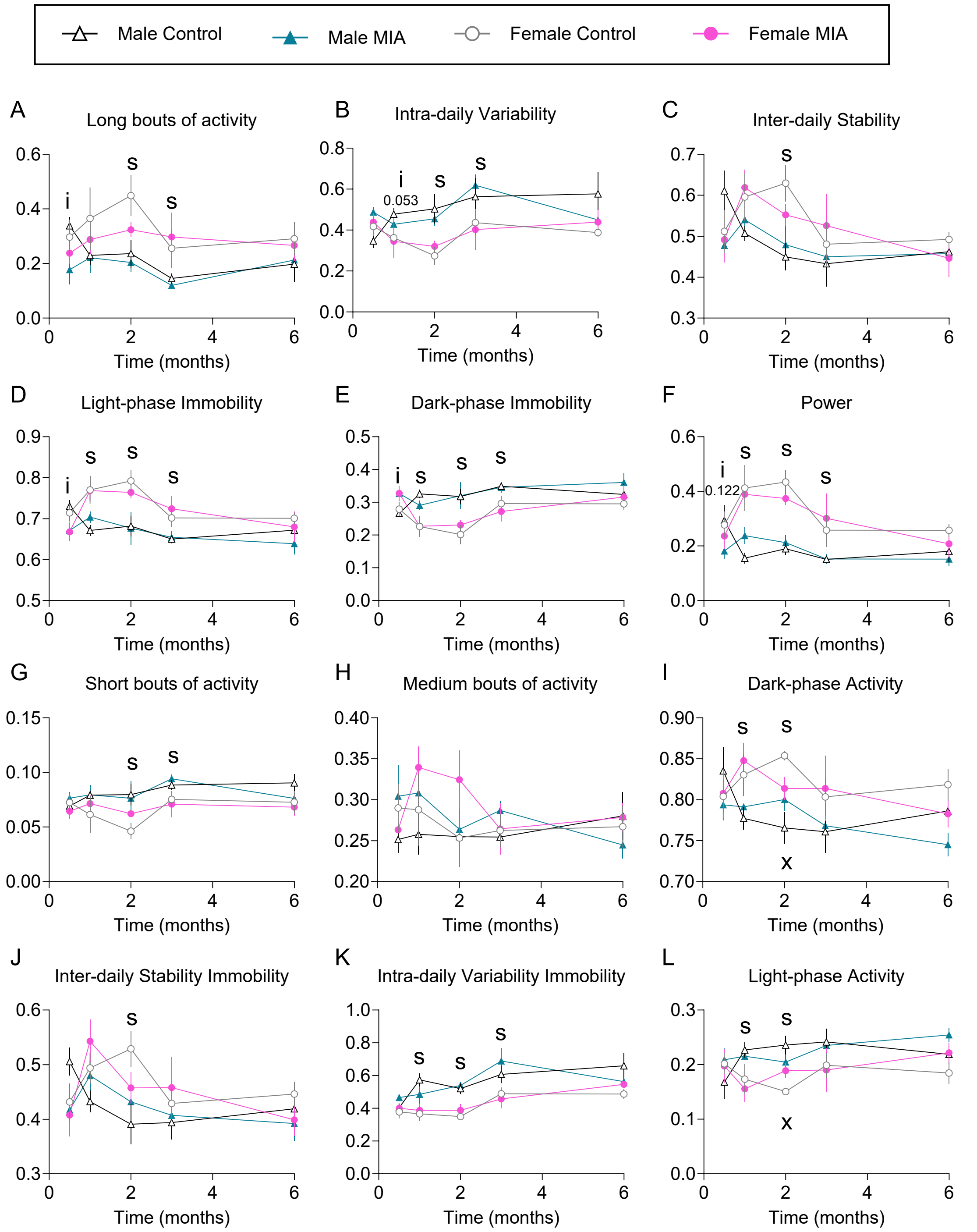

**Figure S4. From one month after MIA injection, summary statistics of daily activity rhythms indicate no impact of MIA injection but obvious sex differences.** Various summary statistics of circadian disruption calculated across a 6 day-period at each time point. Data plotted as time course for ease, but not all animals were followed for the whole time course. N=4/4/4/4 at each time point. Data shows mean ± S.E.M. s: sex effect, P<0.005; i: injury effect, P<0.05; x: injury x sex interactions. s: sex effect P<0.05; i: injury effect P<0.05; x: sex x injury P<0.05. See full statistical analysis in Supplementary Table S3.

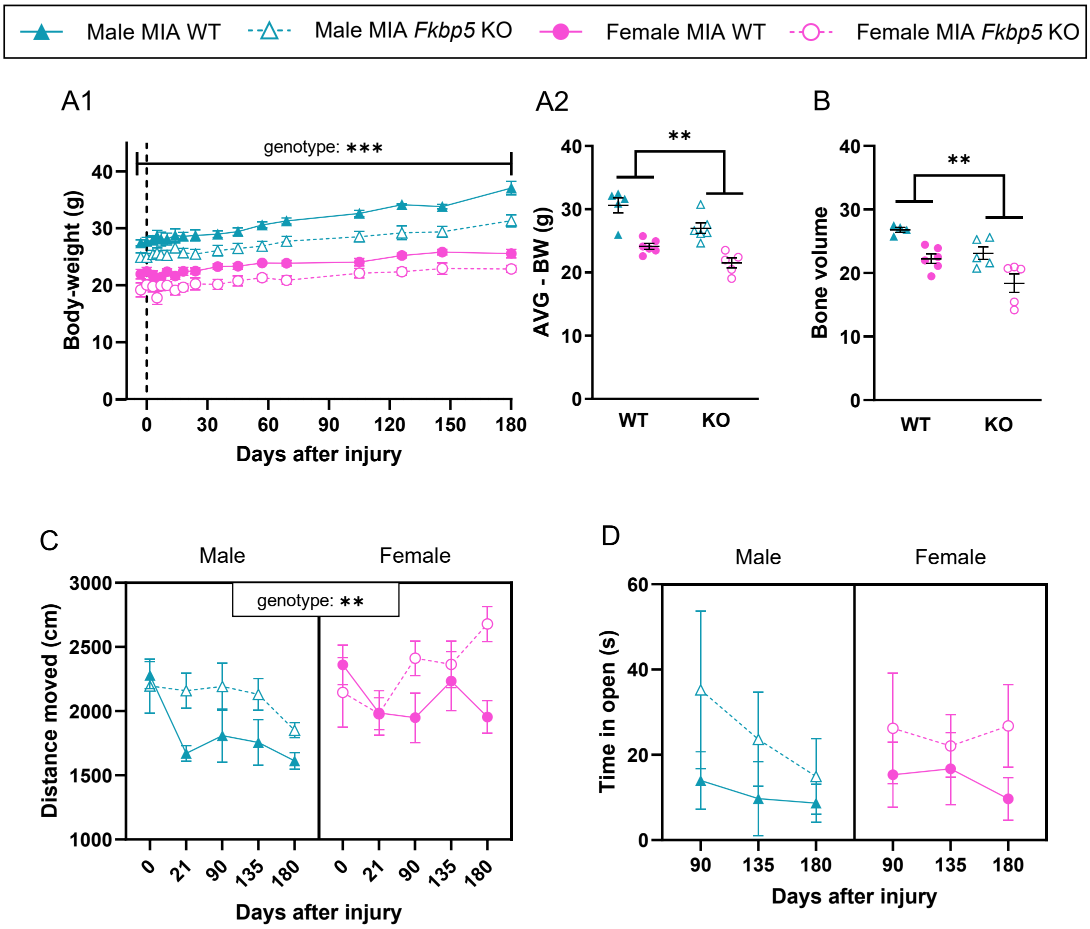

**Figure S5. *Fkbp5* global knockdown has an impact on body weight and on a range of outcomes measured after MIA injection.** Male and female *Fkbp5* global knockout and wild type littermates were all exposed to MIA injury and followed for 6 months after injury. (**A**) Bodyweight was significantly lower in KO animals. (N=5-6). (**B**) Bone volume, as assessed using microCT at 6 months after injury (N=4-6). (**C**) Distance moved in the open field test (N=5-6). (**D**) Anxiety-like behavior was also assessed using the Elevated Plus Maze, assessing the time in the open parts of the maze (N=4-6). For A2: Weighted average (AVG) was calculated for the full study period. Data shows mean ± S.E.M. *P<0.05, **P<0.01, ***P<0.001, ****P<0.0001, as determined by the ANOVA analysis. See full statistical analysis in Supplementary Table S3.

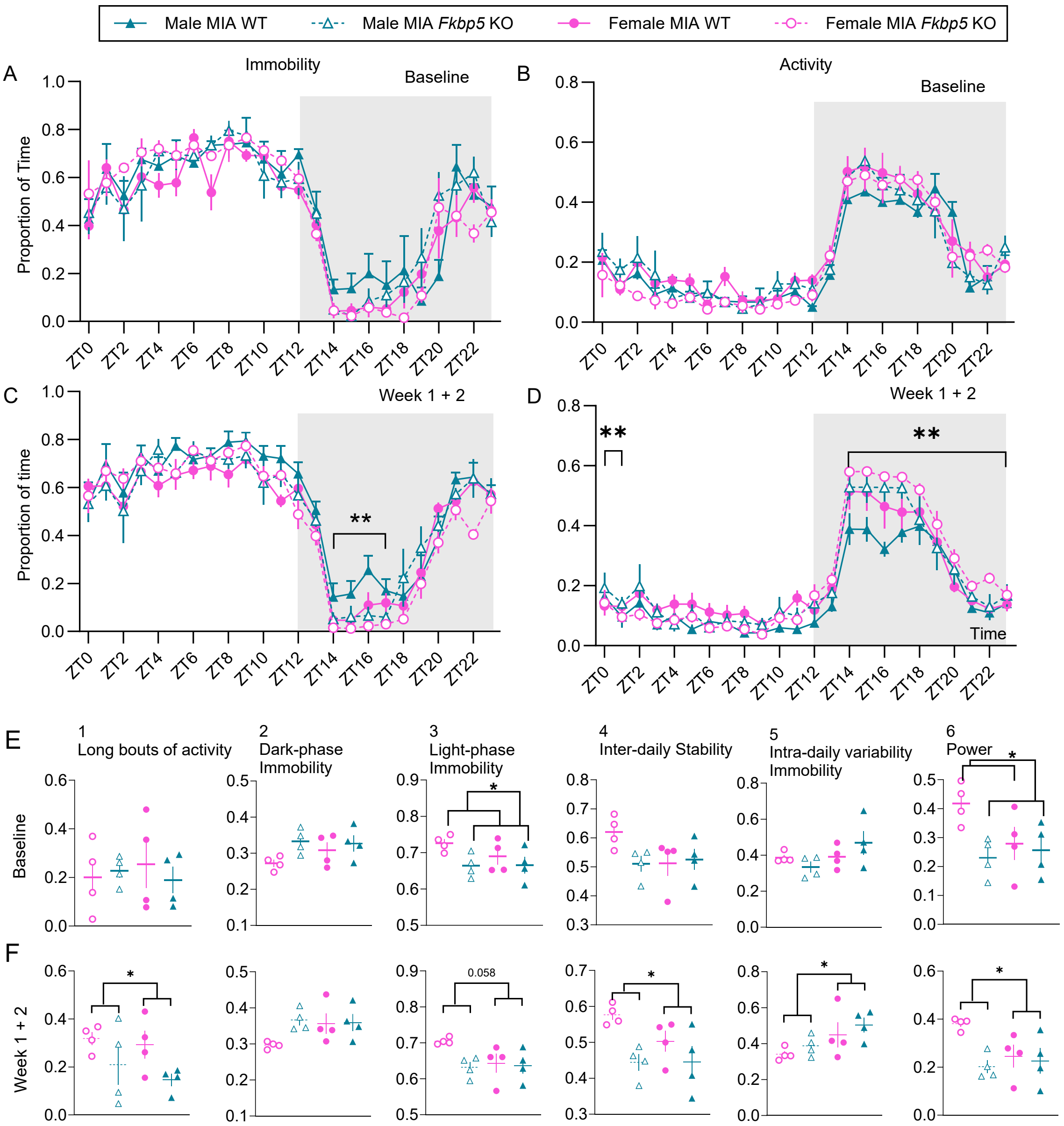

**Figure S6: *Fkbp5* global knockdown improves some of the MIA-induced changes to daily activity rhythms. (A-D)** 24h immobility/activity plots using 1h bins. **(A, B)** Average of 24h plots across the 6 days before MIA injection. N=8/8. There were no differences in immobility (A) and activity (B) patterns in naive, 8-week-old male and female, WT and KO mice. **(C, D)** Average of 24h plots across the first 2 weeks after MIA injection. N=4/4/4/4. *Fkbp5* knock-down improved some of the MIA induced changes to immobility and activity patterns. **(E,F) (1)** Proportion of time spent in long bouts (>10min) activity during the dark period, **(2)** proportion of time spent immobile during the dark phase, **(3)** proportion of time spent immobile during the light phase, **(4)** interdaily stability, **(5)**, intra-daily variability immobility and **(6)** A power of the 24-hour activity cycle . (E) Average across 6 days before MIA. (F) Average across first 2 weeks after MIA. N=4/4/4/4. ZT0=7am, ZT12=7pm. Data shows mean ± S.E.M. *P<0.05; ** P<0.01, genotype effect. (E6) sex effect plotted. See full statistical analysis in Supplementary Table S3.

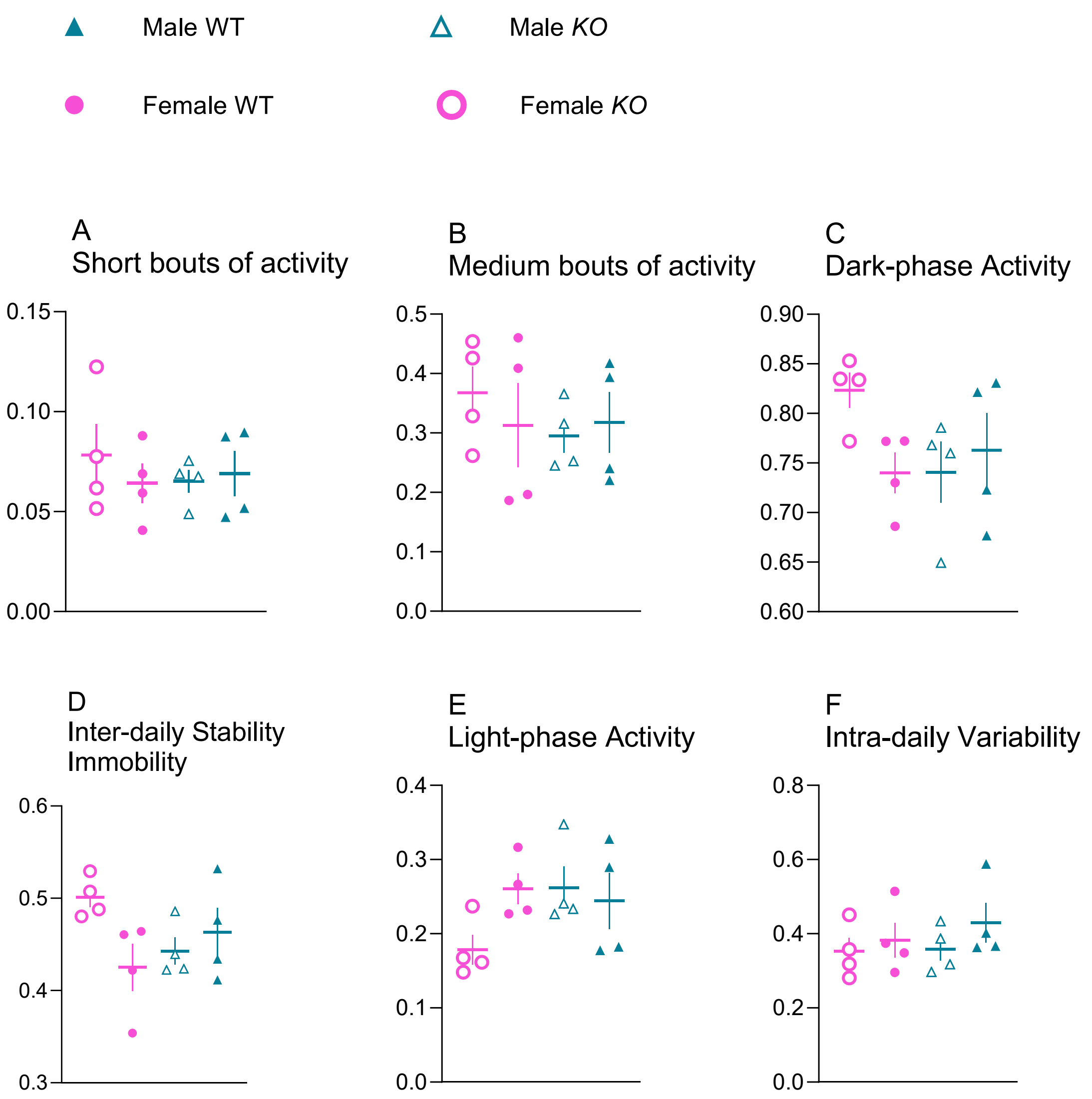

**Figure S7. There are no obvious differences in daily activity rhythms between 8-week-old male and female naïve *Fkbp5* KO and WT mice.** Various summary statistics of circadian disruption calculated across a 6 day-period pre-MIA injection. Data shows mean ± S.E.M. and single data points. N=4/4/4/4. See full statistical analysis in Supplementary Table S3.

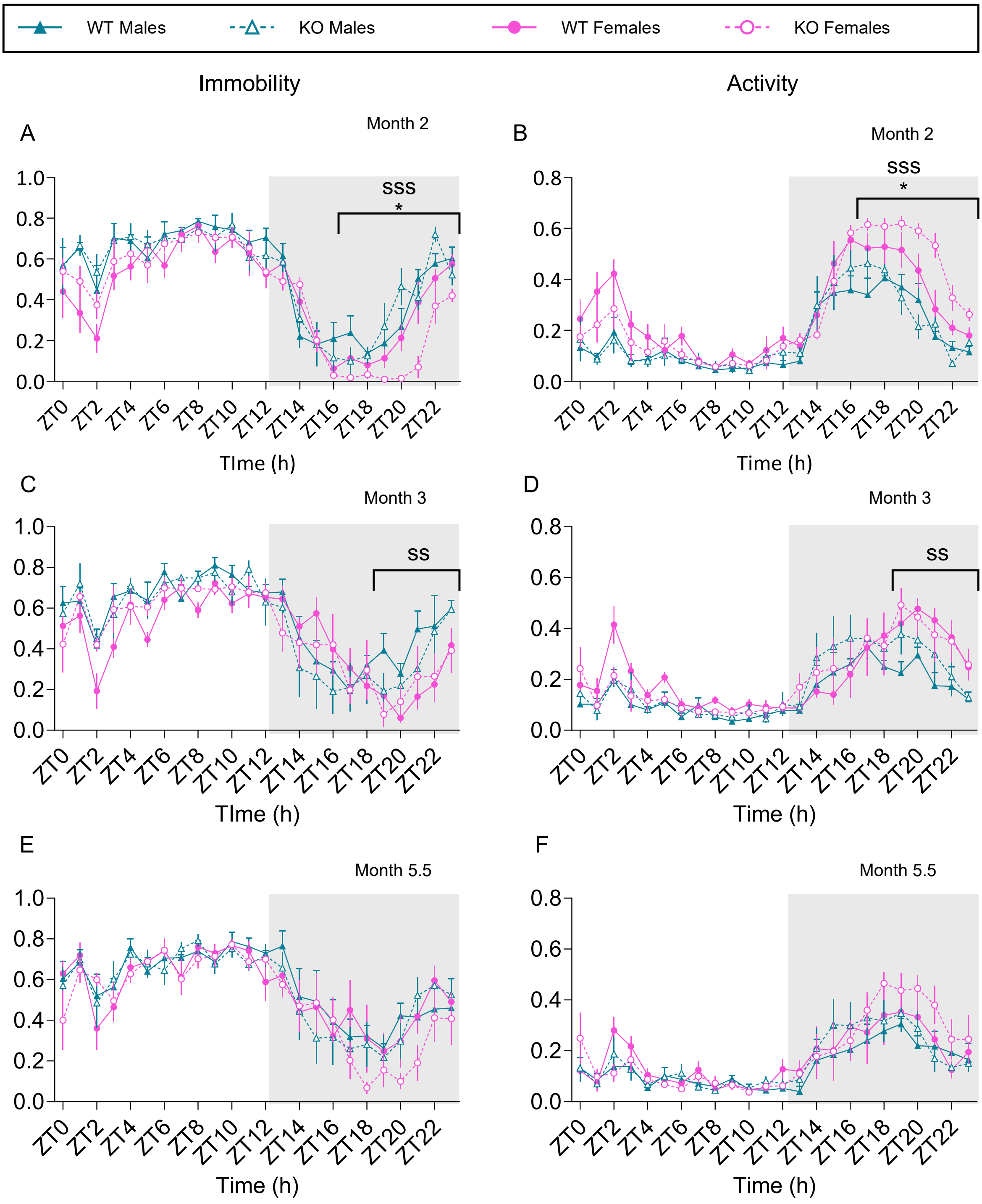

**Figure S8. There is an effect of genotype on daily activity rhythms between male and female *Fkbp5* KO and WT up to 2 months after MIA injection.** (**A-F**) 24h immobility/activity plots using 1h bins. (**A, B**) Average of 24h plots across 7 days, 2 months after MIA injection. N=4/4/4/4. (**C, D**) Average of 24h plots across 7 days, 3 months after MIA injection. N=4/4/4/4. (**E, F**) Average of 24h plots across 7 days, 5.5 months after MIA injection. N=4/4/4/4. ZT0=7am, ZT12=7pm. Data shows mean ± S.E.M. Genotype effect: *P<0.05; sex effect: $^{ss}$ P<0.01; $^{sss}$ P<0.01. See full statistical analysis in Supplementary Table S3.

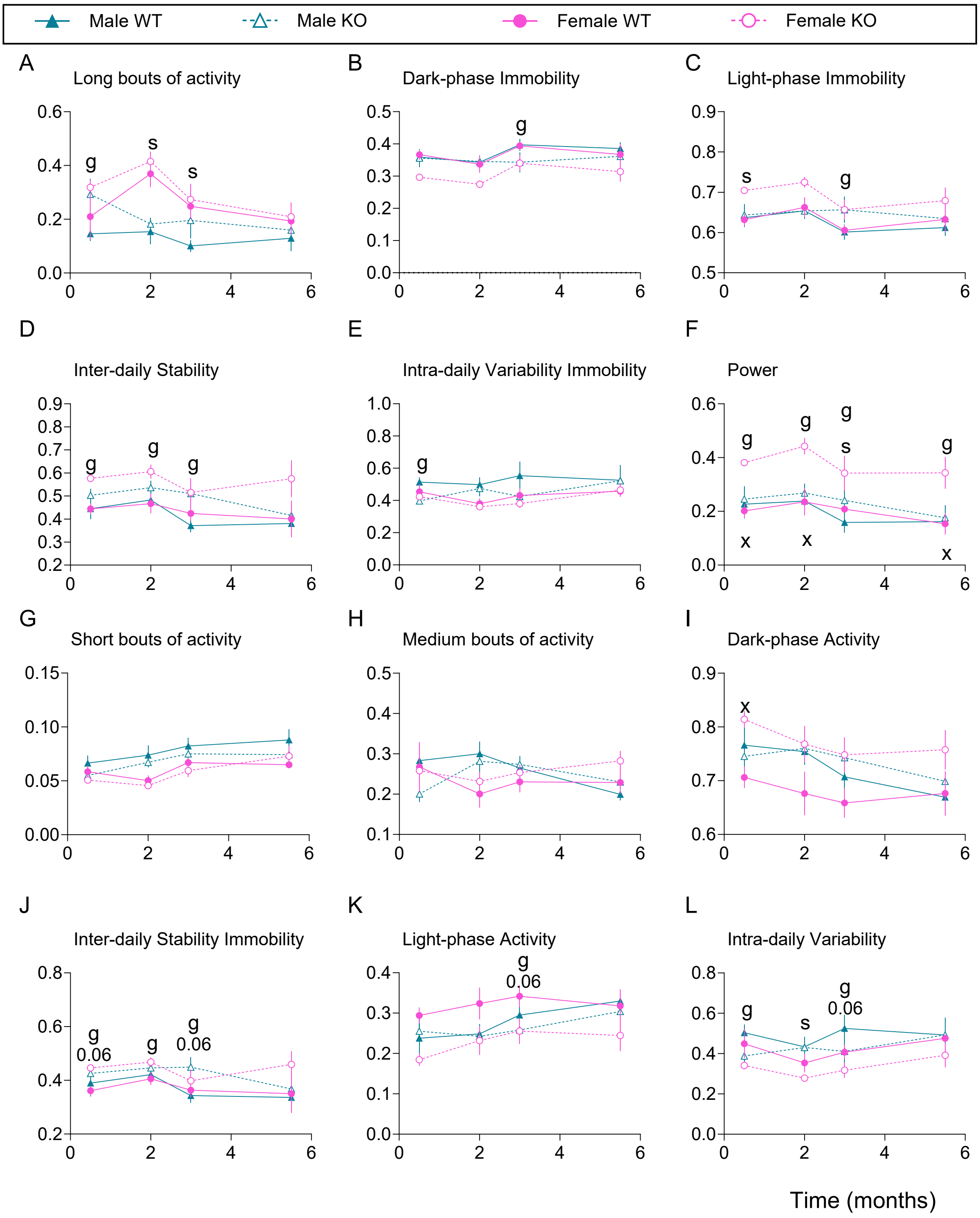

**Figure S9. The effect of *Fkbp5* global deletion on daily activity rhythms is not significant after 3 months.** Various summary statistics of circadian disruption calculated across a 6 day-period post-MIA injection. Data shows mean ± S.E.M. N=4/4/4/4. Genotype effect: g: $P<0.05$; sex effect: s: $P<0.05$; sex x genotype interactions: g x s. s: sex effect $P<0.05$; g: genotype effect $P<0.05$; x: sex x genotype $P<0.05$. See full statistical analysis in Supplementary Table S3.

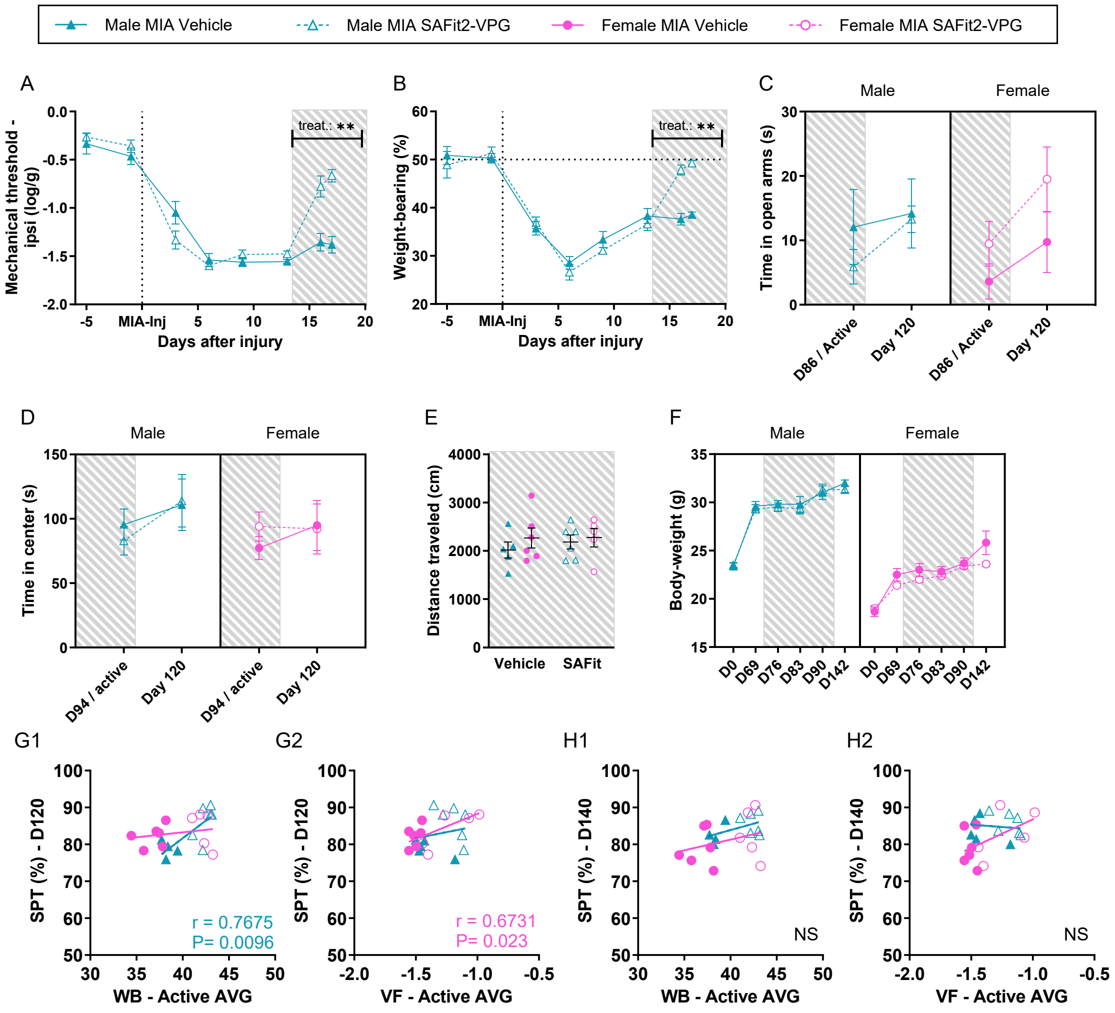

**Figure S10. Pharmacological inhibition of FKBP51 shortly after MIA induction improves sensory and functional deficits, but does not modify anxiety-like behavior, locomotor activity or body-weight.** All mice were subjected to MIA-injury and SAFit2-VPG was administered at different time points after injury, grey area suggests when compound was active. (**A**) SAFit2-VPG produced significant improvements on mechanical hypersensitivity when administered 2 weeks after injury in male mice. (N=7-8). (**B**) SAFit2-VPG produced significant improvements on weight bearing asymmetry when administered 2 weeks after injury in male mice. (N=7-8). (**C**) Anxiety-like behavior at 13 weeks in the Elevated Plus Maze was unaltered by SAFit2-VPG administration from 10-14 weeks after injury. (N=5-6). (**D**) Anxiety-like behavior at 14 weeks in the Open Field Test, was unaltered by administration of SAFit2-VPG from 10-14 weeks after injury (N=5-6). (**E**) Locomotor activity at 14 weeks, as suggested by the distance traveled in the Open Field Test, was unaltered by the administration of SAFit2-VPG at 10-14 weeks after injury (N=5-6). (**F**) Bodyweight was unaltered by treatment with SAFit2-VPG at 10-14 weeks after injury (N=5-6). (**G1,2**) Correlations between the weighted average weight bearing asymmetry (WB) or mechanical hypersensitivity (VF) from the active phase of administration (10-14weeks) and the level of depressive-like behavior 3 weeks after the active phase. (**H1,2**) Correlations between the weighted average weight bearing asymmetry (WB) or mechanical hypersensitivity (VF) from the active phase of administration (10-14weeks) and the level of depressive-like behavior 6 weeks after the active phase were not significant. Data shows mean ± S.E.M. *P<0.05, **P<0.01, ***P<0.001, as determined by the ANOVA analysis. (See full statistical analysis in Supplementary Table S3).

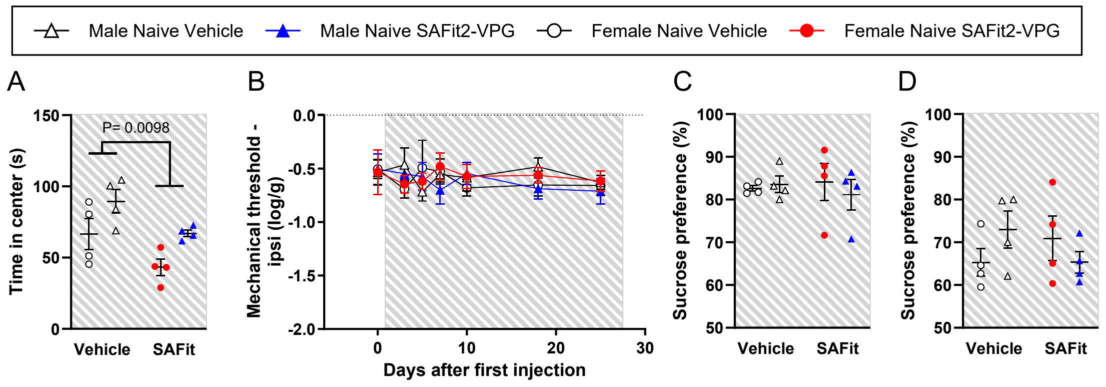

**Figure S11. Pharmacological inhibition of FKBP51 in naïve mice induces anxiety-like behaviour, but does not modify mechanical threshold or depressive-like behaviour.** All mice were naïve and SAFit2-VPG was administered for a total duration of 4 weeks, hatched grey area suggests when compound was active. (**A**) SAFit2-VPG produced a significant anxiety-like behavior across sex, as assessed in the Open Field Test at D23 after first injection (N=4). P-value suggested by 2way ANOVA result. (**B**) Mechanical sensitivity was unaltered by administration of SAFit2-VPG for 4 weeks (N=4). (**C**) Depressive-like behavior, as measured via the preference for a 1% sucrose solution, was unaltered by administration of SAFit2-VPG for 4 weeks (N=4). (**D**) Depressive-like behavior, as measured via the preference for a lower concentration (0.5%) of sucrose solution, was also unaltered by administration of SAFit2-VPG for 4 weeks (N=4). For all graphs; Data shows mean ± S.E.M. (See full statistical analysis in Supplementary Table S3).

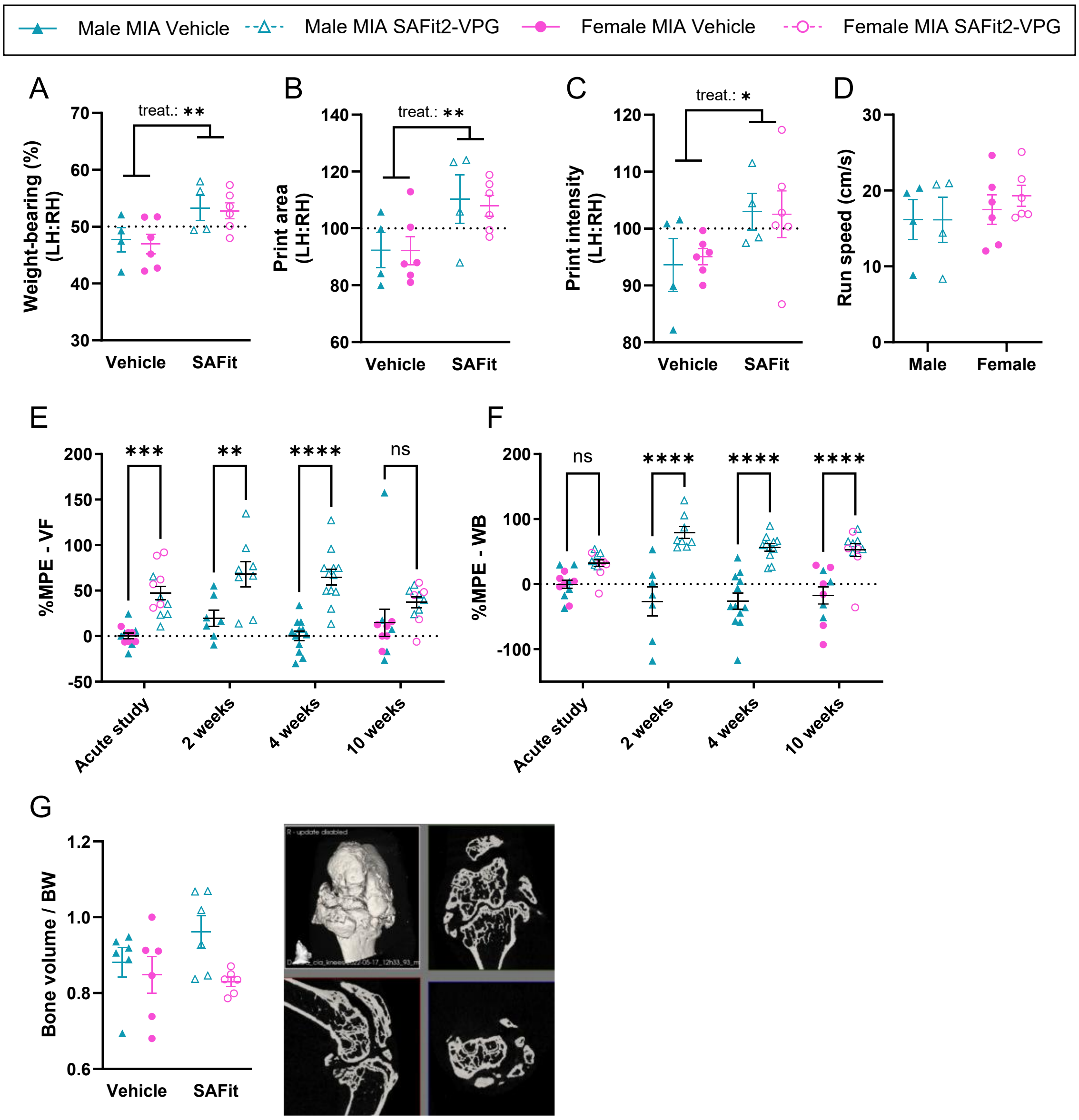

**Figure S12. Pharmacological inhibition of FKBP51 at any time during the pain state improves mechanical hypersensitivity and weight bearing. A-D, G** All mice were subjected to MIA-injury and SAFit2-VPG was administered from 3 days before and lasting for approximately 11 days after injury. Gait assessment on the catwalk was performed at 6 weeks after injury. Early treatment prevented the full development of gait deficits across sex (measured via 2-way ANOVA) on all of the following parameters: (**A**) Dynamic weight bearing as a measure of the amount of weight places on the left vs right hindleg while walking, (**B**) the size of the print-area of the ipsilateral (left) paw, compared with the contralateral (right), and the (**C**) print intensity of the paw-print from the ipsi- vs contra-lateral paw. (**D**) The run speed was unaltered by treatment, suggesting that any changes in gait, was not secondary effects of alteration in general speed of movement. (**E-F**) %MPE of FKBP51-inhibition was calculated for the first measure after the first injection of SAFit2-VPG in the different studies (D3 for post-MIA treatment studies, and day 2 for pre-MIA/acute treatment), for comparison across studies (N=7-12), for the mechanical sensitivity (E) and weight bearing asymmetry (F). (**G**) Joint deformation at 6 months after injury was unaltered by early treatment with SAFit2-VPG. For all graphs: N=4-6. Data shows mean ± S.E.M. *P<0.05, **P<0.01, ***P<0.001, ****P<0.0001, as determined using ANOVA for A-D, and appropriate post-tests for E-F. See full statistical analysis in Supplementary Table S3.

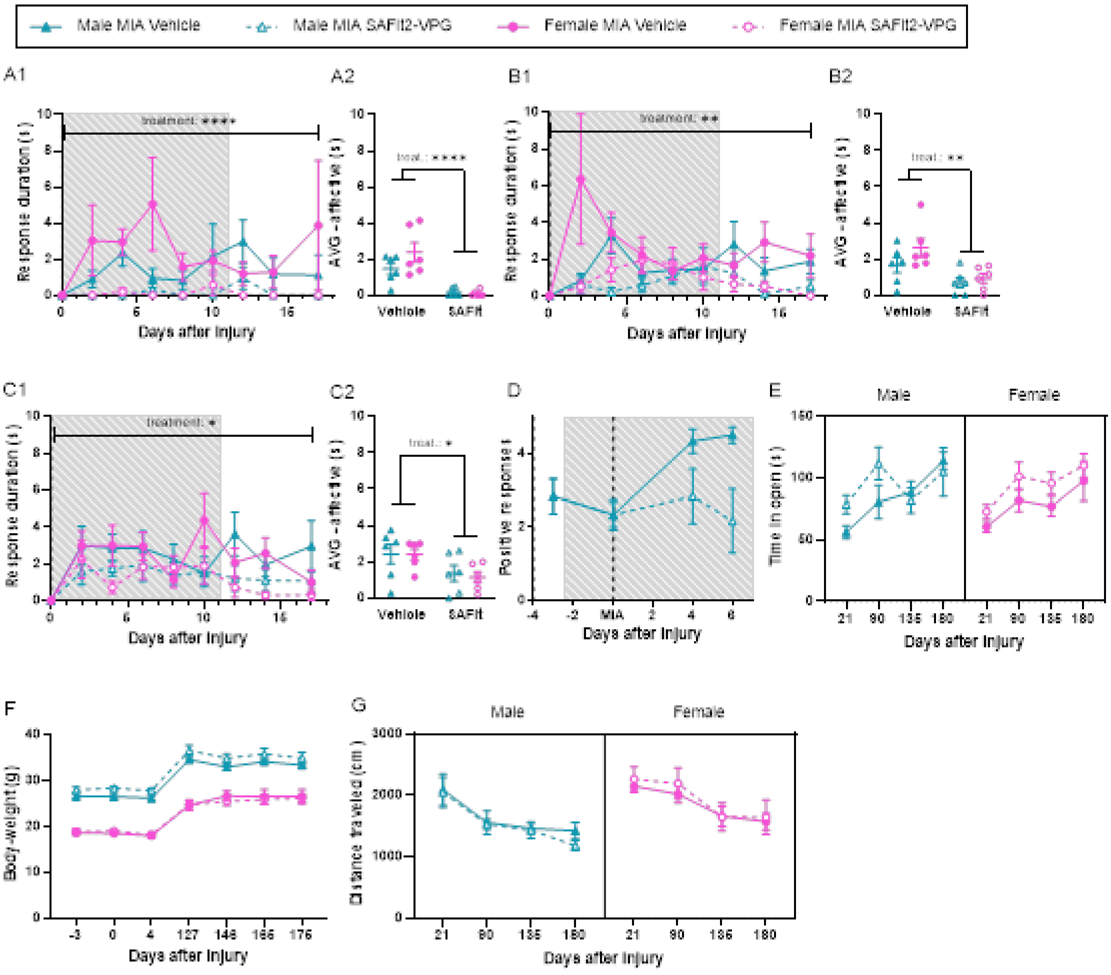

**Figure S13. Acute pharmacological inhibition of FKBP51 during disease onset affects affective behaviours.** All mice were subjected to MIA-injury and SAFit2-VPG was administered from 3 days before and lasting for approximately 11 days after injury, hatched grey area suggests when the compound was active. (**A1,2**) Early treatment with SAFit2-VPG produced a significant lower level of affective responding to mechanical stimulation with a low intensity filament (0.04g) following MIA injury (N=6). (**B1,2**) Early treatment with SAFit2-VPG produced a significant lower level of affective responding to mechanical stimulation with a medium intensity filament (0.16g) following MIA injury (N=6). (**C1,2**) Early treatment with SAFit2-VPG produced a significant lower level of affective responding to mechanical stimulation with a high intensity filament (1.0g) following MIA injury (N=6). (**D**) Early treatment with SAFit2-VPG significantly decreased the level of brush allodynia after injury (N=6). (**E**) Time course development of anxiety-like behavior in the Open Field Test, when treatment was SAFit2-VPG was administered around the onset of injury. (**F**) Body weight development was unaltered by early SAFit2-VPG treatment (N=6) (**G**) Distance traveled (cm) in the open field was unaltered by early treatment with SAFit2-VPG (N=6). For Fig A2, B2, C2: Weighted average (AVG) was calculated for the full study period. Data shows mean ± S.E.M. *P<0.05, **P<0.01, ***P<0.001, as determined via ANOVA analysis. See full statistical analysis in Supplementary Table S2.

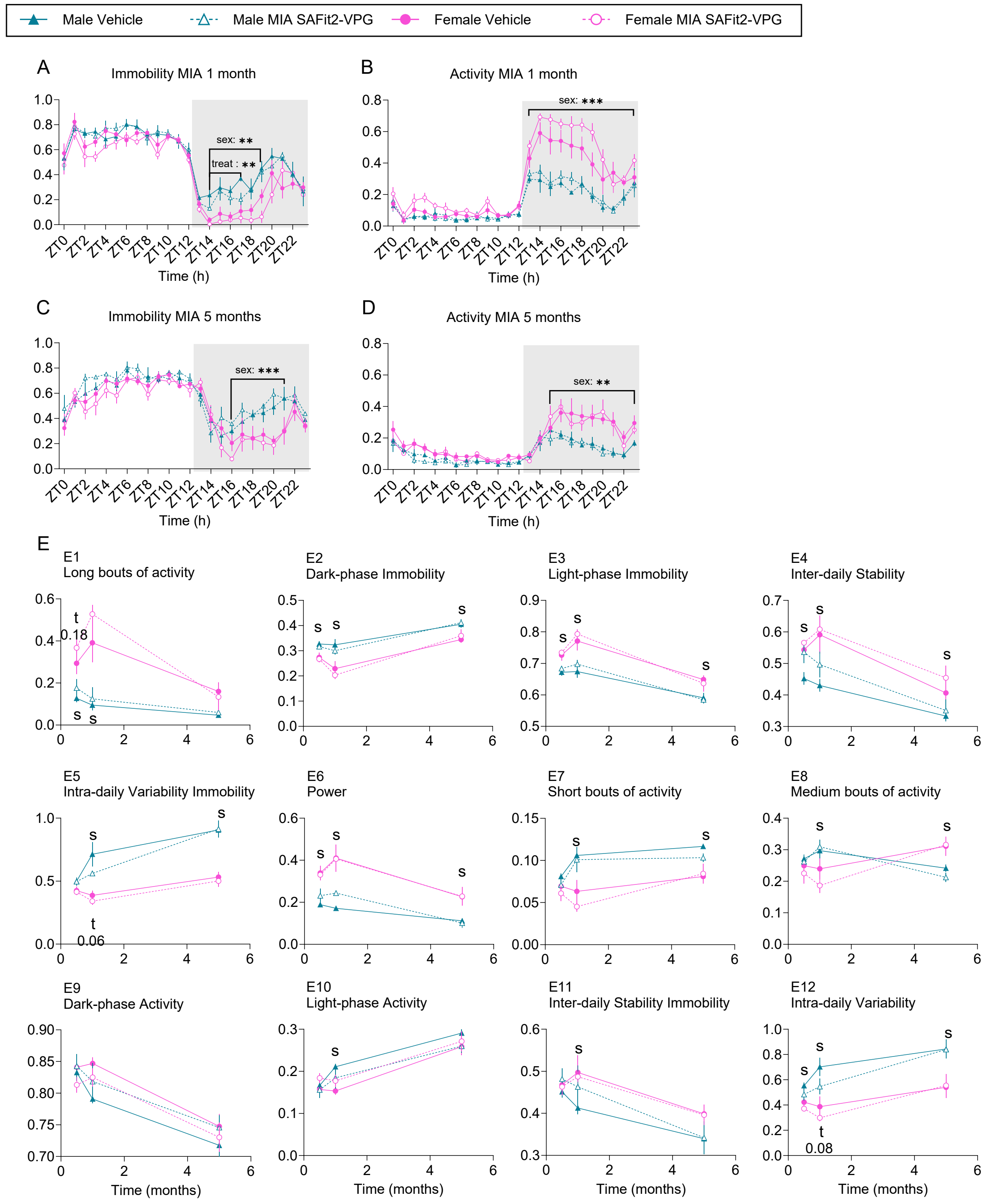

**Figure S14: There are no obvious effects of acute SAFit2-VPG treatment on daily activity rhythms between male and female WT from one month after MIA injection.** (**A-D**) 24h immobility/activity plots using 1h bins. (**A, B**) Average of 24h plots across 7 days, 1 month after MIA injection. N=4/4/4/4. (**C, D**) Average of 24h plots across 7 days, 5 months after MIA injection. N=4/4/4/4. (**E**) Various summary statistics of circadian disruption calculated across a 6 day-period post-MIA injection. ZT0=7am, ZT12=7pm. Data shows mean ± S.E.M. N=4/4/4/4. Treatment effect: t; sex effect: s: P<0.05. See full statistical analysis in Supplementary Table S3.

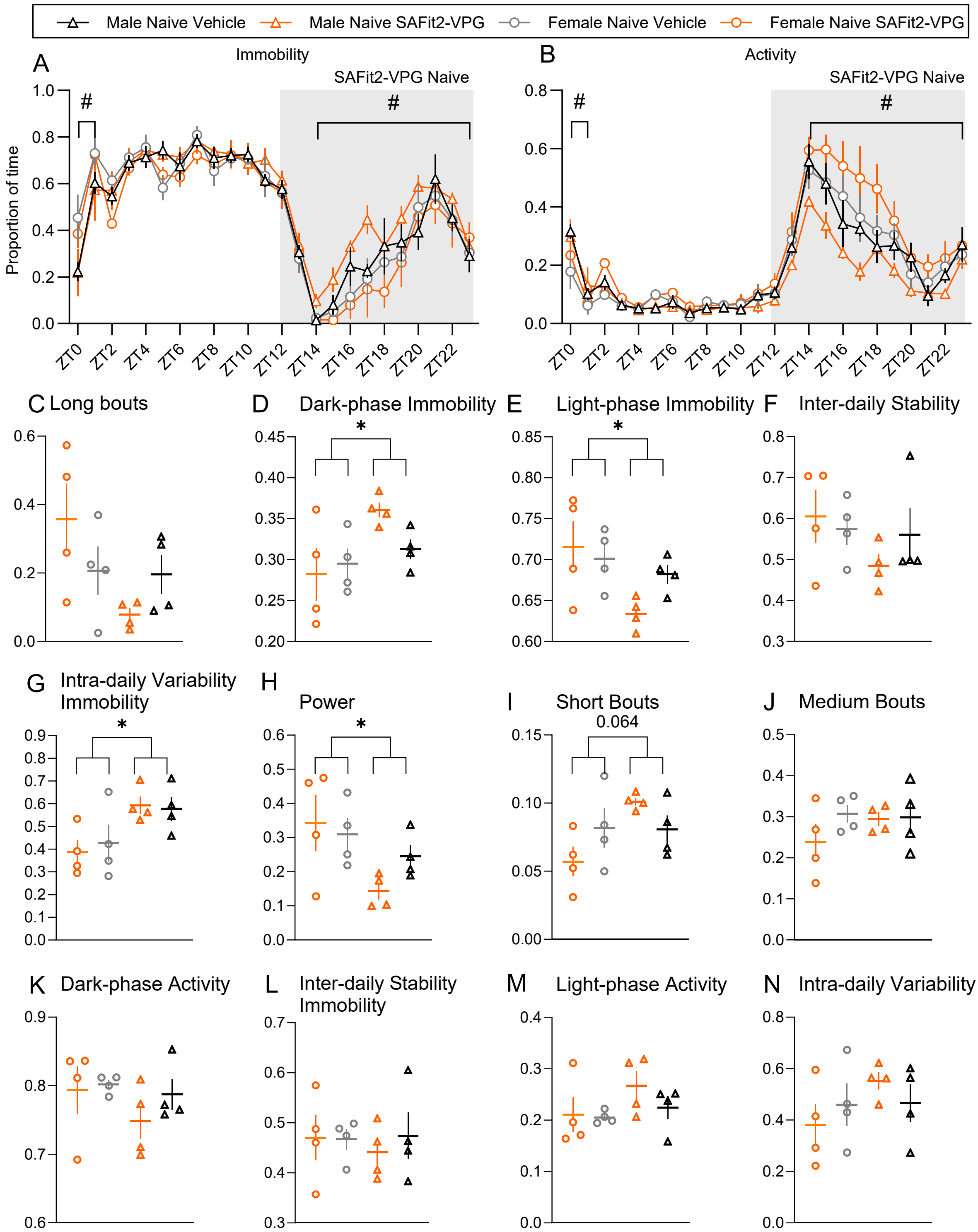

**Figure S15: SAFit2 alone promotes sex differences in daily activity rhythms in naïve 8-week-old mice. (A, B)** 24h immobility/activity plots using 1h bins. Average of 24h plots across 6 days of SAFit2-VPG exposure. N=8. SAFit2-VPG alone induces changes to immobility (A) and activity (B) patterns in naive, 8-week-old male and female mice. (**C-N**) Various summary statistics of circadian disruption calculated across a 6 day-period post-MIA injection. ZT0=7am, ZT12=7pm. Data shows mean ± S.E.M. and single data points. N=4/4/4/4. *P<0.05. See full statistical analysis in Table S3.

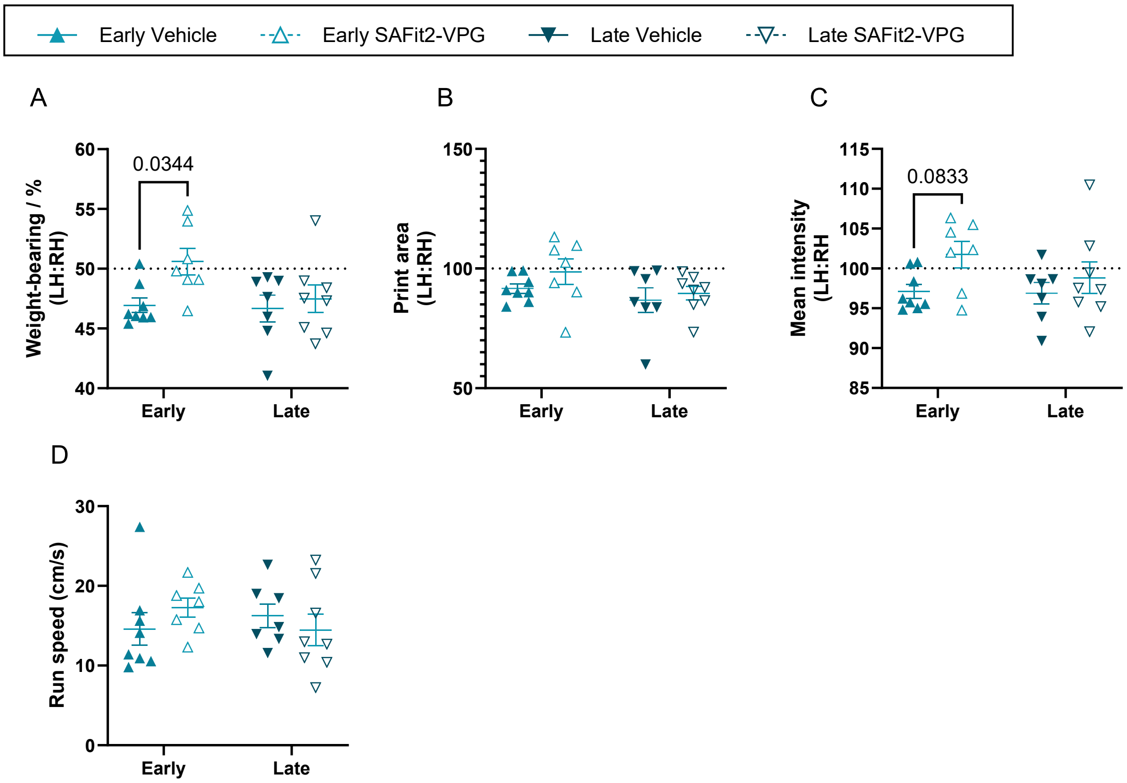

**Figure S16. Acute pharmacological inhibition of FKBP51 initiated before or after MIA induction prevents the full development of gait deficits.** All male mice were subjected to MIA-injury, and SAFit2-VPG was administered either “Early” from 3 days before and lasting for approximately 11 days after injury, or “Late” from D20-30. (**A-D**) Gait assessment on the catwalk was performed at D23 after injury, where both treatment-types showed beneficial effects on mechanical sensitivity, but only "late"-treatment was in active treatment. SAFit2-VPG treatment induced improvement across treatment timing (measured via 2-way ANOVA) on Dynamic weight bearing (A), measured as the amount of weight places on the left vs right hindleg while walking, and the print intensity (C), but not the size of the print-area (B) of the ipsilateral (left) paw, compared with the contralateral (right). (**D**) The run speed was unaltered by treatment, suggesting that any changes in gait was not secondary effects of alteration in general speed of movement. For all graphs: N=7-8. Data shows mean ± S.E.M. and single data points. See full statistical analysis in Supplementary Table S3.

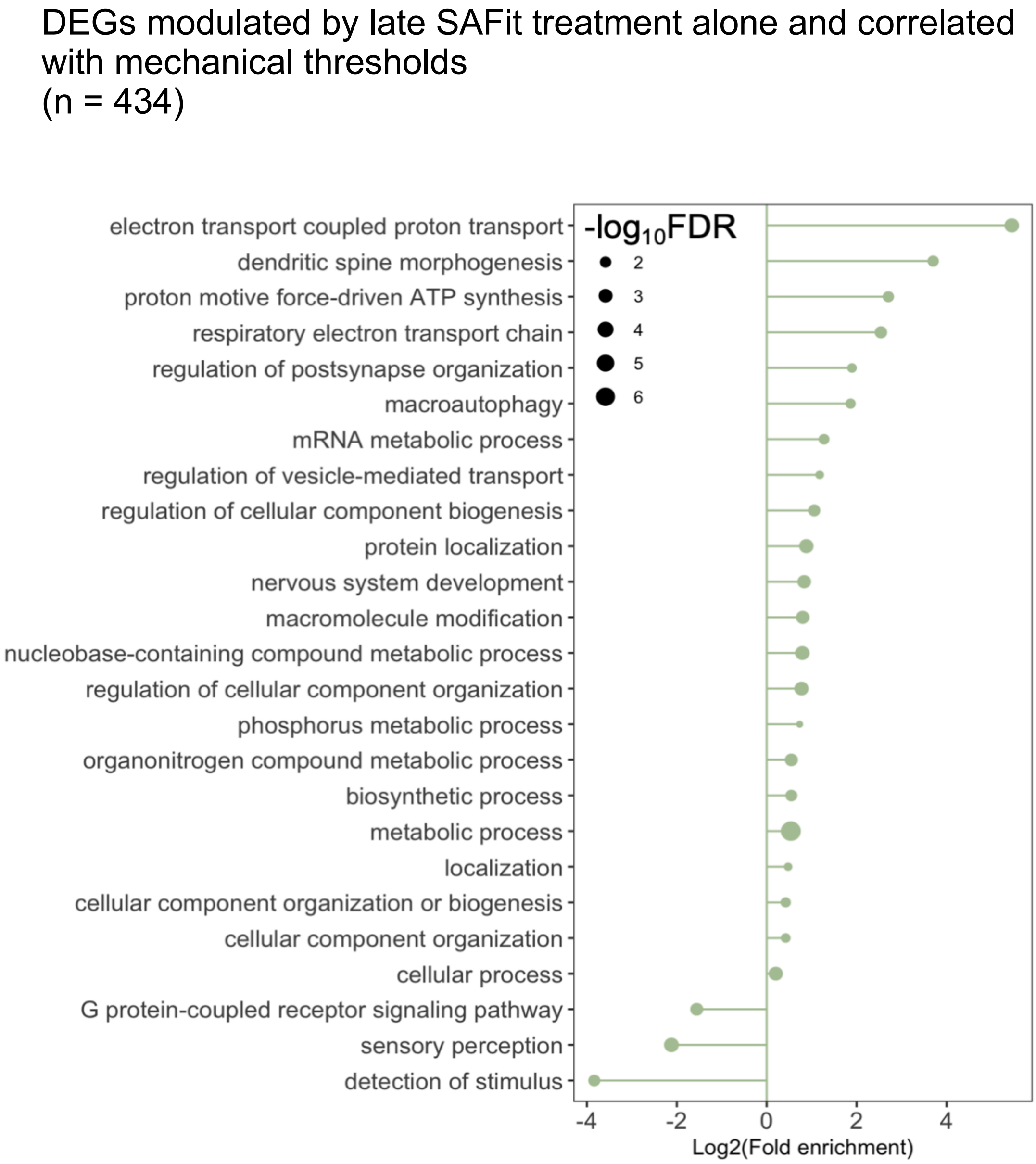

**Figure S17: Lollipop plot of enriched GO terms after gene expression analysis.** This figure displays the enriched Gene Ontology (GO) terms identified from the differentially expressed genes (DEGs) between $Late_{SAFit2}$ and $Late_{Vehicle}$ treated mice. The X-axis represents GO terms associated with biological processes while the Y-axis shows the fold change in enrichment. Significance (-log10 adjusted P-value) of each term is denoted by size.

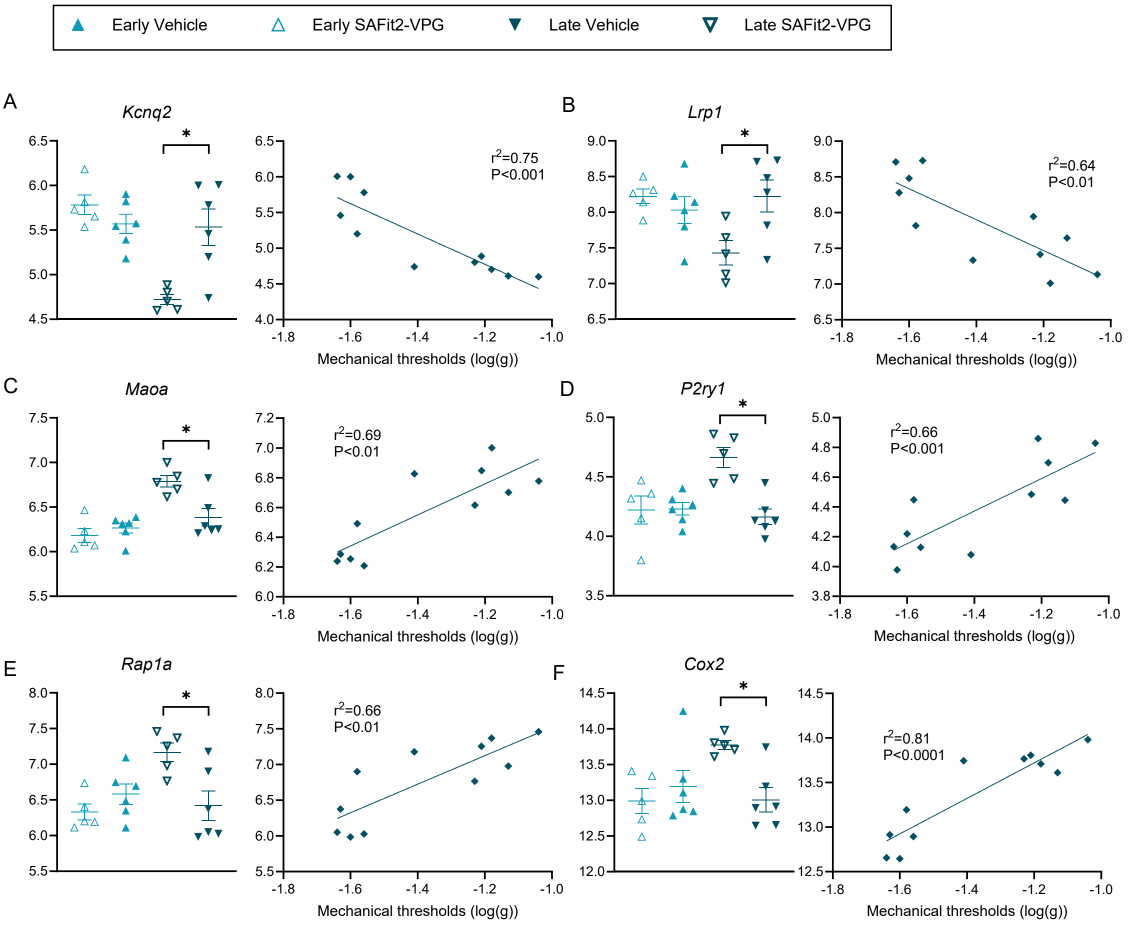

**Figure S18: Selected DEGs are regulated by late SAFit2 treatment in a direction that may promote hypersensitivity.** *Kcnq2* encode the protein Kv7.2; *Lrp1* encode the Low-Density Lipoprotein Receptor-Related Protein 1; *Maoa* encode the Mono oxydase A protein that breakdown monoamines; *P2ry1* encode the purinergic receptor P2Y1; *Rap1a1* encode the protein Rap1a, a small GTPase from the Ras family; *Cox2* encode the cyclooxygenase-2 enzyme, involved in the synthesis of prostaglandins that mediate inflammation and pain. Data shows mean ± S.E.M. and single data points. Y-axis: $log_2$CPM.

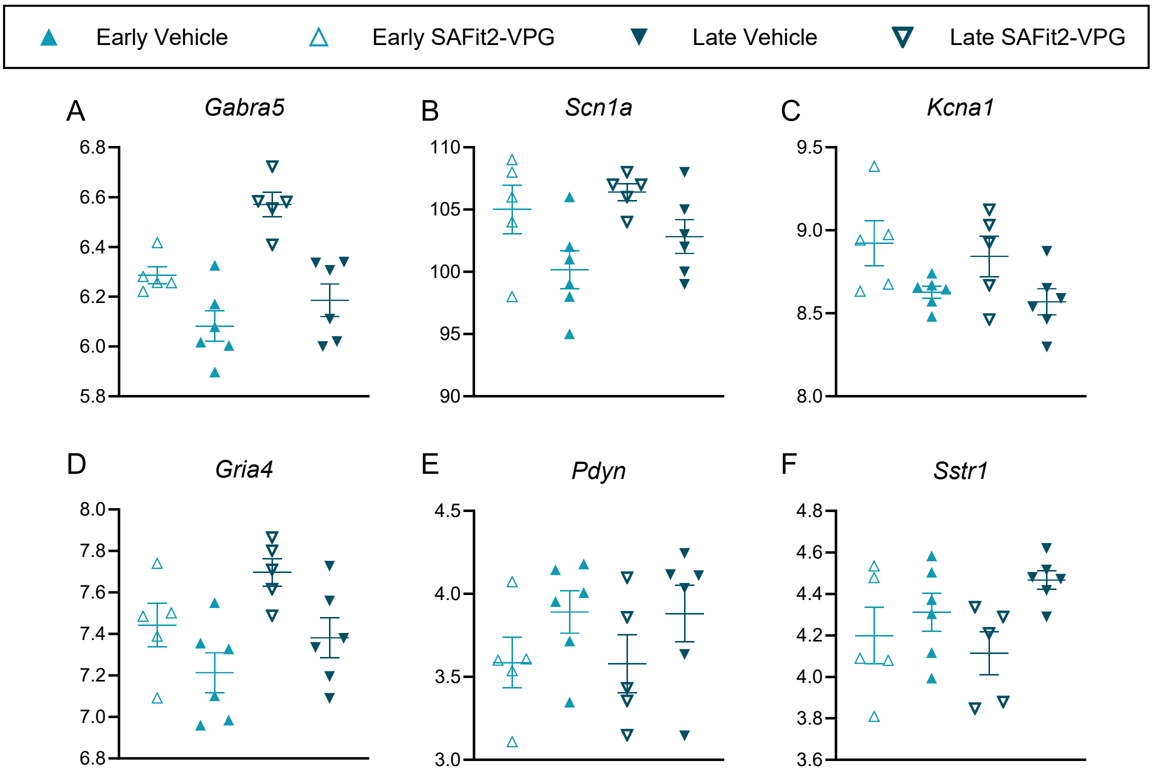

**Figure S19: A number of DEGs similarly regulated by early and late SAFit2 treatment are involved with nociceptive signalling.** *Gabra5* encodes the a5 subunit of the GABA A receptor; *Scna1a* encodes the sodium channel Nav1.1; *Kcna1* encodes the potassium channel Kv1.1; *Gria4* encodes the glutamate ionotropic receptor AMPA type subunit 4; *Pdyn* encodes prodynorphin, a pro-nociceptive peptide, and *Sstr1* encodes the somatostatin receptor 4. Data shows mean ± S.E.M. and single data points. Y-axis: log2CPM. For all target, SAFit2-VPG *vs* Vehicle, $P<0.05$.

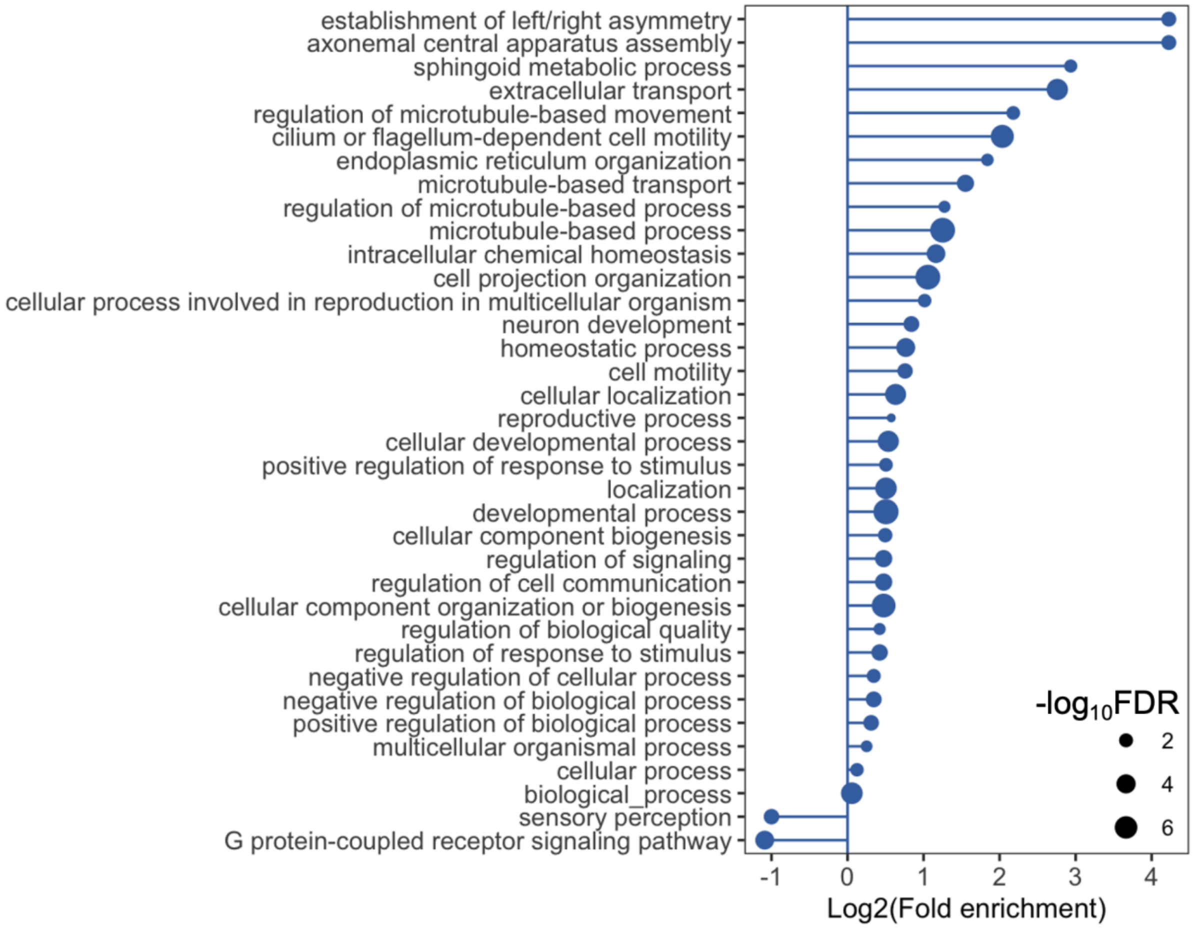

**Figure S20: Lollipop plot of enriched GO terms after differential gene expression analysis.** This figure displays the enriched Gene Ontology (GO) terms identified by PANTHER using differentially expressed genes derived from the main treatment effect of SAFit2-VPG *vs* vehicle (Figure 8 in the main text). The X-axis represents GO terms associated with biological processes while the Y-axis shows the fold change in enrichment. The size of the lollipop represents the significance (-log10 adjusted P-value) of each term.

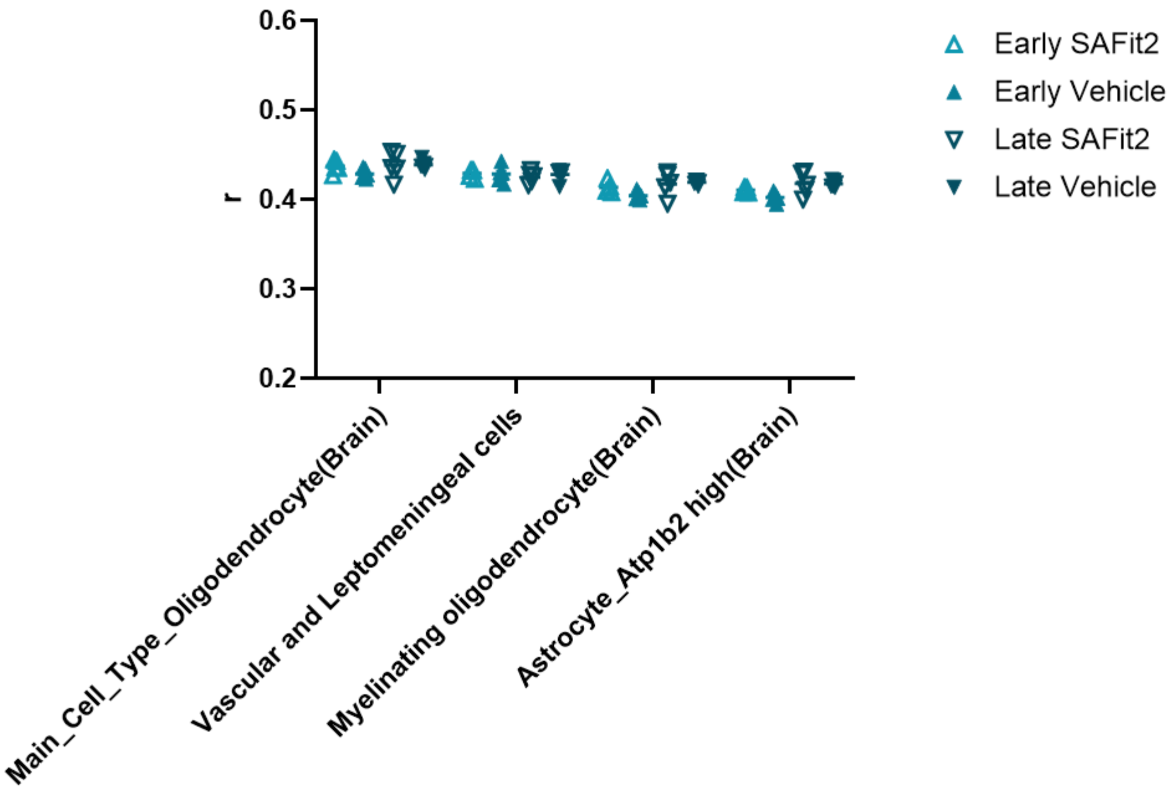

**Figure S21: Correlation analysis between the major cell types in spinal cord samples showed no difference between SAFit2 and Vehicle treated mice.** Correlation coefficients between spinal cord samples and the predominant cell types derived from the mouse cell atlas were evaluated using RNA sequencing data. The degree of correlation between each cell type identified from bulk RNA sequencing data was no different between SAFit2 and vehicle treated mice spinal cord samples. These results suggested that cell type proportions had not changed after SAFit2. N=5-6/group.

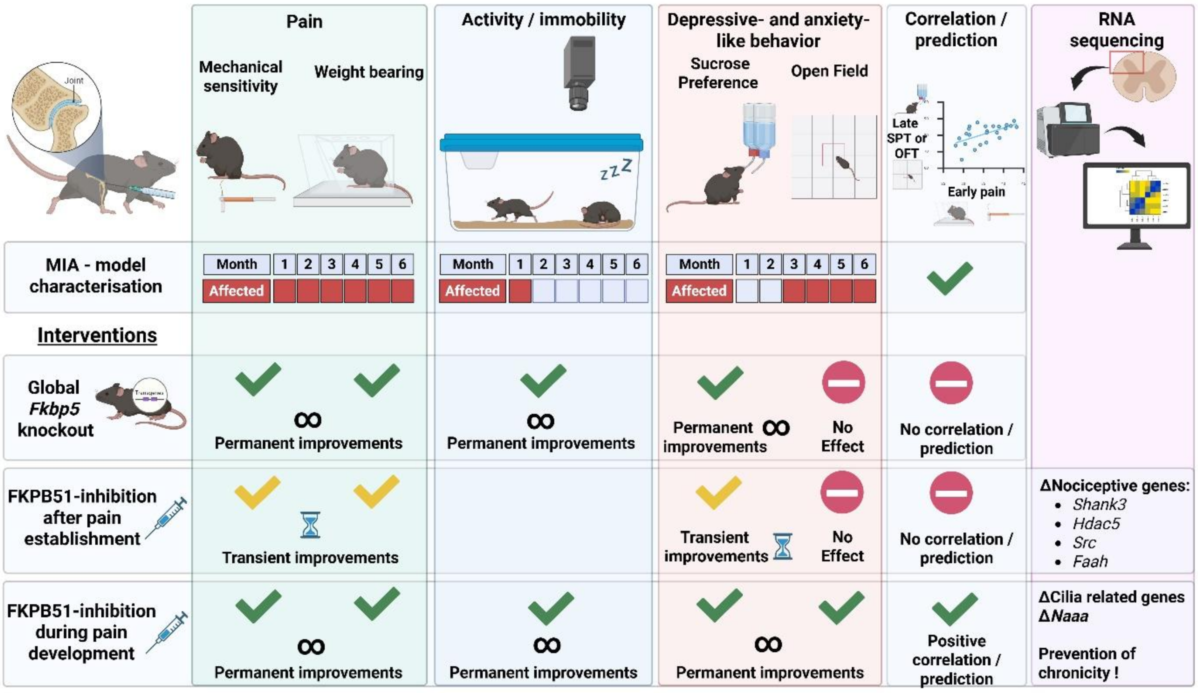

**Figure S22: Summary of findings**. The MIA model induces a long-lasting pain state with: (1) pain-like behaviour evident from disease onset for up to 6 months, (2) no obvious changes to immobility and activitypatterns beyond the first month and (3) emotional comorbidities apparent from 3 to 6 months. Crucially, early pain symptoms can predict long-term sensory and emotional outcomes. Global knock down of *Fbpb5* improves pain-, activity- and depressive-related symptoms, but there is no correlation between pain- and emotional outcomes. Pharmacological inhibition of the stress regulator FKBP51 after the full establishment of the pain state only provides transient pain and emotional relief that is accompanied by transient changes in expression of nociceptive genes at spinal cord level. In contrast, FKBP51 inhibition at disease and pain onset permanently improves the sensory and emotional symptoms and persistently downregulates genes related to pain chronicity.

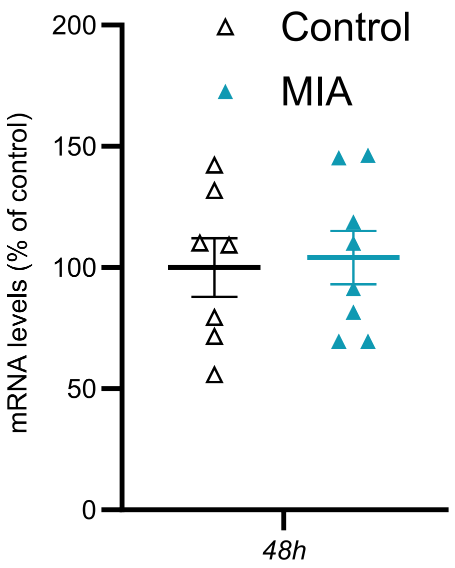

**Figure S23: There is no changes in *Fkbp5* mRNA in L4, L5 DGRs 48h after injection of MIA into the knee joint.** Data shows single data point and mean +/- SEM.

**Table S2. Complete information for statistical analysis of figures from the main manuscript.**

| Fig | Analysis | F-values | N pr group |
|---|---|---|---|
| **Fig 1. MIA characterization in male and female mice** | | | |
| Fig 1.A1. Mechanical allodynia (VF) | 3 way RM ANOVA, injury*sex*time | $F_{injury}$ (1,19)= 372.2, P<0.0001<br>$F_{time}$ (9,171)= 15.78, P<0.0001<br>$F_{time*injury}$ (9,171)= 15.81, P<0.0001 | 5-7 |
| Fig 1.A2. VF - weighted AVG. | 2 way ANOVA, injury*sex | $F_{injury}$ (1,19)= 444.4, P<0.0001<br>$F_{sex}$ (1,19)= 3.894, P=0.0632 (NS)<br>$F_{sex*injury,}$ P=0.1207 (NS) | 5-7 |
| Fig 1.B1. Functional deficit (WB) | 3 way RM ANOVA, injury*sex*time | $F_{injury}$ (1,18)= 517.4, P<0.0001<br>$F_{sex}$ (1,18)= 6.530, P=0.0199<br>$F_{time}$ (15,240)= 41.8, P<0.0001<br>$F_{time*injury}$ (15,240)= 42.52, P<0.0001<br>$F_{sex*injury}$ (1,18)= 13.19, P=0.0019<br>$F_{sex*injury*time}$ (15,270)= 1.815, P=0.0326. | 4-6 |
| Fig 1.B2. WB - weighted AVG | 2 way ANOVA, injury*sex | $F_{injury}$ (1,18)= 470.2, P<0.0001<br>$F_{sex}$ (1,18)= 8.210, P=0.0103<br>$F_{sex*injury}$ (1,18)= 14.77, P=0.0012 | 4-6 |
| Fig 1.C1. Joint deformation | 2 way ANOVA, injury*sex | $F_{injury}$ (1,17)= 101.4, P<0.0001<br>$F_{sex}$ (1,17)= 21.01, P<0.0001<br>$F_{sex*injury}$ = NS | 4-11 |
| Fig 1.D. SPT, 1-6M | Mixed-effects model$^{\#}$, injury*sex*time | $F_{injury}$ (1,41)= 11.81, P=0.0013<br>$F_{time*injury}$ (5,119)= 6.086, P<0.0001 | 4-12 |
| Fig 1.D2. SPT, 3-6M AVG | 2 way ANOVA, injury*sex | $F_{injury}$ (1,41)= 11.87, P=0.0013 | 11-12 |
| Fig 1.E1. OFT - 2M | 2 way ANOVA, injury*sex | NS | 4-6 |
| Fig 1.E2. OFT - 3M | 2 way ANOVA, injury*sex | $F_{injury}$ (1,18)= 6.462, P=0.0204 | 4-6 |
| Fig 1.E3. OFT - 6M | 2 way ANOVA, injury*sex | $F_{injury}$ (1,19)= 3.768, P=0.0673 (NS) | 5-7 |
| Fig 1F1. Correlation matrix, 3M | Pearson's correlation analysis | WB/VF: r = 0.951, P<0.0001.<br>WB/OFT: r = 0.510, P=0.015.<br>WB/SPT: r = 0.337, P=0.125 (NS).<br>VF/OFT: r = 0.513, P=0.015.<br>VF/SPT: r = 0.320, P=0.041.<br>OFT/SPT: r = 0.240, P=0.28. (NS) | 22, across groups |
| Fig 1F2, correlation matrix, 6M. | | VF/OFT: r = 0.414, P=0.050.<br>VF/SPT: r = 0.561, P=0.005.<br>OFT/SPT: r = - 0.066, P=0.77. (NS) | 23, across groups |
| Fig 1.G1. VF 2W AVG vs 6M VF | Simple linear regression, and Pearson's correlation analysis | r = 0.8613, F(1,21)=60.34, P<0.0001 | |
| Fig 1.G2. VF-2W AVG vs 6M SPT | Simple linear regression, and Pearson's correlation analysis | r = 0.5935, F(1,21)=11.42, P=0.0028 | |
| **Fig 2. Immobility / activity** | | | |
| Fig 2A | 2-way RM ANOVA | NS | 8 |
| Fig 2B | 2-way RM ANOVA | NS | 8 |

| | | | |
|---|---|---|---|
| Fig 2C | 3-way RM ANOVA injury*sex*time | All time course:<br>Injury effect: F(1,12)= 15.5, P=0.002;<br>Time x sex interactions: F(23, 276)= 2.4, P=0.035; no sex and no sex x injury significance.<br>Time x injury x sex: F(23, 276)= 2.4, P=0.035<br><br>ZT14 to ZT1:<br>Injury: F(1,12)=13.7; P=0.003<br>Time x Sex interactions: F(11, 132)=3.3, P=0.013. no sex and no sex x injury significance.<br>Time x injury x sex: F(11, 132)= 2.9, P=0.024 | 4 |
| Fig 2D | 3-way RM ANOVA injury*sex*time | All time course:<br>Injury effect: F(1,12)=4.4, P=0.058; Sex NS; Sex x Treat: NS.<br><br>ZT14 to ZT1: Injury effect: F(1,12)=4.9, P=0.046; no sex, no sex x treat significance; Time x injury x sex: F(11, 132)= 2.8, P=0.034. | 4 |
| Fig 2E | 2-way ANOVA | No effect of sex or sex x injury interactions; Injury effect: E1, F(1,12)=5.02, P-=0.045; E2 F(1,12)=9.9, P=0.008; E3: F(1,12)=8.6, P=0.012. | 4 |
| **Fig 3. Effects of genetic knockout of FKBP51 on MIA-induced behavior.** | | | |
| Fig 3.A1. KO/WT - VF | 3 way RM ANOVA, genotype*sex*time | $F_{genotype}$ (1,18)= 82.47, P<0.0001<br>$F_{time}$ (16,288)= 67.20, P<0.0001<br>$F_{time*genotype}$(16,288)= 4.896, P<0.0001 | 5-6 |
| Fig 3.A2, KO/WT - VF AVG | 2 way ANOVA, genotype*sex | $F_{genotype}$ (1,18)= 91.55, P<0.0001 | 5-6 |
| Fig 3.B1. KO/WT - WB | 3 way RM ANOVA, genotype*sex*time | $F_{genotype}$ (1,18)= 194.2, P<0.0001<br>$F_{sex}$ (1,18)= 20.92, P=0.0002<br>$F_{time}$ (17,306)= 106.7, P<0.0001<br>$F_{time*sex}$ (17,308)= 1.853, P=0.0217<br>$F_{time*genotype}$(17,306)= 11.33, P<0.0001<br>$F_{sex*genotype}$= P=0.54 (NS) | 5-6 |
| Fig 3.B2, KO/WT / WB-AVG | 2 way ANOVA, genotype*sex | $F_{genotype}$ (1,18)= 224.3, P<0.0001<br>$F_{sex}$ (1,18)= 14.74, P=0.0012<br>$F_{sex*genotype}$ = P=0.2 (NS) | 5-6 |
| Fig 3.C. Bone volume (corrected for BW) | 2 way ANOVA, genotype*sex | $F_{genotype,}$ P=0.59 (NS)<br>$F_{sex}$ (1,16)= 6.668, P=0.0200<br>$F_{sex*genotype}$ = P=0.38 (NS) | 4-6 |
| Fig 3.D. SPT | Mixed-effects model#, injury*sex*time | $F_{genotype}$ (1,34)= 8.256, P=0.0057<br>$F_{time}$ (3,70)= 3.532, P=0.0191 | 5-9 |
| Fig 3.E. OFT | 3 way RM ANOVA, genotype*sex*time | $F_{genotype}$, P=0.72 (NS)<br>$F_{time}$ (4,72)= 6.246, P=0.0002 | 5-6 |
| Fig 3.F1. correlation matrix, 3M | Pearson's correlation analysis | WB/VF: r = 0.616, P=0.002.<br>WB/OFT: r = 0.024, P=0.92 (NS).<br>WB/SPT: r = 0.279, P=0.21 (NS).<br>VF/OFT: r = 0.139, P=0.54. (NS)<br>VF/SPT: r = 0.230, P=0.30. (NS)<br>OFT/SPT: r = 0.319, P=0.126. (NS) | 22, Across groups |
| Fig 3.F2. correlation matrix, 6M | | WB/VF: r = 0.761, P<0.0001.<br>WB/OFT: r = - 0.011, P=0.96 (NS).<br>WB/SPT: r = 0.239, P=0.31 (NS).<br>VF/OFT: r = - 0.009, P=0.97 (NS).<br>VF/SPT: r = 0.281, P=0.23 (NS). | |

| | | | |
|---|---|---|---|
| | | OFT/SPT: r = 0.357, P=0.057. (NS) | |
| Fig 3.G1. VF 2W AVG vs 6M VF | Simple linear regression, and Pearson's correlation analysis | r = 0.7906, F(1,18)=30.01, P<0.0001 | |
| Fig 3.G2. VF-2W AVG vs 6M SPT | Simple linear regression, and Pearson's correlation analysis | r = 0.3412, F(1,19)=2.503, P=0.13 (NS) | |
| Fig 3.G3. VF 2W AVG vs 6M OFT | Simple linear regression, and Pearson's correlation analysis | r = 0.2011, F(1,18)=0.7587, P=0.39 (NS) | |
| **Fig 4. Treatment post injury** | | | |
| Fig 4.A1. Mechanical allodynia (VF) 4week study. | 2 way RM ANOVA, time*treatment | $F_{treatment}$ (1,22)= 49.52, P<0.0001.<br>$F_{time}$(19,418)= 79.44, P<0.0001<br>$F_{treatment*time}$(19,428)= 9.796, P<0.0001 | 12 |
| Fig 4.A2. Weighted Average (VF), Day 30-59 | Unpaired t-test | T=6.928, df=22. P<0.0001 | 12 |
| Fig 4.B1. Weight bearing (WB) 4week study | 2 way RM ANOVA, time*treatment | $F_{treatment}$(1,21)= 27.13, P<0.0001<br>$F_{time}$(16,336)= 64.38, P<0.0001<br>$F_{treatment*time}$(16,336)= 5.212, P<0.0001 | 11[$]-12 |
| Fig 4.B2. Weighted Average (WB), Day 30-59 | Unpaired t-test | T=8.097, df=21. P<0.0001 | 11[$]-12 |
| Fig 4.C1. Mechanical allodynia (VF) 2-3month | 3 way RM ANOVA, time*treatment*sex | $F_{treatment}$ (1,18)= 12.68, P=0.0022<br>$F_{time}$(19,342)= 72.63, P<0.0001<br>$F_{time*treatment}$(19,342)= 4.084, P<0.0001<br>$F_{sex,}$ P=0.1219 (NS)<br>$F_{sex*treatment,}$ P=NS | 5-6 |
| Fig 4.C2. Weighted Average (VF), Day 69-105 | 2 way ANOVA, treatment*sex | $F_{treatment}$(1,18)= 19.95, P=0.0003<br>$F_{sex}$ P= NS<br>$F_{sex*treatment,}$ P=NS | 5-6 |
| Fig 4.D1. Weight bearing (WB) 2-3month | 3 way RM ANOVA, time*treatment*sex | $F_{treatment}$(1,17)= 40.62, P<0.0001<br>$F_{time}$(20,340)= 40.26, P<0.0001<br>$F_{sex}$(1,17)= 4.304, P=0.0535 (NS)<br>$F_{time*sex}$(20,340)= 1.808, P=0.0188<br>$F_{time*treatment}$(20,340)= 6.037, P<0.0001 | 4[$]-6 |
| Fig 4.D2. Weighted Average (WB), Day 69-105 | 2 way ANOVA, treatment*sex | $F_{treatment}$(1,17)= 109.2, P<0.0001<br>$F_{sex}$(1,17)= 4.104, P=0.0588 (NS)<br>$F_{treatment*sex,}$ P=NS | 4[$]-6 |
| Fig 4.E1. Anhedonia (SPT) 2-3month | 3 way RM ANOVA, time*treatment*sex | $F_{treatment}$(1,18)= 6.255, P=0.0223<br>$F_{time}$(2,36)= 16.36, P<0.0001<br>$F_{sex,}$ P=0.2950 (NS)<br>$F_{treatment*sex,}$ P=0.1932 (NS) | 5-6 |
| Fig 4.E2. Anhedonia, | 2 way ANOVA, treatment*sex | $F_{treatment}$(1,18)= 6.255, P=0.0223<br>$F_{sex,}$ P= NS<br>$F_{treatment*sex,}$ P=NS | 5-6 |

| | | | |
|---|---|---|---|
| weighted AVG | | | |
| Fig 4.F. OFT 2-3month | 2 way ANOVA, treatment*sex | NS | 5-6 |
| Fig 4.G Correlation matrix, 3M | Pearson's correlation analysis | WB/VF: r = 0.471, P=0.031.<br>WB/OFT: r = - 0.048, P=0.84 (NS).<br>WB/SPT: r = 0.656, P=0.001.<br>VF/OFT: r = - 0.024, P=0.92. (NS)<br>VF/SPT: r = 0.651, P=0.001.<br>OFT/SPT: r = 0.197, P=0.38. (NS) | across groups Total: 21$ |
| **Fig 5. Treatment at disease onset - sensory** | | | |
| Fig 5.A1. VF | 3 way RM ANOVA, treatment*sex*time | $F_{treatment}$ (1,20)= 82.47, P<0.0001<br>$F_{time}$ (19,380)= 67.77, P<0.0001<br>$F_{time*treatment}$(19,380)= 4.729, P<0.0001 | 6 |
| Fig 5.A2. VF AVG | 2 way ANOVA, treatment*sex | $F_{treatment}$ (1,20)= 50.50, P<0.0001 | 6 |
| Fig 5.B1. WB | 3 way RM ANOVA, treatment*sex*time | $F_{treatment}$ (1,20)= 27.96, P<0.0001<br>$F_{sex}$ (1,20)= 7.658, P=0.0119<br>$F_{time}$ (20,400)= 100.2, P<0.0001<br>$F_{time*sex}$ (20,400)= 2.232, P=0.0019<br>$F_{time*treatment}$(20,400)= 5.824, P<0.0001<br>$F_{sex*treatment}$= P=0.54 (NS) | 6 |
| Fig 5.B2, WB-AVG | 2 way ANOVA, treatment*sex | $F_{treatment}$ (1,20)= 119.6, P<0.0001<br>$F_{sex}$ (1,20)= 11.73, P=0.0027<br>$F_{sex*treatment}$ = P=0.94 (NS) | 6 |
| Fig 5C. Guarding index, 6 weeks catwalk | 2 way ANOVA, treatment*sex | $F_{treatment}$ (1,16)= 9.316, P=0.0076 | 4-6 |
| Fig 5D1. SPT | 3 way RM ANOVA, time*treatment*sex | $F_{treatment}$(1,20)= 22.75, P=0.0001 | 6 |
| Fig 5.D2. SPT weighted AVG | 2 way ANOVA, treatment*sex | $F_{treatment}$ (1,20)= 22.75, P=0.0001 | 6 |
| Fig 5E. OFT 3M | 2 way ANOVA, treatment*sex | $F_{treatment}$ (1,20)= 4.452, P=0.0477 | 6 |
| Fig 5F1. Correlation matrix, 3M | Pearson's correlation analysis | WB/VF: r = 0.704, P<0.0001.<br>WB/OFT: r = 0.413, P=0.045.<br>WB/SPT: r = 0.595, P=0.002.<br>VF/OFT: r = 0.300, P=0.15. (NS)<br>VF/SPT: r = 0.659, P<0.0001.<br>OFT/SPT: r = 0.321, P=0.126. (NS) | 24, across groups |
| Fig 5F2. Correlation matrix, 4.5M | | WB/VF: r = 0.641, P=0.001.<br>WB/OFT: r = 0.112, P=0.60. (NS)<br>WB/SPT: r = 0.468, P=0.021.<br>VF/OFT: r = 0.076, P=0.72. (NS)<br>VF/SPT: r = 0.435, P=0.034.<br>OFT/SPT: r = 0.389, P=0.06. (NS) | |
| Fig 5F3. Correlation matrix, 6M. | | WB/VF: r = 0.206, P=0.33. (NS)<br>WB/OFT: r = 0.079, P=0.72. (NS)<br>WB/SPT: r = - 0.030, P=0.89. (NS)<br>VF/OFT: r = 0.114, P=0.60. (NS)<br>VF/SPT: r = - 0.105, P=0.63. (NS)<br>OFT/SPT: r = 0.186, P=0.38. (NS) | |
| Fig 5G1. Prediction. | Simple linear regression, and | r = 0.4242, F(1,22)=4.828, P=0.0388 | 24 |

| | | | | |
|---|---|---|---|---|
| OFT-3M vs WB 2W | Pearson's correlation analysis | | | |
| Fig 5G2. Prediction. OFT-3M vs VF 2W | Simple linear regression, and Pearson's correlation analysis | r = 0.4488, F(1,22)=5.548, P=0.028 | | 24 |
| Fig 5H1. Prediction. SPT-3M vs WB 2W | Simple linear regression, and Pearson's correlation analysis | r = 0.5915, F(1,22)=11.84, P=0.0023 | | 24 |
| Fig 5H2. Prediction. SPT-3M vs VF 2W | Simple linear regression, and Pearson's correlation analysis | r = 0.6350, F(1,22)=14.87, P=0.0009 | | 24 |
| Fig 5I | 3-way RM ANOVA<br>Sex<br>Treatment<br>Time | All time points:<br>Sex: F(1,12)=48.5; P<0.001<br><br>ZT14 to ZT01:<br>Sex: F(1,12)=33.7, P<0.001<br><br>ZT02 to ZT04:<br>Treat: F(1,12)=12.4, P=0.004<br>Sex: F(1,12)=78; P<0.001;<br>Sex x Treatment: F(1,12)=8.3; P=0.014<br><br>ZT16 to ZT17<br>Treat: F(1,12)=5.1, P=0.044<br>Sex: F(1,12)=58.5; P<0.001 | 2-way RM for females:<br><br>ZT02 to ZT04: Females only: Treat: F(1,6)=16; P=0.007<br><br>ZT16 to ZT17 Females only: Treat: NS | 4 |
| Fig 5J | 3-way RM ANOVA<br>Sex<br>Treatment<br>Time | All time points:<br>Sex: F(1,12)=25.9; P<0.001<br>Treat: F(1,12)=3.6, P=0.082<br><br>ZT14 to ZT01:<br>Sex: F(1,12)=25.6, P<0.001<br>Treat: F(1,12)=2.5, P=0.139<br><br>ZT02 to ZT04:<br>Sex: F(1,12)=22.1; P<0.001;<br>Treat: F(1,12)=7.6, P=0.017<br><br>ZT13 to ZT19:<br>Sex: F(1,12)=28.9; P<0.001<br>Treat: F(1,12)=4.9, P=0.046 | 2-way RM:<br><br>ZT02 to ZT04: Females only: Treat: NS<br><br>ZT02 to ZT03: Females only: F(1,6)=10.1; P=0.019<br><br>ZT13 to ZT19: Females only: NS | 4 |

| | | | |
|---|---|---|---|
| | | ZT16 to ZT19 Females only: NS | |
| Fig 5K | 2-way ANOVA<br>Sex<br>Treatment | L1: sex: F(1,12)= 17, P=0.001<br>L2: sex: F(1,12)=19, P<0.001<br>L3: sex: F(1,12)=21, P<0.001<br>L4: sex: F(1,12)=7.8; P=0.016<br>Treatment: F(1,12)=6.1, P=0.03<br>L5: sex: F(1,12)=7.7; P=0.07<br>L6: F(1,12)=25, P<0.001 | 4 |
| **Fig 6. FKPB51 early vs late – RNA sequencing** | | | |
| Fig 6A1. VF | 3 way RM ANOVA, treatment*timing*time | $F_{treatment}$(1,26)= 63.00, P<0.0001<br>$F_{time}$(8,208)= 158.3, P<0.0001<br>$F_{time*timing}$(8,208)= 1.748, P=0.0871 (NS)<br>$F_{time*treatment}$(8,208)= 7.960, P<0.0001<br>$F_{timing*treatment}$(1,26)= 8.682, P=0.0067<br>$F_{time*timing*treatment}$(8,208)= 2.467, P=0.0142 | 7-8 |
| Fig 6A2. VF AVG. | 3 way RM ANOVA, treatment*timing*phase | $F_{treatment}$(1,26)= 82.86, P<0.0001<br>$F_{phase}$(1,26)= 8.248, P=0.008<br>$F_{timing}$(1,26)= 5.895, P=0.0224<br>$F_{phase*timing}$(1,26)= 5.123, P=0.0322<br>$F_{phase*treatment}$(1,26)= 25.53, P<0.0001<br>$F_{timing*treatment}$(1,26)= 10.18, P=0.0037<br>$F_{timing*phase*treatment}$(1,26)= 9.267, P=0.0053 | 7-8 |
| Fig 6B. guarding index, D23 | 2 way ANOVA, Treatment*timing | $F_{treatment}$(1,26)= 4.188, P=0.05<br>$F_{timing}$(1,26)= 2.801, P=0.1062 (NS)<br>$F_{interaction}$(1,26)= 2.494, P=0.1263 (NS) | 7-8 |
| Fig 6C, corticosterone D31 | 2 way ANOVA, Treatment*timing | $F_{treatment}$(1,26)= 6.156, P=0.0199<br>$F_{timing}$(1,26)= 4.112, P=0.0530 (NS) | 7-8 |
| Fig 6D. cort vs VF correlation | Simple linear regression and Pearson's correlation analysis | r = -0.4058, F(1,28)=5.521, P=0.0261 | 30 |
| Fig 6G *Fkbp5* seq | See Sequencing analysis | | |
| Fig 6J1. Shank 3 | Left panel: result of sequencing nested analysis of Late SAFit2 vs Late Vehicle Right Panel: Pearson's correlation analysis with Bonferroni correction | Left: F=6.03, P<0.05;<br>Right: r2=0.75; P=0.00012, t=-7.8, dF=20 | 5-6 |
| Fig 6J2. Pten | Left panel: result of sequencing nested analysis of Late SAFit2 vs Late Vehicle Right Panel: Pearson's correlation | Left: F=9.16, P<0.05; Right: r2=0.76; P=0.00010, t=7.93, dF=20 | 5-6 |

| | | | |
|---|---|---|---|
| | analysis with Bonferroni correction | | |
| Fig 6J3. Nfkbib | Left panel: result of sequencing nested analysis of Late SAFit2 vs Late Vehicle Right Panel: Pearson's correlation analysis with Bonferroni correction | Left: F=7.91, P<0.05; Right: r2=0.76; P=0.000098, t=-7.94, dF=20 | 5-6 |
| Fig 6L | Result of t-test Early SAFit2 vs Early Vehicle | See full statistical method in RNAseq section. | |

[#] missing values due to different duration of experimental cohorts. All cohorts included equal numbers of animals from the different groups. Weighted average was calculated based on the period that the animals in question was tested.

[$]one animal removed from this outcome, as it was not complying with WB-training before treatment-initiation (Fig 4B), or developed a contralateral injury (broken nail) during the experiment, rendering the result unreliable (Fig 4D).

RM= Repeated Measures. NS = not significant.

**Table S3 Complete information for statistical analysis of figures from Supplementary Data.**

| Fig | Analysis | F-values | N |
|---|---|---|---|
| **Fig S1. Characterisation** | | | |
| Fig S1A. Cort | 2 way ANOVA, injury*sex | NS | 8 |
| Fig S1B1. Affective-motivati onal, low | 3 way RM ANOVA, injury*sex*time | $F_{injury}$ (1,18)= 36.28, P<0.0001<br>$F_{time}$ (10,180)= 2.821, P=0.0028<br>$F_{time*injury}$ (10,180)= 3.029, P=0.0014<br>$F_{time*sex}$ (10,180)= 3.363, P=0.0005<br>$F_{sex*injury}$ (1,18)= 4.601, P=0.0459<br>$F_{sex*injury*time}$ (10,180)= 2.932, P=0.002. | |
| Fig S1.B2. Low – weighted AVG | 2 way ANOVA, injury*sex | $F_{injury}$ (1,18)= 27.31, P<0.0001 | |
| Fig S1C1. Affective-motivational, medium (0.16g) | 3 way RM ANOVA, injury*sex*time | $F_{injury}$ (1,18)= 74.02, P<0.0001<br>$F_{time}$ (10,180)= 3.500, P=0.0003<br>$F_{time*injury}$ (10,180)= 4.068, P<0.0001<br>$F_{sex*injury}$ (1,18)= 8.092, P=0.0108 | 4-6 |
| Fig S1C2. medium – weighted AVG | 2 way ANOVA, injury*sex | $F_{injury}$ (1,18)= 55.09, P<0.0001<br>$F_{sex*injury}$ (1,18)= 4.572, P=0.0465 | 4-6 |
| Fig S1D1. Affective-motivational, high (1.0g) | 3 way RM ANOVA, injury*sex*time | $F_{injury}$ (1,18)= 83.70, P<0.0001<br>$F_{time}$ (10,180)= 6.048, P<0.0001<br>$F_{sex*}$ (1,18)= 13.70, P<0.0001<br>$F_{time*injury}$ (10,180)= 5.417, P<0.0001<br>$F_{sex*injury}$ (1,18)= 17.97, P=0.0005 | 4-6 |
| Fig S1D2. High – weighted AVG | 2 way ANOVA, injury*sex | $F_{injury}$ (1,18)= 67.70, P<0.0001<br>$F_{sex}$ (1,18)= 14.95, P=0.0011<br>$F_{sex*injury}$ (1,18)= 16.59, P=0.0007 | 4-6 |
| Fig S1E. VF 2W AVG vs 6M OFT | Simple linear regression, and Pearson's correlation analysis | r = 0.3568, F(1,21)=3.063, P=0.0947 (NS) | |
| **Fig S2. Activity Supplementary Naive Baseline** | | | |
| Fig S2 | Inter-daily Stability Sleep<br>Student's T test | P<0.005 | 8 |
| **Fig S3. Activity Supplementary Post-MIA 1** | | | |
| Fig S3A | 3 way RM ANOVA injury*sex*time | **ZT14 to ZT24**: sex: F(1,12)=15.5; P=0.002 | 4 |
| Fig S3B | 3 way RM ANOVA injury*sex*time | **ZT14 to ZT24**: sex: F(1,12)=5.1; P=0.043 | 4 |
| Fig S3C | 3 way RM ANOVA injury*sex*time | **ZT14 to ZT24**: sex: F(1,12)=11.7; P=0.005 | 4 |
| Fig S3D | 3 way RM ANOVA injury*sex*time | **ZT14 to ZT24**: sex: F(1,12)=18.6; P=0.001 | 4 |
| Fig S3E | 3 way RM ANOVA injury*sex*time | **ZT14 to ZT24**: sex: F(1,12)=8.04; P=0.015 | 4 |
| Fig S3F | 3 way RM ANOVA injury*sex*time | **ZT14 to ZT24**: sex: F(1,12)=7.8; P=0.016 | 4 |
| Fig S3G | 3 way RM ANOVA injury*sex*time | **ZT14 to ZT24**: sex: F(1,12)=4.2; P=0.063 | 4 |

| | | | | |
|---|---|---|---|---|
| Fig S3H | 3 way RM ANOVA injury*sex*time | **ZT14 to ZT24**: sex: F(1,12)=6.8; P=0.023 | | 4 |
| **Fig S4. Activity Supplementary Post-MIA 2** | | | | |
| Fig S4 A-L | 2-Way ANOVA at each single time point as not same animals throughout | i: P<0.05 injury<br>s: P<0.05 sex<br>x: P<0.05 sex x injury | | 4 |
| **Fig S5. Effects of genetic knockout of FKBP51 on MIA-induced behavior.** | | | | |
| Fig S5A1. BW | Mixed-effects model[#], genotype*sex*time | $F_{genotype}$ (1,18)= 17.86, P=0.0005<br>$F_{sex}$ (1,18)= 91.93, P<0.0001<br>$F_{time}$ (16,286)= 77.90, P<0.0001<br>$F_{time*sex}$ (16,286)= 9.466, P<0.0001<br>$F_{time*genotype}$(16,286)= 1.725, P=0.0416<br>$F_{sex*genotype}$= P=0.54 (NS) | | 5-6 |
| Fig S5A2. BW - AVG | 2 way ANOVA, genotype*sex | $F_{sex}$ (1,18)= 51.69, P<0.0001<br>$F_{genotype}$ (1,18)= 13.86, P=0.0016 | | 5-6 |
| Fig S5B. Bone volume, non-transformed | 2 way ANOVA, genotype*sex | $F_{sex}$ (1,16)= 20.86, P=0.0003<br>$F_{genotype}$ (1,16)= 13.65, P=0.0020 | | 4-6 |
| Fig S5D. OFT. Distance traveled | 3 way RM ANOVA, genotype*sex*time | $F_{sex}$ (1,18)= 8.574, P=0.0090<br>$F_{genotype}$ (1,18)= 9.245, P=0.0070<br>$F_{time*sex}$ (4,72)= 2.129, P=0.0860 (NS)<br>$F_{time*genotype}$(4,72)= 2.399, P=0.0579 (NS) | | 5-6 |
| Fig S5D. EPM. Time in open | Mixed-effects model[#], genotype*sex*time | NS | | 4-6 |
| **Fig S6. Transgenic modification, immobility/activity** | | | | |
| Fig S6A | 3-way RM ANOVA<br>Sex<br>Genotype<br>Time | NS | | 4 |
| Fig S6B | 3-way RM ANOVA<br>Sex<br>Genotype<br>Time | NS | | 4 |
| Fig S6C | 3-way RM ANOVA<br>Sex<br>Genotype<br>Time | All time course:<br>Sex: F(1,12)=4.9; P= 0.047;<br><br>ZT14 to ZT17<br>Gene: F(1,12)=11.3; P=0.006;<br>Sex: F(1,12)=6.6; P=0.024. | 2-way RM:<br>Males:<br>21:00 to 24:00:<br>Gene: F(1,6)=7.8; P=0.032;<br>Females: NS | 4 |
| Fig S6D | 3-way RM ANOVA<br>Sex<br>Genotype<br>Time | All time course:<br>Gene: F(1,12)=4.75; P=0.050;<br>Sex: F(1,12)=3.6; P= 0.081;<br><br>ZT14 to ZT1<br>Gene: F(1,12)=9.6, P=0.009<br>Sex: F(1,12)=3.4; p=0.090<br><br>ZT14 to ZT23<br>Gene: F(1,12)=9.8; P=0.009;<br>Sex: F(1,12)=4.5; P= 0.056; | 2-way RM:<br>Males:<br>21:00 to 01:00:<br>Gene: F(1,6)=8.8; P=0.025;<br>Females: NS | 4 |

| | | | | |
|---|---|---|---|---|
| | | ZT14to ZT18:<br>Gene: F(1,12)=9.5; P=0.009;<br>Sex: F(1,12)=5.0; P= 0.045; | | |
| Fig S6E | 2-way ANOVA<br>Sex<br>Genotype | No significance of genotype; sex significance:<br>E3: F(1,12)=5.3; P=0.040; E6: F(1,12)=5.7;<br>P=0.034 | | 4 |
| Fig S6F | 2-way ANOVA<br>Sex<br>Genotype | No significance of sex; genotype significance:<br>F1: F(1,12)=5.6; P=0.036; F3: F(1,12)=4.4;<br>P=0.058; F(1,12)=10.2; P=0.008; F(1,12)=6.5;<br>P=0.025; F(1,12)=6.7; P=0.024. | | 4 |
| **Fig S7. Activity Supplementary KO/WT 1** | | | | |
| Fig S7 | 2-way ANOVA | D: Interactions Gene x Sex: F(1,12)=5.5; P=0.037 | | 4 |
| **Fig S8. Activity Supplementary KO/WT 2** | | | | |
| Fig S8A | 3-way RM ANOVA | ZT16 to ZT23:<br>Gene: F(1,12)=4.6; P=0.053;<br>Sex: F(1,12)=23.5; P<0.001;<br>Gene x sex: F(1,12)=4.6; P=0.052 | 2-way RM:<br>Females:<br>23:00 to<br>06:00:<br>Gene:<br>F(1,6)=7.9;<br>P=0.031;<br>Males: NS | 4 |
| Fig S8B | 3-way RM ANOVA | ZT16 to ZT23:<br>Gene: F(1,12)=4.8; P=0.048;<br>Sex: F(1,12)=36.2; P<0.001;<br>Gene x sex: F(1,12)=2.8, P=0.117. | 2-way RM:<br>Females:<br>02:00 to<br>06:00:<br>Gene:<br>F(1,6)=6.5;<br>P=0.044;<br>Males: NS | 4 |
| Fig S8C | 3-way RM ANOVA | ZT18 to ZT23:<br>Sex: F(1,12)=9.5; P=0.010; | | 4 |
| Fig S8D | 3-way RM ANOVA | ZT18 to ZT23:<br>Sex: F(1,12)=13.1; P=0.004; | | 4 |
| Fig S8E | 3-way RM ANOVA | NS across the whole time course | | 4 |
| Fig S8F | 3-way RM ANOVA | ZT18 to ZT21:<br>Sex: F(1,12)=4.2; P=0.063 | | 4 |
| **Fig S9. Activity Supplementary KO/WT 3** | | | | |
| Fig S9 A to L | 2-Way ANOVA at each single time point as not same animals throughout | g: P<0.05 genotype<br>s: P<0.05 sex<br>x: P<0.05 sex x genotype | | 4 |
| **Fig S10. FKBP51 inhibition after injury** | | | | |
| Fig S10A. VF.<br>2 week study | 2 way RM ANOVA, treatment*time | $F_{time}$ (7,91)= 92.88, P<0.0001<br>$F_{treatment}$ (1,13)= 12.89, P=0.0033<br>$F_{treatment*time}$ (7,91)= 10.31, P<0.0001 | | 7-8 |
| Fig S10B. WB.<br>2 week study | 2 way RM ANOVA, treatment*time | $F_{time}$ (7,91)= 62.50, P<0.0001<br>$F_{treatment}$ (1,13)= 11.16, P=0.0053<br>$F_{treatment*time}$ (7,91)= 6.913, P<0.0001 | | 7-8 |
| Fig S10C. EPM. Time in open<br>2-3M study | 3-way RM ANOVA, treatment*sex*time | $F_{time}$ (1,17)= 7.921, P=0.0119 | | 5-6 |

| | | | |
|---|---|---|---|
| Fig S10D. OFT Time in center 2-3M study | 3-way RM ANOVA, treatment*sex*time | $F_{time}$, P=0.0797 (NS) | 5-6 |
| Fig S10E. OFT - Distance travelled. 2-3M study | 2 way ANOVA, treatment*sex | NS | 5-6 |
| Fig S10F. BW. 2-3M study | 3 way RM ANOVA, treatment*sex*time | $F_{sex}$ (1,18)= 214.4, P<0.0001<br>$F_{time}$ (5,90)= 246.2, P<0.0001<br>$F_{time*sex}$ (5,90)= 12.61, P<0.0001<br>$F_{time*treatment}$ (5,90)= 3.611, P=0.0051 | 5-6 |
| Fig S10.G1 WB-active/SPT D120. | Simple linear regression, and Pearson's correlation analysis | Female; P=0.2099 = NS.<br>Male; r=0.7675, F(1,8)=11.46, P=0.0096 | |
| Fig S10.G2 VF-active/SPT D120. | Simple linear regression, and Pearson's correlation analysis | Female; r=0.6731, F(1,9)=7.454, P=0.023<br>Male; P= 0.5523 (NS) | |
| Fig S10.H1 WB-active/SPT D140. | Simple linear regression, and Pearson's correlation analysis | Female; P=0.1108 = (NS).<br>Male; P=0.7173 (NS) | |
| Fig S10.H2 VF-active/SPT D140. | Simple linear regression, and Pearson's correlation analysis | Female; P=0.3697 = (NS).<br>Male; P=0.1511 (NS) | |
| **Fig S11. FKBP51 inhibition in naïve controls.** | | | |
| Fig S11A. OFT | 2 way ANOVA, treatment*sex | $F_{sex}$ (1,12)= 9.895, P=0.0084<br>$F_{treatment}$ (1,12)= 9.410, P=0.0098 | 4 |
| Fig S11B. VF | 3 way RM ANOVA, treatment*sex*time | NS | 4 |
| Fig S11C. SPT – 1% | 2 way ANOVA, treatment*sex | NS | 4 |
| Fig S11D. SPT - 0.5% | 2 way ANOVA, treatment*sex | NS | 4 |
| **Fig S12. FKBP51 inhibition at onset. Catwalk at 6weeks** | | | |
| Fig S12A. Dynamic WB | 2 way ANOVA, treatment*sex | $F_{treatment}$ (1,16)= 9.518, P=0.0071 | 4-6 |
| Fig S12B. Print area | 2 way ANOVA, treatment*sex | $F_{treatment}$ (1,16)= 8.810, P=0.0091 | 4-6 |
| Fig S12C. Print intensity | 2 way ANOVA, treatment*sex | $F_{treatment}$ (1,16)= 5.741, P=0.0291 | 4-6 |
| Fig S12D. Run speed | 2 way ANOVA, treatment*sex | NS | 4-6 |
| Fig S12E. %MPE - VF | 2way ANOVA, Treatment*timing | $F_{treatment}$ (1,77)= 50.52, P<0.0001 | 7-12, sex-combined |

| | | | |
|---|---|---|---|
| Fig S12F. %MPE - WB | 2way ANOVA, Treatment*timing | $F_{treatment}$ (1,75)= 95.26, P<0.0001<br>$F_{treatment*timing}$ (3,75)= 4.212, P=0.0083 | 7-12, sex-combined |
| Fig S12G. Bone volume (corrected for BW) | 2 way ANOVA, treatment*sex | $F_{sex}$ (1,20)= 4.618, P=0.0441 | 6 |
| **Fig S13. FKBP51 inhibition at onset. Affective etc** | | | |
| Fig S13A1. Affective low | 3 way RM ANOVA, time*treatment*sex | $F_{treatment}$(1,20)= 29.48, P<0.0001 | 6 |
| Fig S13A2. Affective low, AVG | 2 way ANOVA, treatment*sex | $F_{treatment}$ (1,20)= 32.71, P<0.0001 | 6 |
| Fig S13B1. Affective medium | 3 way RM ANOVA, treatment*sex*time | $F_{treatment}$ (1,20)= 12.53, P=0.0021 | 6 |
| Fig S13B2. AVG affective medium | 2 way ANOVA, treatment*sex | $F_{treatment}$ (1,20)= 12.19, P=0.0023 | 6 |
| Fig S13C1. Affective high | 3 way RM ANOVA, treatment*sex*time | $F_{treatment}$ (1,20)= 7.925, P=0.0107<br>$F_{time}$(8,160)= 4.900, P<0.0001 | 6 |
| Fig S13C2. AVG affective high | 2 way ANOVA, treatment*sex | $F_{treatment}$ (1,20)= 8.041, P=0.0102 | 6 |
| Fig S13D. brush | 2 way RM ANOVA, treatment*time | $F_{time}$ (3,30)= 4.005, P=0.0164<br>$F_{treatment}$(1,10)= 2.811, P=0.1246 (NS)<br>$F_{timer*treatment}$ (3.30)= 4.367, P=0.0115 | 6 |
| Fig S13E. OFT - repeated | 3 way RM ANOVA, treatment*sex*time | $F_{time}$(3,60)= 12.33, P<0.0001<br>$F_{treatment}$(1,20)= 3.369, P=0.0814 (NS) | 6 |
| Fig S13F. BW | 3 way RM ANOVA, treatment*sex*time | $F_{sex}$ (1,20)= 156.7, P<0.0001<br>$F_{time}$(6,120)= 220.9, P<0.0001<br>$F_{time*sex}$(6,120)= 3.769, P=0.0018 | 6 |
| Fig S13G. OFT distance | 3 way RM ANOVA, treatment*sex*time | $F_{sex}$ (1,20)= 4.121, P=0.0559 (NS)<br>$F_{time}$(3,60)= 26.10, P<0.0001<br>$F_{time*sex}$(3,60)= 2.440, P=0.0731 (NS) | 6 |
| **Fig S14. Activity Inhibition at onset** | | | |
| Fig S14 A | 3 way RM ANOVA, treatment*sex*time | ZT14 to ZT19:<br>Sex: F(1,12)=46.5; P<0.001.<br>Treat: F(1,12)=4.6, P=0.054<br><br>ZT14 to ZT17:<br>Sex: F(1,12)=47.4, P<0.001<br>Treat: F(1,12)=5.5, P=0.037 | 4 |
| Fig S14 B | 3 way RM ANOVA, treatment*sex*time | ZT13 to ZT23<br>Sex<br>F(1,12)=25.8, P<0.001 | 4 |
| Fig S14 C | 3 way RM ANOVA, treatment*sex*time | ZT14 to ZT21<br>Sex:<br>F(1,12)=26.0, P<0.001 | 4 |
| Fig S14 D | 3 way RM ANOVA, treatment*sex*time | ZT15 to ZT23:<br>Sex: F(1,12)=12.5, P=0.004 | 4 |

| | | | |
|---|---|---|---|
| Fig S14 E | 2-way ANOVA at individual time points as not all animals were measured at every time points | T; treatment effect<br>S: treatment effect<br>s: P<0.05 | 4 |
| **Fig S15. Activity SAFit alone** | | | |
| Fig S15 A | 3-way RM ANOVA<br>Sex<br>Treatment<br>Time | All time points:<br>Sex: F(1,12)=3.7; P= 0.079;<br>Sex x Treatment: F(1,12)=3.6; P=0.083<br><br>**ZT14 to ZT01**:<br>Sex: F(1,12)=3.4, P=0.091<br>Sex x treatment: F(1,12)=4.5, P=0.054<br><br>**ZT14 to ZT20**:<br>Sex: F(1,12)=12.5; P=0.004;<br>Sex x Treatment: F(1,12)=6.6; P=0.024. | 4 |
| Fig S15 B | 3-way RM ANOVA<br>Sex<br>Treatment<br>Time | All time points:<br>Sex: F(1,12)=6.1; P= 0.030;<br>Sex x Treatment: F(1,12)=5.5; P=0.037<br><br>**ZT14 to ZT01:**<br>Sex: F(1,12)=5.1, P=0.043<br>Sex x treat: F(1,12)=5.05, P=0.044<br><br>**ZT14 to ZT20**:<br>Sex: F(1,12)=7.1; P=0.021;<br>Sex x Treatment: F(1,12)=4.9; P=0.047. | 4 |
| Fig S15 C to N | 2-way ANOVA<br>Sex x treat | No treatment significance.<br>Sex:<br>C: NS<br>D: F(1,3)=5.8; P=0.033<br>E: F(1,3)=6.5; P=0.026<br>F: NS<br>G: F(1,3)=9.4; P=0.010<br>H: F(1,3)=6.6; P=0.024<br>I: F(1,3)=4.2; P=0.064<br>J: NS<br>K: NS<br>L: NS<br>M:NS<br>N: NS | 4 |
| **Fig S16. RNA seq supplementary** | | | |
| Fig S16A. Dynamic WB. Catwalk D23 | 2 way ANOVA, treatment*timing | $F_{treatment}$=(1,26)=4.795, P=0.0377<br>$F_{timing}$=(1,26)=2.790, P=0.1069 (NS) | 7-8 |
| Fig S16B. Print area. Catwalk D23 | 2 way ANOVA, treatment*timing | $F_{timing}$=(1,26)=3.193, P=0.0856 (NS) | 7-8 |
| Fig S16C. Mean intensity. Catwalk D23 | 2 way ANOVA, treatment*timing | $F_{treatment}$=(1,26)=4.597, P=0.0416 | 7-8 |
| Dig S16D. Run speed. D23. | 2 way ANOVA, treatment*timing | NS | 7-8 |

| Fig S18. Sequencing data | | | |
|---|---|---|---|
| A. Kcnq2 | Left panel: result of sequencing nested analysis of Late SAFit2 vs Late Vehicle Right Panel: Pearson's correlation analysis with Bonferroni correction | Left: P<0.05; Right: r2=0.75; P=0.000148, t=-7.7, dF=20 | 5-6 |
| B. Lrp1 | Left panel: result of sequencing nested analysis of Late SAFit2 vs Late VehicleRight Panel: Pearson's correlation analysis with Bonferroni correction | Left: P<0.05; Right: r2=0.64; P=0.0071, t=-5.87, dF=20 | 5-6 |
| C. Maoa | Left panel: result of sequencing nested analysis of Late SAFit2 vs Late VehicleRight Panel: Pearson's correlation analysis with Bonferroni correction | Left: P<0.05; Right: r2=0.69; P=0.0014, t=6.62, dF=20 | 5-6 |
| D. P2ry | Left panel: result of sequencing nested analysis of Late SAFit2 vs Late VehicleRight Panel: Pearson's correlation analysis with Bonferroni correction | Left: P<0.05; Right: r2=0.66; P=0.0039, t=6.15, dF=20 | 5-6 |
| E. Rap1a | Left panel: result of sequencing nested analysis of Late SAFit2 vs Late VehicleRight Panel: Pearson's correlation analysis with Bonferroni correction | Left: P<0.05; Right: r2=0.66; P=0.0037, t=6.18, dF=20 | 5-6 |
| F. Cox2 | Left panel: result of sequencing nested analysis of Late SAFit2 vs Late VehicleRight Panel: Pearson's correlation analysis with | Left: P<0.05; Right: r2=0.81; P<0.0001, t=9.22, dF=20 | 5-6 |

| | |
|---|---|
| | Bonferroni correction |
| Fig S19. Statistical analysis discussed in the sequencing section | |

RM= Repeated Measures. NS = not significant.

**Table S4. Percentage of mice "affected" by emotional comorbidities associated with chronic joint pain (data presented in Fig 1)**

| % Affected mice | Depressive-like behaviour | | | | Anxiety-like behaviour | |
|---|---|---|---|---|---|---|
| **Time (months)** | 3M | 4M | 5M | 6M | 3M | 6M |
| **MIA Females** | 54.5 | 0 | 80 | 100 | 45 | 80 |
| **MIA Males** | 25 | 75 | 25 | 42.9 | 27 | 43 |
| **Control Females** | 16.7 | 16.7 | 16.7 | 16.7 | 8.3 | 33.3 |
| **Control Males** | 20 | 0 | 25 | 20 | 9.1 | 20 |

**Table S5. Early outcomes can be used as predictors of late symptoms.** Values are Pearson’s r. *P<0.05. #P=0.05-0.1

| Predictions | | OFT | | | SPT | | | VF | | | WB | | |
|---|---|---|---|---|---|---|---|---|---|---|---|---|---|
| Exp. | AVG | 3M | 4.5M | 6M | 3M | 4.5M | 6M | 3M | 4.5M | 6M | 3M | 4.5M | 6M |
| Characterisation Ctrl/ MIA (Fig.1) | WB-2W | * 0.52 | | | 0.35 | | | * 0.96 | | | * 0.99 | | |
| | VF-2W | * 0.53 | | # 0.36 | * 0.34 | | * 0.59 | * 0.93 | | * 0.86 | * 0.96 | | |
| KO/ WT (Fig.3) | WB-2W | 0.28 | 0.28 | -0.26 | 0.34 | 0.15 | 0.19 | * 0.65 | * 0.73 | 0.81 | * 0.72 | * 0.85 | * 0.78 |
| | VF-2W | 0.11 | 0.22 | 0.20 | # 0.39 | 0.28 | 0.34 | * 0.71 | * 0.79 | 0.79 | * 0.69 | * 0.78 | * 0.59 |
| SAFit2-VPG (2-3M) (Fig.4 C to F) | WB-active | 0.26 | | | * 0.56 | # 0.41 | | * 0.59 | - 0.3 | | * 0.79 | 0.26 | |
| | VF-active | 0.09 | | | 0.34 | # 0.38 | | * 0.89 | -0.01 | | * 0.45 | 0.21 | |
| | WB-2W | 0.35 | | | 0.09 | -0.13 | | -0.16 | 0.12 | | 0.26 | 0.29 | |
| | VF-2W | -0.24 | | | 0.21 | * 0.44 | | 0.14 | 0.34 | | -0.06 | 0.22 | |
| SAFit2-VPG Prevention (Fig.5) | WB-2W | * 0.42 | 0.22 | 0.24 | * 0.59 | * 0.63 | 0.33 | * 0.76 | * 0.66 | * 0.44 | * 0.67 | * 0.64 | # 0.35 |
| | VF-2W | * 0.45 | 0.21 | 0.06 | * 0.64 | * 0.67 | 0.23 | * 0.90 | * 0.67 | * 0.73 | * 0.77 | * 0.68 | 0.26 |

**Table S6. Percentage of mice "resilient" to emotional behaviour associated with chronic joint pain (*Fkbp5* KO and WT) (data presented in Fig 3).**

| % Resilient mice | Depressive-like behaviour | | | | Anxiety-like behaviour | | | |
|---|---|---|---|---|---|---|---|---|
| Time (months) | 0M | 3M | 4.5M | 6M | 0M | 3M | 4.5M | 6M |
| WT Females | 16.7 | 12.5 | 12.5 | 11.1 | 16.7 | 16.7 | 16.7 | 25 |
| WT Males | 20 | 14.3 | 14.3 | 12.5 | 20 | 20 | 20 | 16.7 |
| KO Females | 0 | 37.5 | 25 | 44.4 | 20 | 20 | 40 | 12.5 |
| KO Males | 0 | 50 | 50 | 55.6 | 33.3 | 16.7 | 16.7 | 28.6 |

**Table S7. Percentage of mice "resilient" to emotional behaviours associated with chronic joint pain following FKBP51 inhibition 2 months after injury (data presented in Fig 4C-F). "**Active" = while the compound was active, "Post" = 3 weeks after the end of the active period, "Late" = 6 weeks after end of active period.

| % Resilient mice | Depressive-like behaviour | | | Anxiety-like behaviour | |
|---|---|---|---|---|---|
| **Time** | Active | Post | Late | Active | Post |
| **Veh Females** | 16.7 | 16.7 | 33.3 | 16.7 | 16.7 |
| **Veh Males** | 20 | 0 | 20 | 20 | 0 |
| **SAFit2 Females** | 40 | 60 | 40 | 20 | 20 |
| **SAFit2 Males** | 50 | 83.3 | 33.3 | 0 | 16.7 |

**Table S8. Percentage of mice "resilient" to emotional behaviours associated with chronic joint pain following FKBP51 inhibition during disease onset (data presented in Fig 5).**

| % Resilient mice | Depressive-like behaviour | | | Anxiety-like behaviour | | |
|---|---|---|---|---|---|---|
| **Time (months)** | 3M | 4.5M | 6M | 3M | 4.5M | 6M |
| **Veh Females** | 0 | 16.7 | 16.7 | 16.7 | 33.3 | 16.7 |
| **Veh Males** | 16.7 | 16.7 | 0 | 0 | 50 | 16.7 |
| **SAFit2 Females** | 83.3 | 100 | 83.3 | 16.7 | 50 | 0 |
| **SAFit2 Males** | 66.7 | 33.3 | 16.7 | 50 | 33.3 | 16.7 |

**Table S9 DEGs in MIA *vs* Control animals at day 30 and their modulation by SAFit2 (Fig 6).** In a separate study, we identified MIA-modified genes in the ipsilateral spinal cord 30 days after MIA injection. When we compared those identified DEGs (n=704, P<0.05) with DEGs modulated by SAFit2, we found only 37 common DEGs; SAFit2 reversed the MIA-induced changes in expression in 11 of those. Data shows normalised counts (log per-million). DEGs in bold indicate genes for which the MIA-induced changes are reversed by SAFit2.

| Gene | MIA SAFit2 Early | MIA Vehicle Early | MIA SAFit2 Late | MIA Vehicle Late | MIA | Control | SAFit2 reverses MIA effect |
|---|---|---|---|---|---|---|---|
| 1700016K19Rik | -1.77 ± 0.35 | -0.18 ± 0.67 | -1.33 ± 0.79 | 0.58 ± 0.67 | -1.21 ± 0.37 | 0.17 ± 0.34 | No |
| 1700094D03Rik | 1.40 ± 0.25 | 2.51 ± 0.15 | 1.88 ± 0.19 | 2.27 ± 0.41 | 1.27 ± 0.32 | 2.01 ± 0.17 | No |
| Aimp2 | 4.46 ± 0.11 | 4.56 ± 0.04 | 4.52 ± 0.08 | 4.82 ± 0.05 | 4.31 ± 0.06 | 4.75 ± 0.07 | No |
| Atp5k | 5.63 ± 0.26 | 5.79 ± 0.29 | 6.73 ± 0.10 | 5.71 ± 0.26 | 6.27 ± 0.16 | 5.81 ± 0.17 | No |
| Ccdc180 | -2.65 ± 0.33 | -0.69 ± 0.71 | -1.58 ± 0.68 | -0.23 ± 0.54 | -2.51 ± 0.52 | 0.04 ± 0.26 | No |
| **Ccdc24** | -1.09 ± 0.26 | -0.76 ± 0.32 | -1.84 ± 0.40 | -0.68 ± 0.29 | -0.59 ± 0.32 | -2.05 ± 0.58 | **Yes** |
| Cdhr3 | -0.91 ± 0.21 | 0.23 ± 0.36 | 0.13 ± 0.74 | 0.91 ± 0.51 | -0.25 ± 0.36 | 0.61 ± 0.21 | No |
| Dusp19 | 3.93 ± 0.05 | 3.85 ± 0.14 | 4.48 ± 0.05 | 3.86 ± 0.11 | 4.28 ± 0.17 | 3.80 ± 0.14 | No |
| Dusp6 | 4.23 ± 0.08 | 4.09 ± 0.08 | 4.37 ± 0.08 | 4.00 ± 0.10 | 4.48 ± 0.08 | 4.19 ± 0.03 | No |
| **Efnb1** | **2.28 ± 0.19** | **2.56 ± 0.23** | **1.95 ± 0.25** | **2.52 ± 0.15** | **2.80 ± 0.17** | **2.37 ± 0.10** | **Yes** |
| Fam216b | -2.50 ± 0.40 | 0.96 ± 0.91 | 0.47 ± 1.03 | 1.57 ± 0.47 | 0.35 ± 0.59 | 1.40 ± 0.33 | No |
| Gabpa | 4.48 ± 0.08 | 4.67 ± 0.04 | 4.98 ± 0.12 | 4.25 ± 0.11 | 4.86 ± 0.16 | 4.34 ± 0.09 | No |
| Gas6 | 6.94 ± 0.09 | 6.76 ± 0.11 | 6.45 ± 0.16 | 7.04 ± 0.12 | 6.84 ± 0.12 | 7.17 ± 0.10 | No |
| **Gm3636** | **2.87 ± 0.16** | **2.55 ± 0.15** | **2.74 ± 0.17** | **2.21 ± 0.19** | **1.98 ± 0.24** | **2.54 ± 0.08** | **Yes** |
| Gmfb | 7.84 ± 0.07 | 8.02 ± 0.07 | 8.31 ± 0.14 | 7.74 ± 0.15 | 8.16 ± 0.20 | 7.76 ± 0.09 | No |
| **H2bu1-ps** | **-1.47 ± 0.31** | **-1.87 ± 0.39** | **-0.79 ± 0.43** | **-1.98 ± 0.27** | **-1.31 ± 0.44** | **-0.19 ± 0.29** | **Yes** |
| Hdc | -0.14 ± 0.42 | 1.08 ± 0.30 | 0.41 ± 0.19 | 0.87 ± 0.18 | 0.12 ± 0.24 | 0.85 ± 0.14 | No |
| **Itga7** | **4.22 ± 0.06** | **3.93 ± 0.15** | **3.39 ± 0.19** | **4.14 ± 0.08** | **4.21 ± 0.14** | **3.76 ± 0.19** | **Yes** |
| Lpl | 2.52 ± 0.12 | 3.02 ± 0.21 | 3.41 ± 0.33 | 2.17 ± 0.27 | 2.62 ± 0.23 | 1.97 ± 0.19 | No |
| **Lyz2** | **3.61 ± 0.08** | **4.40 ± 0.27** | **3.14 ± 0.61** | **3.51 ± 0.15** | **4.59 ± 0.25** | **3.96 ± 0.11** | **Yes** |
| Marchf1 | 4.29 ± 0.05 | 4.49 ± 0.09 | 4.89 ± 0.09 | 4.32 ± 0.13 | 4.60 ± 0.12 | 4.24 ± 0.10 | No |
| **Mast1** | **6.59 ± 0.13** | **6.25 ± 0.14** | **5.97 ± 0.16** | **6.73 ± 0.22** | **6.58 ± 0.24** | **7.08 ± 0.13** | **Yes for early** |
| Ngp | -0.58 ± 0.39 | -0.94 ± 0.46 | 0.60 ± 0.52 | -0.97 ± 0.45 | 1.15 ± 0.31 | -1.08 ± 0.67 | No |
| Nrde2 | 3.53 ± 0.11 | 3.72 ± 0.10 | 3.35 ± 0.12 | 3.74 ± 0.12 | 3.38 ± 0.11 | 3.78 ± 0.09 | No |
| Ppp1r32 | -1.22 ± 0.25 | -0.08 ± 0.25 | -1.20 ± 0.70 | -0.07 ± 0.27 | -0.90 ± 0.22 | 0.13 ± 0.26 | No |
| Prag1 | 3.83 ± 0.11 | 3.63 ± 0.16 | 3.22 ± 0.12 | 4.19 ± 0.20 | 4.01 ± 0.21 | 4.52 ± 0.09 | No |
| Ptgfr | 2.33 ± 0.06 | 2.00 ± 0.10 | 2.14 ± 0.21 | 1.75 ± 0.17 | 2.53 ± 0.16 | 1.72 ± 0.21 | No |
| **Renbp** | **2.24 ± 0.05** | **2.45 ± 0.11** | **2.21 ± 0.13** | **2.49 ± 0.09** | **2.49 ± 0.16** | **1.89 ± 0.26** | **Yes** |
| Rims4 | 1.86 ± 0.31 | 1.54 ± 0.31 | 1.12 ± 0.15 | 2.44 ± 0.16 | 2.57 ± 0.19 | 2.06 ± 0.19 | No |
| **Rogdi** | **5.79 ± 0.09** | **5.76 ± 0.07** | **5.21 ± 0.17** | **5.85 ± 0.14** | **5.39 ± 0.18** | **5.89 ± 0.09** | **Yes for late** |
| Shprh | 4.83 ± 0.11 | 4.64 ± 0.10 | 4.91 ± 0.10 | 4.58 ± 0.10 | 4.94 ± 0.11 | 4.63 ± 0.05 | No |
| Slc14a2 | -2.68 ± 0.23 | -1.76 ± 0.31 | -2.07 ± 0.45 | -0.35 ± 0.46 | -2.02 ± 0.33 | -0.54 ± 0.36 | No |
| Slfn5 | 3.82 ± 0.08 | 3.97 ± 0.15 | 4.38 ± 0.12 | 3.73 ± 0.12 | 4.05 ± 0.15 | 3.62 ± 0.05 | No |
| **Smyd1** | **3.46 ± 0.31** | **2.86 ± 0.08** | **3.78 ± 0.26** | **3.48 ± 0.08** | **3.49 ± 0.32** | **4.26 ± 0.17** | **Yes** |
| **Snx29** | **2.51 ± 0.20** | **2.30 ± 0.19** | **1.78 ± 0.27** | **2.96 ± 0.08** | **3.21 ± 0.07** | **2.82 ± 0.17** | **Yes for late** |
| Ubn2 | 4.86 ± 0.11 | 5.10 ± 0.19 | 4.79 ± 0.15 | 3.96 ± 0.15 | 4.59 ± 0.19 | 4.13 ± 0.10 | No |
| Ugt8a | 7.96 ± 0.06 | 8.15 ± 0.12 | 8.64 ± 0.16 | 8.09 ± 0.13 | 8.28 ± 0.14 | 7.96 ± 0.08 | No |
